# Integrating targeted genome mining and structure-guided modeling reveals unexplored 7-deazapurine-containing pathways

**DOI:** 10.64898/2026.04.15.718813

**Authors:** José D. D. Cediel-Becerra, Marc G. Chevrette, Valérie de Crécy-Lagard, Raquel Dias

## Abstract

7-deazapurines are nucleoside analogs that play key roles in nucleic acid modification and can serve as building blocks for diverse, bioactive secondary metabolites. Despite their biological significance, their biosynthetic diversity, distribution, and enzymatic determinants of structural diversification remain poorly understood. Here, we leverage large-scale targeted genome mining, phylogenetic, and network analysis to explore 7-deazapurine-containing pathways across ∼2 million bacterial genomes. We identified over 900 candidate biosynthetic gene clusters (BGCs), grouped into more than 100 families, most of which remain uncharacterized. These GATOR-GC-predicted BGCs were predominantly found in *Streptomyces*. We then examined enzyme-substrate interactions in three representative pathways: (i) peptidyl-deazapurines, (ii) huimycin, and (iii) dapiramicin A. Molecular docking and molecular dynamics (MD) simulations recapitulated known enzyme-substrate interactions and highlighted candidate catalytic residues governing amide bond formation, methylation, and glycosylation. Using this genome- and structure-guided framework, we identified a candidate BGC for dapiramicin A and proposed tailoring steps, including scaffold methylation and deoxy-sugar formation. These findings expand the known diversity of 7-deazapurine-containing BGCs and demonstrate how integrating genome mining with structural modeling can link BGCs to chemical function, providing a foundation for discovering and characterizing 7-deazapurine-containing secondary metabolites.

**Graphical abstract:** 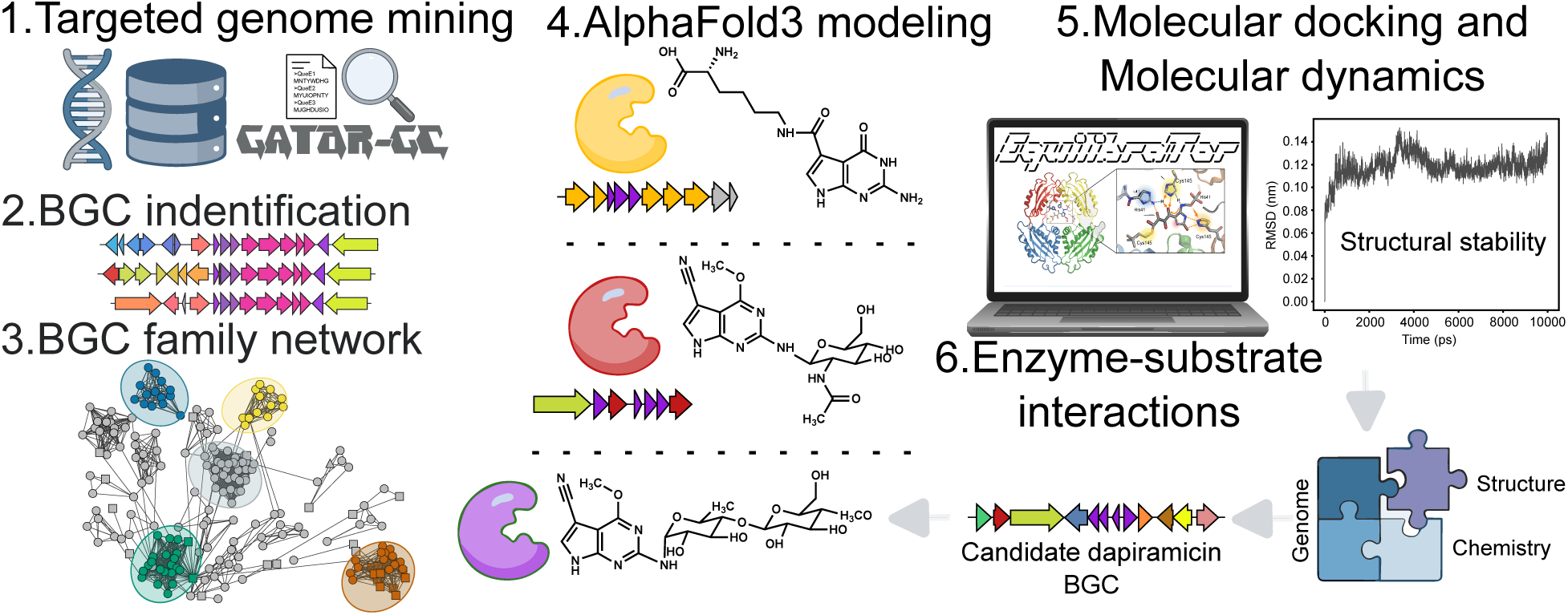

## Introduction

7-deazapurines are nucleoside analogs containing a pyrrolo[2,3-d]pyrimidine core that play critical roles in nucleic acid modification, influencing essential biological processes such as cellular stress resistance, self/non-self discrimination, and the fine-tuning of translation efficiency (1). They also serve as building blocks of structurally diverse secondary metabolites. Notable examples include toyocamycin and tubercidin, which exhibit antibacterial and antifungal activities (2–5). In addition, the deazapurine moiety itself exhibits antitumoral activity (6) and has been exploited in drug discovery as a purine isostere, exemplified by the FDA-approved anticancer agent ribociclib (1). Given their potent and diverse biological activities, 7-deazapurine-containing metabolites may represent a promising yet largely underexplored source of novel therapeutics to address global health challenges, including antimicrobial resistance and cancer.

Genes required in 7-deazapurine assembly are usually clustered in bacterial genomes (7). Their biosynthesis starts with the conversion of guanosine-5’-triphosphate (GTP) to 7,8-dihydroneopterin triphosphate (H_2_NTP) by GTP cyclohydrolase I (FolE/FolE2), followed by the formation of 6-carboxy-5,6,7,8-tetrahydropterin (CPH_4_) via QueD. The radical S-adenosylmethionine (SAM) enzyme QueE catalyzes ring contraction to generate 7-carboxy-7-deazaguanine (CDG), the first core intermediate. Finally, 7-cyano-7-deazaguanine synthase (QueC) converts CDG to 7-cyano-7-deazaguanine (preQ₀) in two sequential ATP-dependent steps, completing assembly of the 7-deazapurine moiety (1, 8). The resulting 7-deazapurine moiety can be incorporated into either DNA or tRNA; enzymes responsible for DNA insertion are often co-localized with these biosynthetic genes, whereas tRNA-inserting transglycosylases are typically encoded elsewhere in the genome (9, 10). In contrast, experimentally characterized pathways for 7-deazapurine-containing secondary metabolites (*i.e.*, toyocamycin/sangivamycin, tubercidin, huimycin, and roseomycin A) are encoded in genomic regions that lack nucleic acid incorporation machinery. These instead contain tailoring enzymes, regulators, or transporters, features consistent with biosynthetic gene clusters (BGCs) (2, 7).

Genome mining tools such as antiSMASH are widely used for BGC discovery, scanning genomes for core biosynthetic enzymes, and applying pathway-specific detection rules based on conserved gene content and organization (11). This rule-based framework has enabled the identification of numerous BGC classes. Nevertheless, 7-deazapurine-containing BGCs have likely remained undetected because dedicated detection rules were only recently implemented in antiSMASH v.8.0.0 (12). Although more than 20 7-deazapurine-containing secondary metabolites have been structurally characterized, only five have experimentally confirmed BGCs, leaving the rest as “orphan” pathways (*i.e*., metabolites whose BGCs are not yet known) (13). Consequently, the biosynthetic diversity, taxonomic distribution, and architectural variation of 7-deazapurine-containing BGCs remain poorly understood, underscoring the need for systematic exploration to map these pathways comprehensively.

Targeted genome mining with GATOR-GC can enable the systematic identification of candidate 7-deazapurine-containing BGCs by searching their core biosynthetic genes across genomes at scale (14, 15). However, linking the presence of tailoring enzymes to the chemistry of the resulting 7-deazapurine-containing metabolite remains challenging. For instance, in peptidyl-deazapurine pathways, divergent QueC homologs such as RoyL have evolved altered substrate specificity, shifting from ammonia to various amino acids (e.g., L-lysine, L-ornithine, L-diaminobutryric acid). These enzymes define a new family of deazapurine amide bond synthetases that attach these and other amino acids to CDG (16). Structural approaches facilitate bridging challenges in biosynthetic prediction: molecular docking can identify active-site residues likely involved in CDG recognition, and subsequent genetic validation via site-directed mutagenesis can confirm their role in L-lysine coupling during roseomycin A biosynthesis (16). While docking can identify catalytically relevant interactions, it provides only static snapshots of enzyme-substrate conformations (17). Molecular dynamics (MD) simulations can complement docking by assessing the stability, conformational flexibility, and persistence of enzyme-substrate interactions over time in solvated, physiologically relevant conditions (18–20). Together, these structure-based approaches offer mechanistic insights that sequence analysis alone cannot provide, improving the prediction of chemical outcomes from genomic data.

Here, we integrate targeted genome mining, phylogenetic and network analysis, AlphaFold3 modeling, molecular docking, and MD simulations to map the biosynthetic diversity of 7-deazapurine-containing secondary metabolites across two million bacterial genomes. Using GATOR-GC, we identified over 900 predicted putative 7-deazapurine-containing BGCs, which clustered into more than 100 families. We then examined enzyme-substrate interactions in three representative pathways: (i) peptidyl-deazapurines as a proof of concept (with known BGCs and enzyme-substrate interactions), (ii) huimycin (with a known BGC but unresolved interactions), and (iii) dapiramicin A (with an unknown BGC and uncharacterized interactions). Our results reveal that only five of the 160 predicted 7-deazapurine-containing families are characterized and demonstrate how an integrated genome-and structure-guided framework can provide mechanistic insights into enzyme-substrate specificity and could support BGC deorphanization.

## Material and Methods

### Databases for targeted genome mining

All genomes from AllTheBacteria release 0.1 (∼1.93M; (21)), all genomes from the Natural Products Discovery Center (NPDC) portal (16K; (22)), all bacterial BGCs (∼2K) from the MIBiG v4.0 database (13), and the Actinomycetota genomes (∼54K) from NCBI (retrieved June 2025), were downloaded in FASTA format. The taxonomy classification for these genomes (∼2M) was performed using classify_wf in the GTDB-Tk v2.3.0 (23). The FASTA files were used to train gene-calling models and predict genes using Prodigal v2.6.3 (24). Feature qualifiers for the predicted genes and amino acid sequences were exported in GenBank format files. The PRE-GATOR-GC module from GATOR-GC V1.0.0, a targeted homology-based genome mining tool for identifying gene clusters (hereafter referred to as GATOR windows), was used to build DIAMOND (25) and the modular domains database (15) from GenBank files.

### Targeted genome mining with GATOR-GC

To identify QueE-containing genomic regions using GATOR-GC, three phylogenetically distant QueE proteins were selected as required queries to mitigate sequence divergence and reduce the likelihood of missing hits in distantly related taxa. These included sequences from the experimentally validated huimycin pathway (locus_tag: KALB_407; protein ID: AHH97435.1; BGC0002354), as well as representative 7-deazapurine pathways from *Bacillus subtilis* (NP_389257.1) and *Vibrio sp* (WP_338417914.1). These protein sequences were combined into a single FASTA file and aligned against the targeted GATOR-GC database using DIAMOND in ultra-sensitive mode with a block size of 20 and a chunk size of 1.0. The resulting DIAMOND output was parsed by GATOR-GC, and GATOR windows were identified using a maximum intergenic distance of 5 kb and a window extension of 10 kb. Unique GATOR windows, representing distinct gene content and genomic architecture, were identified by GATOR-GC using GATOR Focal Scores (GFS), a metric of window-window similarity.

### Protein domain annotation within GATOR windows

To annotate the functional context surrounding QueE hits, Pfam-A models from the InterPro database (retrieved July 2025; (27)) and Hidden Markov Models (HMMs) of Secondary Metabolite-specific Clusters of Orthologous Groups (smCoGs)(12) were used. To identify signature enzymes capable of incorporating 7-deazapurine moieties into nucleic acids, previously curated and validated HMMs were applied (1). These enzymes included archaeal tRNA-guanine transglycosylase (aTGT; EC 2.4.2.48), bacterial Queuine tRNA-ribosyltransferase (bTGT; EC 2.4.2.29) (12), eukaryotic heterodimeric Queuine tRNA-ribosyltransferase (eTGT; EC 2.4.2.64, composed of the catalytic subunit QTRT1 and the accessory subunit QTRT2), and 7-deazapurine DNA insertion protein (DpdA) (12). HMM searches were performed using hmmsearch (HMMER v3.3.2) (26) using an E-value threshold of 1e-5. To account for multi-domain proteins, hits were grouped by HMM profile (domain), and for each profile, only the hit with the highest bit score per protein was retained. In cases where multiple hits overlapped on the protein sequence, the highest-scoring hit in the overlapping region was kept, ensuring that distinct domains were preserved.

### QueE phylogenetic analysis

HMM hits for QueE sequences within unique GATOR windows were aligned using MAFFT v7.515 (28). A phylogeny was constructed in IQ-TREE v2.1.0 (29) using Maximum Likelihood inference, applying the best-fit evolutionary model (*i.e*., LG+G+I), and performing 100 bootstrap replicates. The resulting consensus tree was visualized in iTOL V6.0.0 (30), and annotated with: (i) branch colors indicating phylum-level taxonomy, (ii) presence of 7-deazapurine biosynthetic enzymes, and (iii) presence of signature enzymes that incorporate 7-deazapurines into nucleic acids.

### Identification of full 7-deazapurine-containing pathways

GATOR windows were considered full 7-deazapurine-containing pathways only if they contained HMM hits for all the enzymes responsible for CDG formation (FolE, QueD, and QueE) or preQ_0_ formation (FolE, QueD, QueE, and QueC). Specifically, this was indicated by the presence of the Pfam domains GTP_cyclohydro I (PF01227) or GCHY-1 (PF02649) for FolE, PTPS (PF01242) for QueD, and PF06508 for QueC. In contrast, the detection rule implemented in antiSMASH v8.0.1 requires the presence of enzymes involved in preQ_0_ biosynthesis, which overlooks 7-deazapurines whose moiety derives from CDG.

### Identification of full 7-deazapurine-containing pathways potentially involved in secondary metabolism

To distinguish between nucleic acid modification and secondary metabolism pathways, a genome-mining rule was established to identify 7-deazapurine-containing BGCs. Briefly, full 7-deazapurine-containing pathways were prioritized when dpdA HMM hits were absent within GATOR windows, and when their host genomes lacked HMM hits for tRNA-modifying enzymes (aTGT, or eTGT, or bTGT) (1). The secondary metabolism context of the prioritized pathways was annotated by classifying SmCoGs hits according to antiSMASH categories: additional biosynthetic (tailoring), regulatory, transporter, and other. Criteria for identifying potential 7-deazapurine-containing BGCs were established using experimentally validated 7-deazapurine-containing MIBiG BGCs as a reference. Based on these known pathways, it was determined that a candidate BGC should contain at least one tailoring HMM hit, and/or at least one hit in either the transporter or regulatory category. Full 7-deazapurine-containing pathways meeting these criteria were considered potential 7-deazapurine-containing BGCs, and their genomic neighborhoods were generated using clinker v0.0.32 (31).

### Clustering and network construction of 7-deazapurine-containing pathways

GFS values were used to construct a similarity matrix for prioritized full 7-deazapurine-containing pathways (GATOR windows) not associated with nucleic acid modification, from which pairwise Euclidean distances were calculated. Hierarchical clustering with complete linkage was then performed on the resulting distance matrix. To define groups of similar GATOR windows, a range of distance cutoffs was assessed, and an initial cutoff of 4 was selected using the elbow method. Clustering was validated by confirming that experimentally validated 7-deazapurine MIBiG BGCs formed distinct groups whose members retained the signature tailoring enzymes required for 7-deazapurine modification and production of the corresponding BGC product. Groups containing a 7-deazapurine MIBiG BGC were defined as BGC families, representing groups of GATOR windows similar to reference BGCs, whereas groups lacking a MIBiG representative were classified as unassigned. Both families and unassigned groups were used to construct a network. To improve visualization clarity and avoid an overly cluttered network, only the strongest inter-group connections (GFS ≥ 0.9) were retained. A node table was compiled with GTDB-Tk taxonomy information and subfamily/unassigned group labels.

For comparison, BiG-SCAPE 2.0 was employed to cluster the same windows across multiple cutoffs (0.1, 0.3, 0.5, 0.75, 1) using the default “mix” weighting score, a composite metric applied consistently across all BGC classes without per-class tuning, in global alignment mode (32). A BiG-SCAPE cutoff of 0.75 was used to define gene cluster families (GCFs) based on 7-deazapurine MIBiG BGCs, ensuring non-singleton groups and retention of signature tailoring enzymes, consistent with the GATOR-GC-based network. Networks were visualized in Cystoscope v3.10.4 using the default layout (“preferred”) (33).

### Molecular docking and molecular dynamics simulations

Protein-ligand complexes were prepared and simulated using a standardized workflow. Briefly, ligand structures, including cofactors and substrates (e.g., 7-deazapurine derivatives), were obtained from PubChem (34) or generated from SMILES strings using RDKit v2024.09.2 (35). Hydrogens were added, and the geometries were optimized before exporting the ligands as PDB files. Protein sequences were extracted either from MIBiG v.4.0.0 BGC entries or from GATOR-GC-predicted 7-deazapurine-containing regions and modeled using the AlphaFold 3 server (36). Top-ranked models were selected based on AlphaFold 3 internal confidence ranking (i.e., highest mean predicted Local Distance Difference Test, pLDDT score). The selected structural models were then converted to PDB format with OpenBabel v3.1.1 (37). Potential binding pockets were detected with the CASTpFold server (38) and filtered based on pocket volume and proximity to conserved motifs (Supplementary Figure 1).

Ligands and proteins were prepared for molecular docking using Meeko v0.6.1 (https://pypi.org/project/meeko/). Before docking, ligand geometries were subjected to a short energy minimization of 1,000 steps using the MMFF94 force field in OpenBabel. Molecular docking was performed with AutoDock Vina v.1.2.7 using the Vina forcefield, an exhaustiveness of 18, and a fixed random seed (12345) (39). The search space was defined based on the predicted binding pockets and, when applicable, the known cofactor-binding site, with a 5 Å expansion to accommodate ligand flexibility. For each ligand, the lowest energy-binding pose was extracted using Meeko.

Docked ligands were split into individual PDB files using RDKit and OpenBabel. Protein and ligand PDB files were provided as inputs to EquilibraTor v1.0.0 (18), which was run with default parameters for molecular dynamics (MD) simulations. All simulations were performed on the University of Florida’s high-performance computing cluster (HiPerGator) B200 node, equipped with an AMD EPYC 9655 P 96-core processor, 756 GB of RAM, and NVIDIA B200Tensor Core GPUs (CUDA Compute). Following production MD, solvent ions were removed, and the resulting protein-ligand complexes were visualized in PyMOL v2.1.2 (40) and LigPlot + v.2.3.0 (41) to highlight ligand protein-ligand interactions. Predicted binding affinities (pkD) for each enzyme-substrate complex were calculated using GLM-score v0.15.2 (42).

### Use Case Analysis

For the peptidyl-deazapurine pathways, structural models of deazapurine-bond synthetases from *Streptomyces rosesporus* (SSIG_07344, NZ_ABYX02000001.1) and *Streptomyces rapamycinicus* NRRL 5491 ( D3C57_103080, CP006567) were used as the enzymes. Adenylated CDG (AMP-CDG), L-lysine, and D-alanyl-D-alanine were used as the ligands and docked into the largest predicted binding pocket of each enzyme (Supplementary Figure 1A-B).

For the huimycin biosynthetic pathway (BGC0002354), structural models of the S-adenosyl methionine (SAM)-dependent methyltransferase (KALB_4069) and the glycosyltransferase (KALB_4073) were used to model the biosynthetic steps. For the first committed step, SAM and preQ_0_ were used as the ligands, and the docking search space was constrained to encompass the conserved SAM-binding glycine-rich motif (43). For the second committed step, uridine diphosphate N-acetylglucosamine (UDP-GlcNAc) and 2-amino-6-methoxy-7-cyano-7-deazapurine were used as ligands (Supplementary Figure 1C-D).

For the dapiramicin A pathway, the structural model of HuiC-like enzyme was used together with SAM and preQ_0_ as ligands for the methylation step, with the docking search space constrained to the conserved SAM-binding glycine-rich motif (43). For the sugar modification steps, structural models of RmlB-like and RmlD-like enzymes were generated using the AlphaFold server in complex with their respective cofactors, NAD^+^, and NADP^+^. The substrates dTDP-D-glucose and dTDP-4-keto-6-deoxyglucose were then docked into the largest predicted binding pocket of each enzyme. (Supplementary Figure 1E-G).

## Results

### 7-deazapurine-containing pathways are widespread across bacteria and occur in both nucleic acid modification and secondary metabolism contexts

To assess the taxonomic distribution of 7-deazapurine-containing pathways across ∼2 million publicly available genomes, we first ran GATOR-GC with three phylogenetically distant QueE protein queries (see methods). QueE catalyzes a central step in 7-deazapurine biosynthesis, allowing detection of pathways that produce either CDG or preQ_0_ moieties (1). Additional enzymes such as FolE, QueD, and QueC were not required to avoid missing distant hits; these were later identified with Pfam profiles in downstream analyses. In total, we detected 33,450 QueE-containing GATOR windows, of which 3,152 were unique according to GFS. These unique QueE hits spanned 22 phyla and 311 genera (Supplementary Table 1).

The top 15 genera with the highest number of QueE-containing regions spanned Actinomycetota (n = 1,554; 0.7% of Actinomycetota genomes), Campylobacterota (n = 451; 0.4%), and Pseudomonadota (n = 201; <0.1%). Within Actinomycetota, QueE hits were most frequent in *Streptomyces* (1,137 QueE hits; 5.2% of *Streptomyces* genomes), *Kitasospora* (n = 125; 13.6%), *Micromonospora* (n = 101; 6.4%), and *Amycolatopsis* (n = 59; %13.7), with smaller contributions from *Mycobacterium* (n = 49; <0.1%), *Nocardiopsis* (n = 44; 22.3%), and *Nonomuraea* (n = 39; 8%). In Campylobacterota, QueE hits were largely found in *Campylobacter_D* (n = 365; 0.3% of *Campylobacter_D* genomes), *Campylobacter* (n = 45; 5.7%), and *Helicobacter* (n = 41; 2%). In Pseudomonadota, *Vibrio* (n = 49; 0.2% of *Vibrio* genomes), *Brucella* (n = 43; 1.7%), *Pseudomonas_E* (n = 40; 0.8%), *Sinorhizobium* (n = 39; 5.2%), and *Novosphingobium* (n = 30; 27.8%) collectively accounted for most hits (Figure 1A). Overall, QueE-containing regions were taxonomically widespread, though their prevalence was generally low in most taxa and could appear high in taxa represented by few genomes.

**Figure 1.**
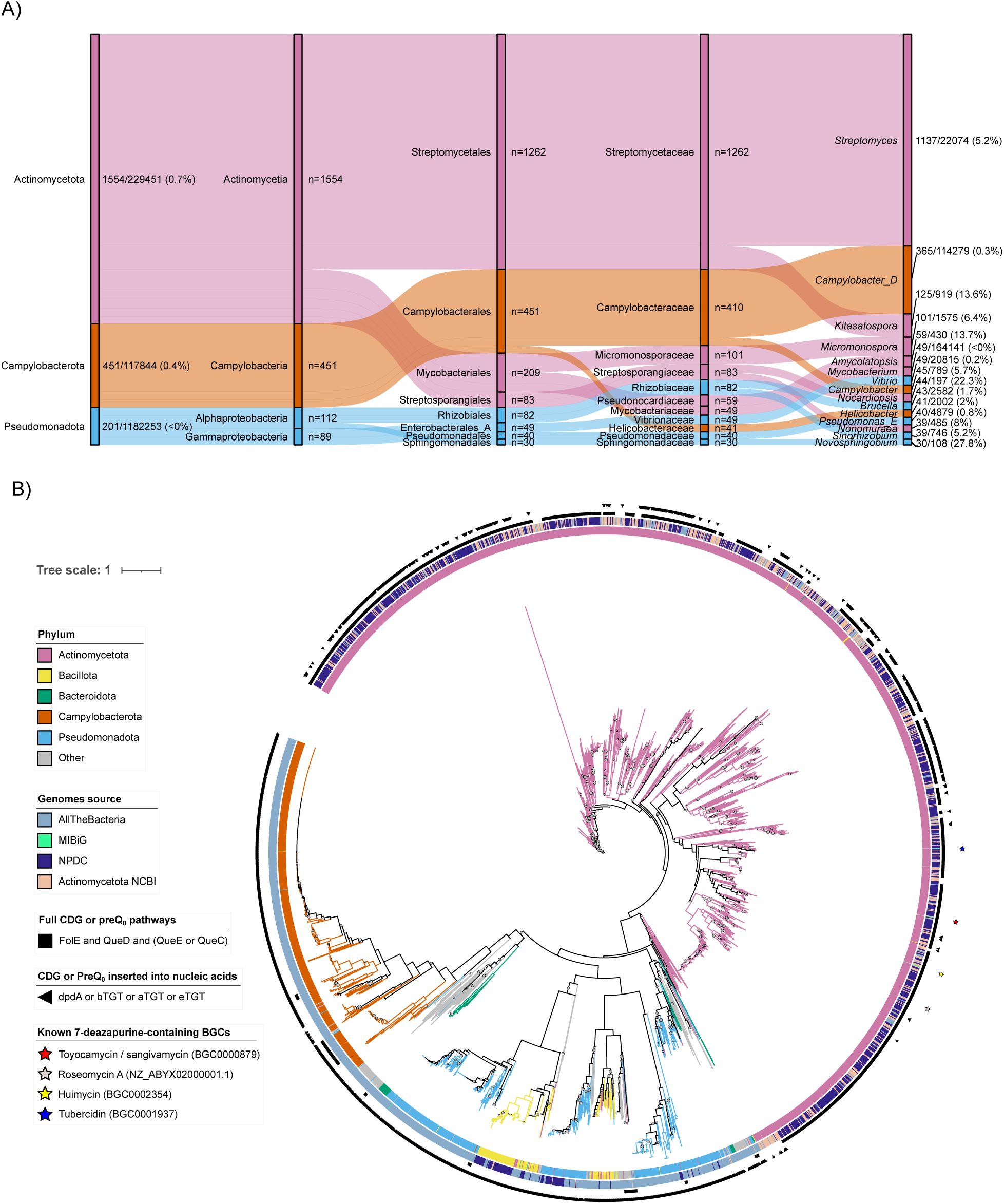
Taxonomic distribution and phylogeny of QueE hits across bacteria. A) Sankey plot illustrating the top 15 genera ranked by the absolute number of QueE-containing regions. Labels indicate both the counts and their proportion of total genomes within each taxon. Ribbon widths reflect absolute counts, with flows representing hierarchical taxonomic assignment from phylum to genus based on GTDB-Tk classification. B) Maximum-likelihood phylogeny of QueE hits, with branches colored by phylum. Full 7-deazapurine-containing pathways (encoding CDG or preQ_0_) are marked with squares, while enzymes that incorporate 7-deazapurine moieties into nucleic acids are marked with triangles. The presence of dpdA is indicated within QueE-containing regions, whereas tRNA-modifying enzymes are indicated at the genome level. Known 7-deazapurine-containing BGCs are shown with stair symbols. Bootstrap values below 70% are displayed.

Next, we evaluated the evolutionary relationships among QueE HMM hits within the 3,152 unique GATOR windows. QueE sequences from Actinomycetota and Campylobacterota formed distinct phylogenetic clades, whereas hits from Bacillota, Bacteroidota, and Pseudomonadota were more broadly distributed across the phylogeny. HMM hits for 7-deazapurine biosynthetic enzymes (FolE and QueD and [QueE or QueC]) were predominantly found in Actinomycetota; QueE-containing regions meeting this criterion are referred to hereafter as full 7-deazapurine-containing pathways. HMM hits responsible for incorporating CDG or preQ_0_ into nucleic acids (dpdA or bTGT or aTGT or eTGT) were broadly distributed across multiple phyla. Notably, MIBiG BGCs encoding experimentally validated 7-deazapurine-containing secondary metabolites contained the full set of HHM hits characteristic of full 7-deazapurine-containing pathways but lacked nucleic acid-incorporation HMM hits (Figure 1B). Based on these observations, we hypothesize that full 7-deazapurine-containing pathways lacking dpdA and tRNA incorporation enzymes elsewhere in the genome may serve alternative roles, including secondary metabolism. These findings informed a detection rule to exclude pathways involved in nucleic acid modification, prioritizing the remaining pathways for downstream analyses.

Applying this detection rule to the full set of 7-deazapurine-containing pathways (n = 1,521), we excluded 555 pathways associated with nucleic acid modification and retained 966 pathways with distinct genomic contexts. To evaluate whether these distinct pathways are associated with secondary metabolism, we annotated their protein-encoding genes using smCoG HMM profiles (see Methods). We defined a secondary metabolism context (i.e., BGC) based on antiSMASH functional categories (i.e., tailoring enzymes, transporters, and regulators), with minimum criteria established from their co-occurrence in experimentally validated 7-deazapurine-containing BGCs (see Methods). Using this framework, we identified 933 putative 7-deazapurine-containing BGCs, distributed as follows: 264 contained at least a tailoring enzyme, regulator, and transporter; 327 contained at least a tailoring enzyme and regulator; 98 contained at least a tailoring enzyme hit; 244 contained only tailoring enzyme hits; and 16 contained none of these smCoG hits (Figure 2A). Among the identified BGCs, we found the experimentally validated pathways encoding toyocamycin/sangivamycin, tubercidin, huimycin, and roseomycin A (Figure 2B), all within *Streptomyces* except huimycin, which is found in *Kutzneria*.

**Figure 2.**
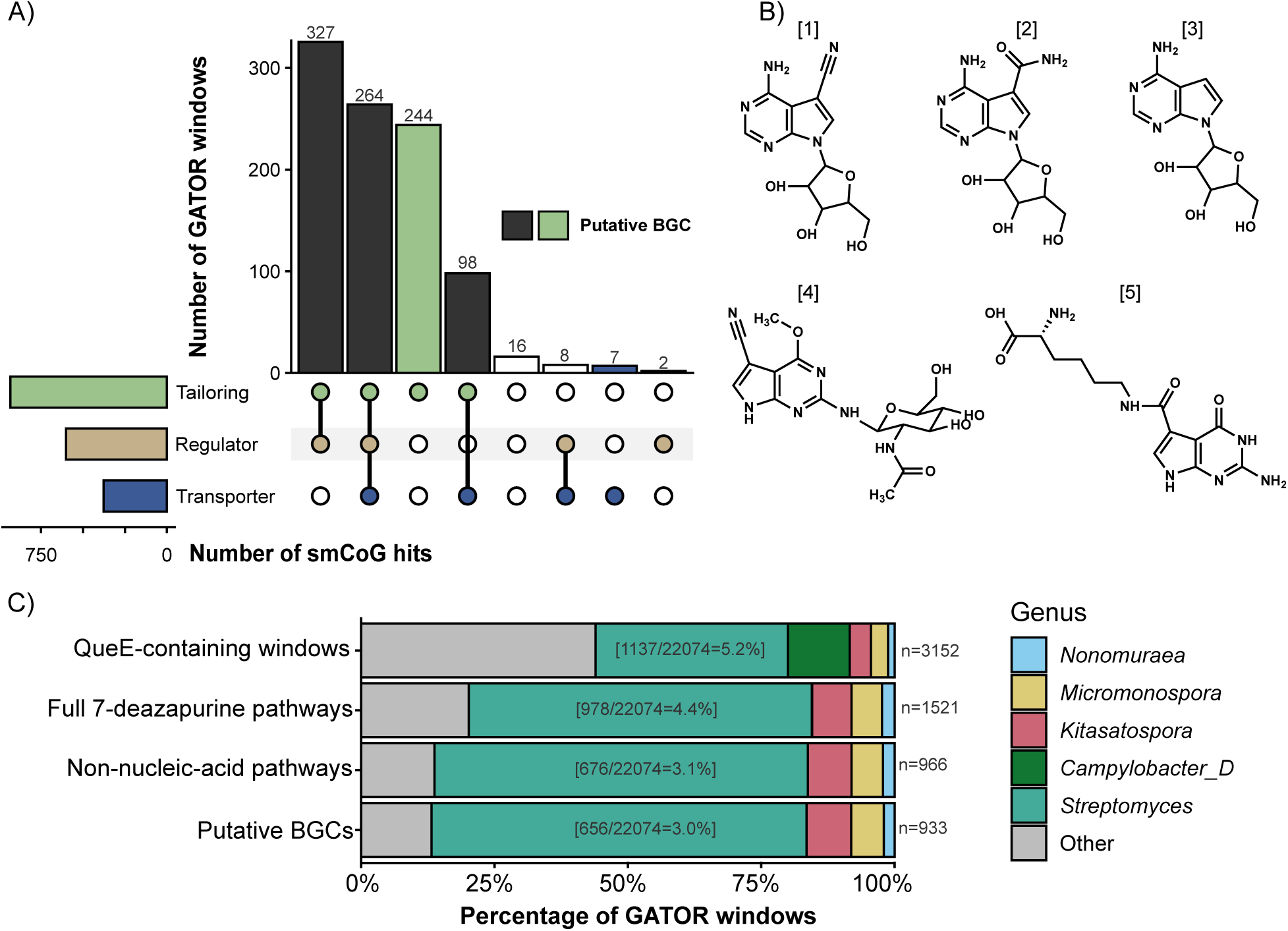
Genomic, chemical, and taxonomic landscape of GATOR-GC-predicted 7-deazapurine-containing pathways. A) Upset plot illustrating the distribution and overlap of smCoG hits across the tailoring, regulator, and transporter categories. Horizontal bars indicate the total number of smCoG hits in each category, while vertical bars (intersection size) represent the number of GATOR windows sharing the specific combination of categories indicated by the connected dots below. B) Chemical diversity of experimentally validated 7-deazapurine metabolites, including toyocamycin [1] and sangivamycin [2] from *S. rimosus*, tubercidin [3] from *S. tubercidicus*, huimycin [4] from *K. albida*, and roseomycin A [5] from *S. roseosporus*. C) Proportion of GATOR windows retained at each step of the filtering pipeline, categorized by genus-level taxonomy with the total number of windows (n) shown on the right of each stacked bar. The contribution of *Streptomyces* is highlighted within the stack, with the label indicating its fraction relative to the total number of genomes sampled.

To examine taxonomic shifts during the detection pipeline, we tracked the relative contribution of genera as GATOR windows progressed from initial QueE-containing regions (n = 3,152) to prioritized putative BGCs (n = 933) (Figure 2C). *Streptomyces* was the dominant genus at every stage, showing a marked increase in proportion as filtering criteria became more stringent. While the number of GATOR windows decreased across the detection pipeline, the relative proportion of *Streptomyces* nearly doubled, rising from ∼35% to ∼70% of the final putative BGCs. The most significant shift in genera composition occurred when requiring the presence of the set of HMM hits for the full 7-deazapurine pathways. This stage effectively filtered out all windows associated with *Campylobacter_D*, suggesting that while QueE is present in this taxon, it likely lacks the full biosynthetic machinery required for CDG/preQ_0_ formation. Conversely, the “Other” taxonomic group shrank by more than half, while additional genera such as *Kitasatospora*, *Micromonospora*, and *Nonomuraea* remained relatively stable. However, when considering the total number of genomes sampled for each genus, *Streptomyces* was not the most enriched genus in terms of putative BGCs. *Kitasatospora* had the highest proportion (78 putative BGCs out of 919 genomes; 8.5%), followed by *Micromonospora* (57/1,575; 3.6%) and *Nonomuraea* (21/485; 4.3%). These findings highlight that, although *Streptomyces* dominates numerically, less sampled genera can have a higher relative frequency of 7-deazapurine-containing BGCs when accounting for the number of genomes surveyed.

### The biosynthetic diversity of putative 7-deazapurine-containing BGCs remains largely unexplored

To compare and group full 7-deazapurine-containing pathways not associated with nucleic acid modification (n = 966) based on shared gene content and architecture, we used both GATOR-GC GFS similarity metric and BiG-SCAPE 2.0. GATOR-GC formed 160 groups of similar 7-deazapurine-containing pathways, including 47 singletons. Only five of these groups were classified as BGC families due to the presence of an MIBiG 7-deazapurine BGC, while the remaining groups remained unassigned (Figure 3). BiG-SCAPE grouped the same pathways into 136 GCFs, which also formed discrete subnetworks that were not exclusively composed of single GCFs, as multiple GCFs were often intermixed within the same subnetwork. However, GCFs corresponding to MIBiG 7-deazapurine BGCs were clearly separated from one another, with their members clustering more tightly (Supplementary Figure 2).

**Figure 3.**
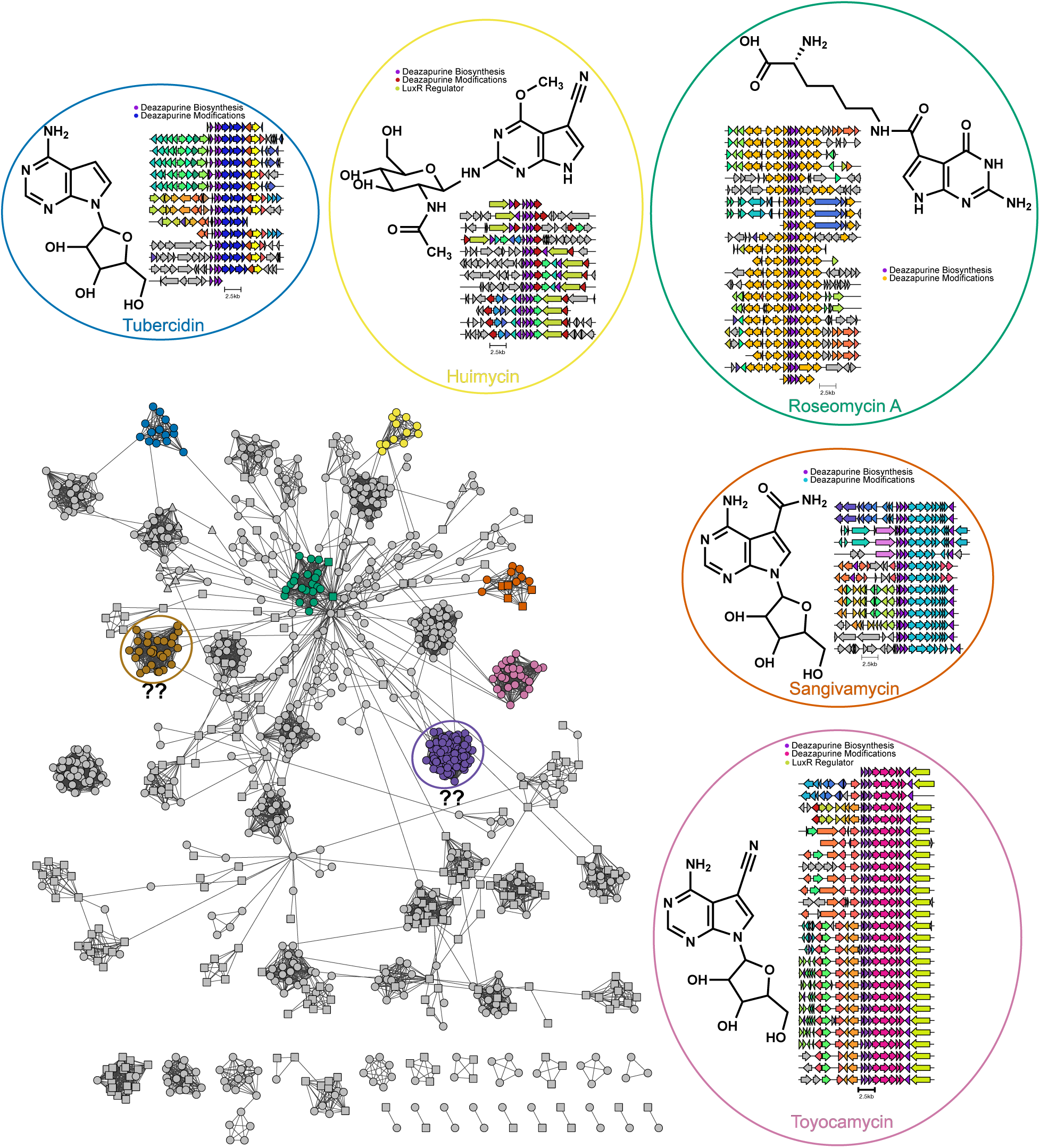
Landscape of 7-deazapurine biosynthetic diversity. Network of full 7-deazapurine-containing pathways not associated with nucleic acid modification (n = 966), identified by GATOR-GC. Nodes represent GATOR windows, and edges indicate window-window similarity based on GFS. Node shapes denote putative BGC classification based on smCoG hits: circles represent windows containing at least one tailoring enzyme hit together with either a transporter or regulatory hit; squares represent windows containing only tailoring enzyme hits; and triangles represent windows lacking these smCoG hits. Singleton groups were excluded to improve visualization. 7-deazapurine-containing BGC families are highlighted along with their genomic neighborhoods. The majority of predicted groups remain unknown, and their genomic contexts are likely associated with secondary metabolism. For example, two uncharacterized groups (circles with question marks) encode distinct tailoring enzymes, regulators, or transporters; their genomic neighborhoods are shown in Supplementary Figures 8 and 9.

GATOR-GC network analyses revealed that all BGC families and most unassigned groups contained members with genomic contexts associated with secondary metabolism. Most groups contained members with at least one tailoring enzyme hit together with either a transporter or regulatory hit (Nodes as circles in Figure 3), or at least one tailoring enzyme hit alone (Nodes as squares in Figure 3). In contrast, a smaller subset of nodes (n = 16) lacked these hits and grouped partially together (Nodes as triangles in Figure 3), suggesting a potentially distinct functional role.

The GATOR-GC toyocamycin family comprised 36 *Streptomyces* putative BGC members, while BiG-SCAPE grouped 10 related BGCs. These BGCs encoded a LuxR-family regulator along with the conserved enzymes required for preQ_0_ scaffold formation. In addition, they harbored the signature tailoring enzymes responsible for converting the preQ_0_ scaffold into toyocamycin, including GMP reductases (PF00478), adenylosuccinate lyases (PF00206), adenylosuccinate synthetases (PF00709), Phosphoribosyl-pyrophosphate transferases (PF00156), and haloacid dehalogenases (PF00702) (Supplementary Figure 3).

In some *Streptomyces* strains, such as *S. rimosus* ATCC 14673 (MIBiG ID BGC0000879), toyocamycin is further converted to sangivamycin by a nitrile hydratase system (α, PF02979; γ, PF02211; β, PF21006), which catalyzes the hydration of the nitrile group of toyocamycin to form an amide (7). In contrast, other toyocamycin BGCs, such as those from *S. diastatochromogenes* 1628 (MIBiG ID BGC0001808) (44) and *S. griseochromogenes* ATCC 14511 (MIBiG ID BGC0000881)(45), lack this nitrile hydratase system, resulting in toyocamycin as the final product.

GATOR-GC clustering identified sangivamycin-associated BGCs as a distinct family closely related to toyocamycin producers. This group included 13 *Streptomyces* members, whereas BiG-SCAPE clustered five related BGCs. These members retained the conserved toyocamycin biosynthetic core but were distinguished by the presence of a complete nitrile hydratase system, enabling sangivamycin-specific modifications (Supplementary Figure 4).

The tubercidin family included 13 *Streptomyces* BGCs identified by GATOR-GC and eight members clustered by BiG-SCAPE. Unlike the toyocamycin and sangivamycin pathways, these pathways lack QueC and instead use CDG as the building block. They encoded a LuxR-family regulator and a distinct set of tailoring enzymes, including phosphoribosyltransferase (PF00156), a UbiD-family decarboxylase (PF01977), GMP reductase (PF00478), and a haloacid dehalogenase (PF00293) that modify CDG to yield tubercidin (Supplementary Figure 5).

The huimycin family comprised 12 BGCs according to GATOR-GC, originating from *Streptomyces* (n=5), *Micromonospora* (n=4), *Kutzneria* (n=2), and *Actinosynnema* (n=1). BiG-SCAPE grouped five related BGCs, including four from *Streptomyces* and one from *Kutzneria*. These BGCs retained preQ_0_ as the biosynthetic core, with a LuxR-family regulator, and decorating enzymes including glycosyltransferases (PF13439) and SAM-dependent methyltransferases (PF08241) (Supplementary Figure 6).

The roseomycin A family included 22 BGCs (GATOR-GC), mostly from *Streptomyces* (n = 19) and *Kitasospora* (n = 3), with four BGCs in BiG-SCAPE. These diverged from other families: although they encode QueC hits, these enzymes do not catalyze the formation of preQ₀. Instead, they catalyze the formation of the amide bond between CDG and L-lysine, generating roseomycin A. Additional tailoring enzymes include ATP-grasp enzymes (PF13535), TauD dioxygenases (PF02668), and fatty acid desaturases (PF00487) (Supplementary Figure 7).

Interestingly, none of the experimentally characterized families corresponded to the largest unassigned families. Among these, one large uncharacterized group comprised 81 putative BGCs across *Streptomyces* (n = 77), *Micromonospora* (n = 1), *Amycolatopsis* (n = 2), and *Actinoplanes* (n = 1) (Purple in Figure 3). These retained the preQ₀ biosynthetic core and encoded non-heme dioxygenases (PF14226), Asp/Asn β-hydroxylases (PF05118), and, in some BGCs, SAM-dependent methyltransferases (PF13649). All members also encoded transcriptional regulators with bacterial activator domains and tetratricopeptide repeats (PF03704) (Supplementary Figure 8).

A second uncharacterized group comprised 27 putative BGCs, mainly from *Streptomyces* and one from *Kitasatospora* (Brown in Figure 3). In addition to the preQ₀ core, these clusters encoded non-heme dioxygenases (PF14226), aminotransferases (PF00202), FAA hydrolases (PF01557), and SAM-dependent methyltransferases (PF13649). This combination of enzymes suggests production of decorated deazapurine analogs, potentially expanding chemical diversity beyond known 7-deazapurine-containing metabolites (Supplementary Figure 9).

Collectively, these findings indicate that currently known chemical diversity of 7-deazapurine-containing secondary metabolites represents only a fraction of the predicted biosynthetic diversity encoded by bacterial genomes. In the following sections, we complement these genomic predictions with structure-guided simulations, including AlphaFold structural modeling, molecular docking, and MD simulations, to gain mechanistic insights into the biosynthesis of representative experimentally validated and predicted 7-deazapurine-containing BGCs.

### Structure-guided simulations recapitulate and predict catalytic interactions in peptidyl 7-deazapurine biosynthesis

To explore how AlphaFold structural modeling, molecular docking, and MD simulations can generate mechanistic hypotheses linking biosynthetic enzymes to 7-deazapurine chemistry, we used the roseomycin A pathway from *Streptomyces rosesoporus* (16) as a proof of concept. Peptidyl 7-deazapurine BGCs encode RoyL, a novel amide-bond-forming enzyme that attaches diverse amino acids to adenylated CDG (1) (AMP-(1)), producing distinct metabolites. Catalytic residues interacting with the ε-amino group of L-lysine, such as Tyr41, His45, and Glu48, have been experimentally validated for roseomycin production (16).

We tested whether our structure-guided framework could recapitulate these interactions. The roseomycin A BGC was identified in our curated database (Green in Figure 3), and modeling RoyL in complex with AMP-(1) and L-lysine showed that the substrate adopted a catalytically competent pose, with its ε-amino group positioned 2.9 Å from AMP-(1). The catalytic residues His45 and Glu48 formed hydrogen bonds with L-lysine, while Tyr41 established a stabilizing hydrophobic contact within the amine-binding pocket. Binding of AMP-(1) was dominated by hydrophobic interactions, yielding a predicted binding affinity of pkD = 6.06. Key residues contributing to AMP-(1) recognition included Arg104, Asn105, Asp132, and Cys139 (Supplementary Figure 10). These results confirm that our framework can reproduce experimentally observed interactions and substrate positioning.

We next applied the approach to a distinct peptidyl 7-deazapurine BCG from *Streptomyces rapamycinicus* NRRL 5491 (Sr-deaz BGC). In this BGC, a RoyL homolog (AGP59494.1) catalyzes the formation of the amide bond between the dipeptide D-alanyl-D-alanine and AMP-(1), producing a novel metabolite (2). Residue-level interactions and BGC context remained unknown (16). The Sr-deaz BGC is located between the actinoplanic acid and rapamycin BGCs (Figure 4A), whose metabolic products target key eukaryotic pathways, including Ras farnesyltransferase and the mTOR signaling pathway, respectively (46, 47). The Sr-deaz BGC encodes two *Streptomyces* antibiotic regulatory protein (SARP) regulators, an ATP-grasp protein, a SAM-dependent methyltransferase, and a β-hydroxylase (Figure 4B).

**Figure 4.**
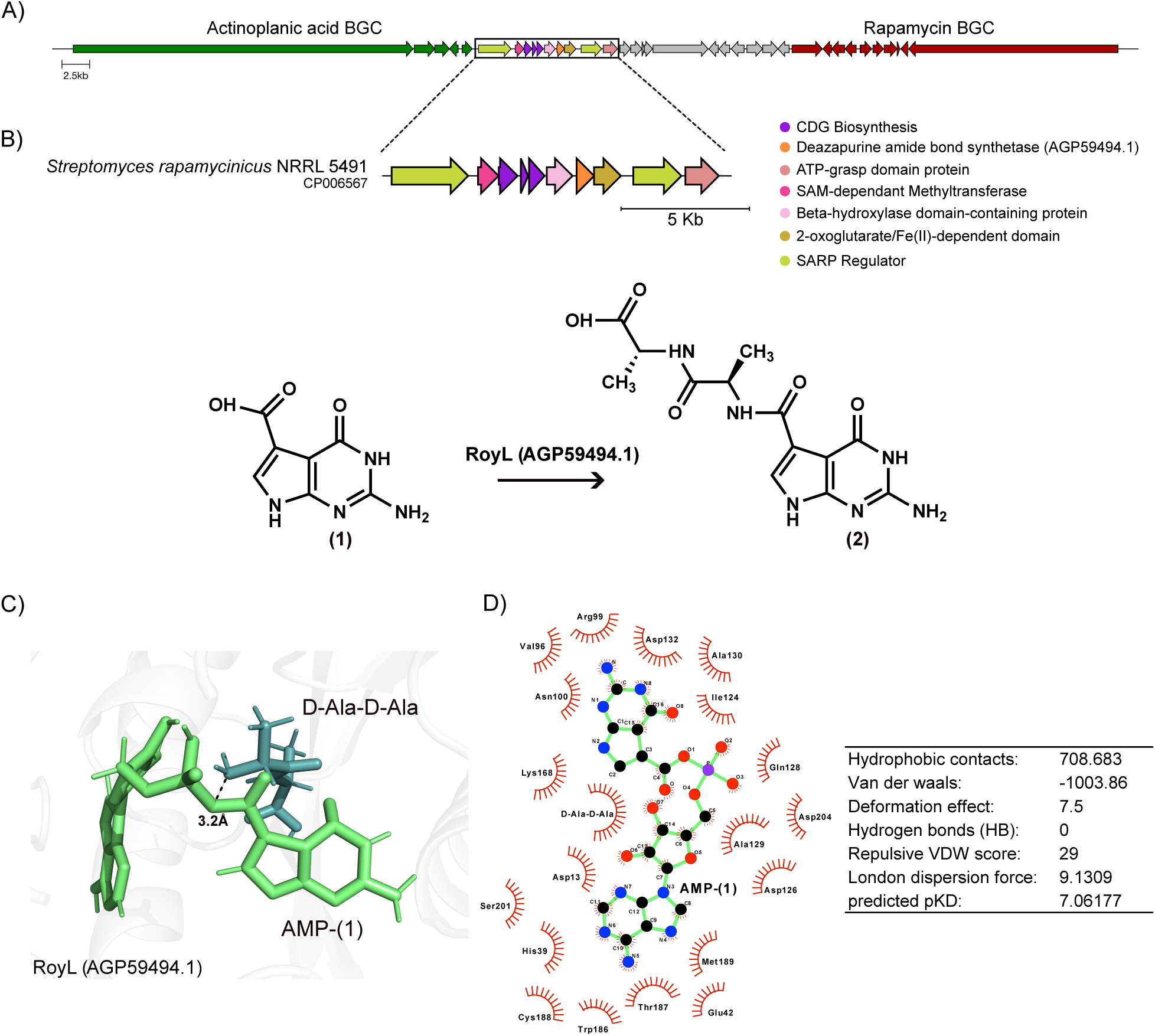
Enzyme-substrate interactions in the predicted Sr-deaz BGC. A) Genomic “supercluster” encoding the actinoplanic acid, candidate Sr-deaz, and rapamycin BGCs. B) Candidate peptidyl 7-deazapurine BGC with predicted enzyme annotations, and the biosynthetic step catalyzed by the RoyL homolog, which forms the amide leading to the peptidyl 7-deazapurine metabolite. C) Representative snapshot from production MD simulations of the RoyL homolog in complex with AMP-(1) and D-alanyl-D-alanine, highlighting a potential pre-reactive amide-bond-forming state. D) Key hydrophobic and hydrogen-bond interactions stabilizing substrate binding within the active site, along with the predicted binding affinity for AMP-(1).

To investigate substrate recognition, we applied our structure-guided approach. We modeled AGP59494.1 in complex with AMP-(1) and D-alanyl-D-alanine and performed production MD simulations. The substrate adopted a catalytically competent pose, with its amino positioned 3.2 Å from the carbonyl group of AMP-(1) (Figure 4C). The ternary complex remained structurally stable throughout the simulation (Supplementary Figure 11). Substrate recognition was dominated by hydrophobic interactions (Leu103, Arg99, Ser9, Gly122, and Ile121) and a hydrogen bond with Asn100, yielding a predicted binding affinity of pKD = 3.6. AMP-(1) interacted with an expanded binding pocket involving 19 residues (e.g., His39, Asn100, Ala129, Ala130, Asp132, and Lys168), resulting in a predicted binding affinity of pKD = 7 (Figure 4D). These results suggest that AGP59494.1 can accommodate D-alanyl-D-alanine in a pre-reactive geometry, providing a mechanistic explanation for its observed substrate specificity and supporting its role in forming metabolite (2).

Together, these results demonstrate that our structure-guided docking and MD framework can recapitulate known catalytic interactions and predict unresolved enzyme-substrate interactions. We next applied this approach to systems in which both enzymes and substrates are known, but residue-level interactions remain unresolved.

### In silico identification of potential key residues mediating huimycin biosynthesis

Although the huimycin biosynthetic pathway from *Kutzneria albida* DSM 43870 has been characterized (48), the residue-level determinants governing substrate recognition and catalysis for the SAM-dependent methyltransferase HuiC and the glycosyltransferase HuiG remain unknown. To address this gap, we constructed structural models of HuiC and HuiG and analyzed their enzyme-substrate complexes using our structure-guided approach. In this pathway, HuiC catalyzes O-methylation of preQ_0_ (1) using SAM as the methyl donor, generating the intermediate 2-amino-6-methoxy-7-cyano-7-deazapurine (2), which is subsequently glycosylated by HuiG using UDP-GlcNAc as the sugar donor, forming huimycin (3) (Figure 5A).

**Figure 5.**
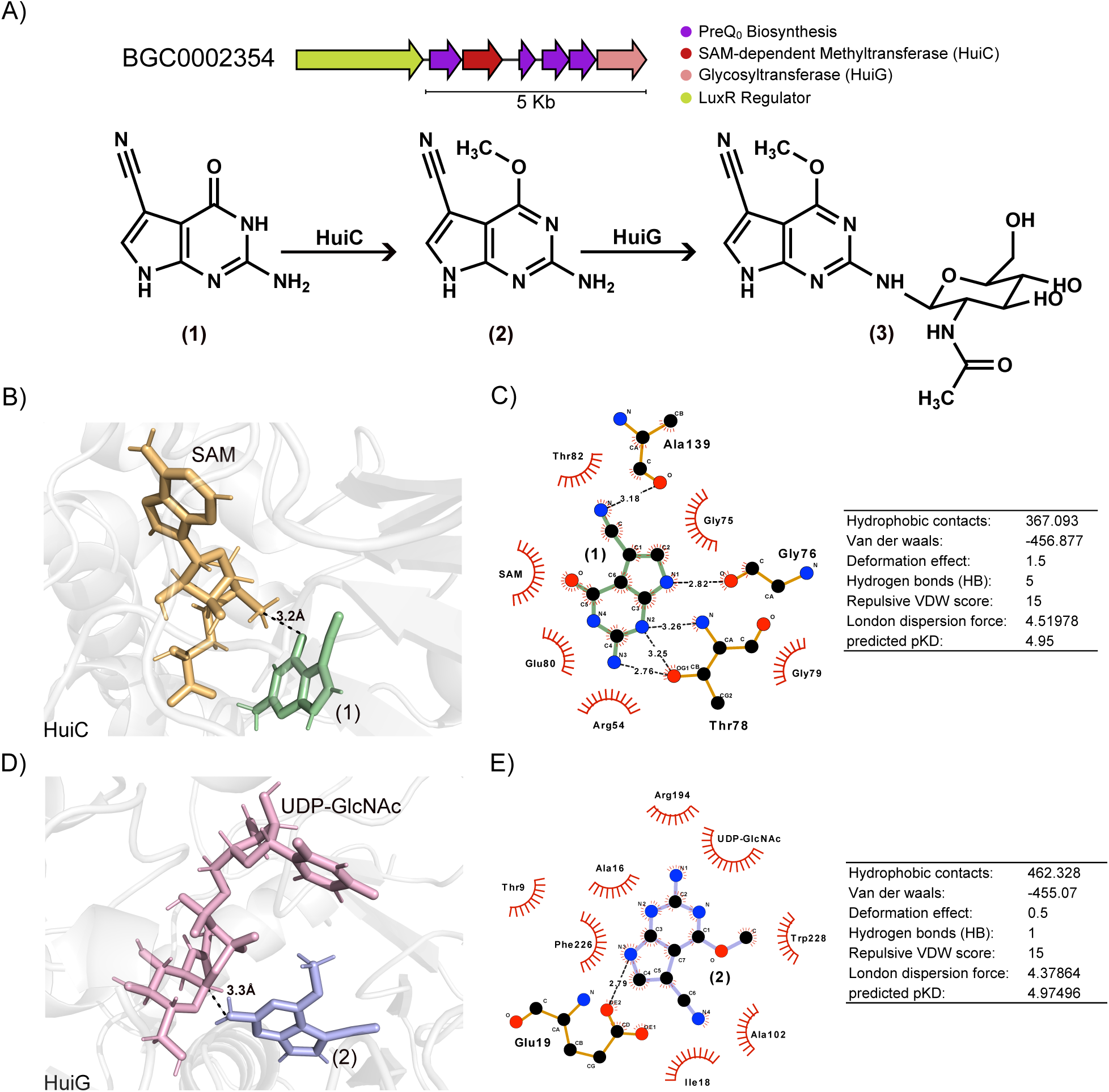
Enzyme-substrate interactions in the huimycin pathway. (A) Huimycin BGC showing predicted enzyme annotations and known biosynthetic steps. B) Representative snapshot from production MD simulations for the methyltransferase HuiC in complex with substrate (1) and SAM, showing the distance between reactive groups consistent with a pre-reactive methyl transfer state, and C) the key hydrophobic and hydrogen-bond interactions stabilizing substrate binding, along with the predicted binding affinity. D) Representative snapshot from production MD simulations of the glycosyltransferase HuiG in complex with substrate (2) and UDP-GlcNAc, highlighting the distance between reactive groups consistent with a pre-reactive glycosyl transfer state, and E) the corresponding hydrophobic and hydrogen-bond interactions stabilizing substrate binding, together with the predicted binding affinity.

Production MD simulations revealed that SAM and (1) adopted a catalytically competent conformation within the HuiC active site, with the SAM methyl carbon positioned 3.2 Å from the phenolic oxygen of (1) (Figure 5B). Substrate stabilization was mediated by five hydrophobic contacts involving residues Arg54, Gly75, Thr82, Gly79, and Glu80, as well as five hydrogen bonds formed with the backbone nitrogen of Gly79, and the OG1 atom of Thr78 (two interactions) (Figure 5C) Binding affinity estimates supported these observations, with moderate binding affinity for (1) (pKD = 4.95) and stronger affinity for SAM (pKD = 7.3). SAM binding was stabilized by hydrophobic interactions with Gly75 and Gly77, residues within the conserved glycine-rich motif characteristic of SAM-dependent methyltransferases, and by a hydrogen bond between the N3 atom of SAM and the OE1 atom of Glu124. The HuiC–SAM–(1) complex remained structurally stable throughout the production simulation, as indicated by the RMSD convergence and low structural fluctuations (Supplementary Figure 12).

We next examined the glycosylation step catalyzed by HuiG. Production MD simulations showed that UDP-GlcNAc and intermediate (2) adopted a reactive configuration in which the carbonyl group of the N-acetyl glucosamine moiety of UDP-GlcNAc was positioned 3.3 Å from the nucleophilic nitrogen of (2) (Figure 5D). This complex was stabilized by seven hydrophobic contacts (Phe226, Thr9, Ala16, Arg194, Trp228, Ala102, and Ile18), and a hydrogen bond between N3 of (2) and the OE2 of Glu19. Consistent with these interactions, predicted binding affinities indicated moderate binding of (2) (pKD = 4.97), and stronger binding for UDP-GlcNAc (pKD = 6.6). Low RMSD fluctuations confirmed that the HuiG–UDP-GlcNAc–(2) complex remained stable throughout the production simulation (Supplementary Figure 13).

Collectively, these simulations define a coherent catalytic framework for huimycin biosynthesis, identifying catalytically competent substrate poses and highlighting residues likely essential for methylation and glycosylation. Residues such as Gly76, Thr78, and Glu124 in HuiC, and Glu19 in HuiG, emerge as strong candidates for future experimental validation.

### Structure-guided inference identifies candidate enzymes underlying dapiramicin A biosynthesis

Having validated our structure-guided framework on experimentally characterized BGCs, we applied it to dapiramicin A, a 7-deazapurine-containing metabolite with an unknown BGC. Discovered in 1983 from *Micromonospora* sp SF-1917 (whose genome has not been sequenced), dapiramicin A exhibits potent antifungal activity against *Rhizoctonia solani,* and is structurally similar to huimycin, but with a different sugar scaffold: 4’-O-methyl-ß-ᴅ-glucopyranosyl-6’-deoxy-α-ᴅ-glucopyranosyl disaccharide (Figure 6A) (49, 50). Based on its structure, we hypothesize that the BGC should minimally encode (i) a SAM-dependent methyltransferase for O-methylation of preQ_0_ (1) (HuiC-like) to yield 2-amino-6-methoxy-7-cyano-7-deazapurine (2), as in huimycin; (ii) a methyltransferase for 4’-O-methylation of ß-ᴅ-glucopyranosyl unit; (iii) enzymes for 6’-deoxy-α-ᴅ-glucopyranosyl formation (RmlB and RmlD from the rhamnose-like pathway); and (iv) one or more glycosyltransferases for disaccharide attachment (Figure 6B).

**Figure 6.**
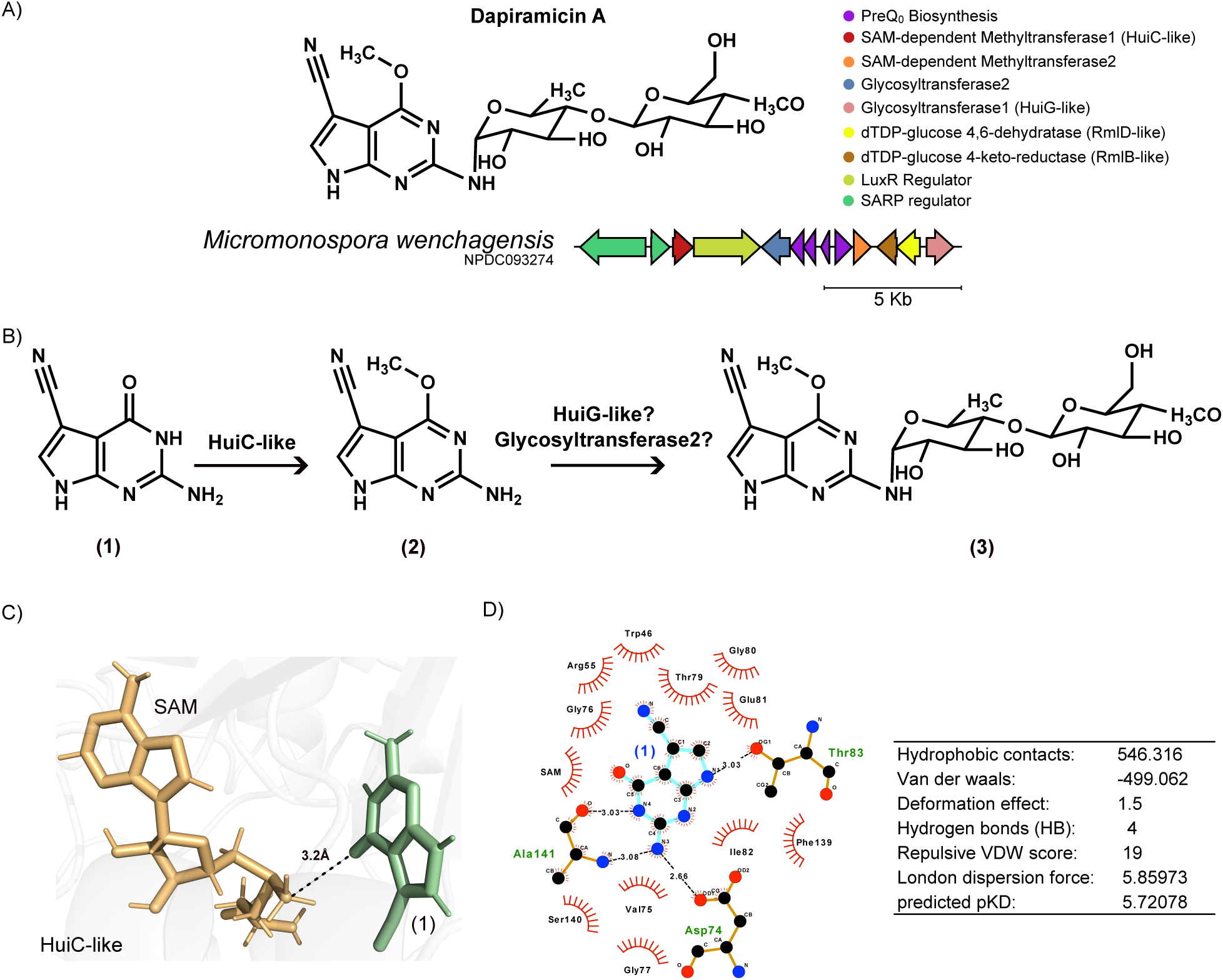
Genomic discovery of a putative BGC encoding dapiramicin A. (A) Candidate BGC showing dapiramicin A chemical structure and predicted enzyme annotations. B) Known and unknown biosynthetic steps. C) Representative snapshot from production MD simulations for the methyltransferase HuiC-lke in complex with substrate (1) and SAM, showing the distance between reactive groups consistent with a pre-reactive methyl transfer state, and D) the key hydrophobic and hydrogen-bond interactions stabilizing substrate binding, along with the predicted binding affinity.

To identify putative BGCs responsible for dapiramycin A production, we searched unassigned families for members containing all expected tailoring enzymes. This revealed a family of nine candidate BCGs distributed across diverse clades [*Micromonospora*, *Actinosynnema*, and *Streptomyces* (Supplementary Figure 14)]. We selected *Micromonospora wenchagensis* (NPDC093274) as a representative BGC for this family for downstream molecular docking and MD simulations. In addition to all expected signature enzymes, this BGC contains two LuxR regulators and one SARP regulator (Figure 6A). We found that the structural model of HuiC-like enzyme was highly similar to HuiC, with an RMSD of 0. 4 Å (Supplementary Figure 15). The structural models of RmlB and RmlC were also similar to the crystal structures of dTDP-D-glucose 4,6-dehydratase (PDB=1G1A, RMSD = 0.9 Å) (Supplementary Figure 16), and dTDP-4-dehydrorhamnose reductase (PDB=3SC6, RMSD = 0. Å (Supplementary Figure 17).

Previous studies suggested that methylation of (1) precedes glycosyltransfer in dapiramicin A biosynthesis (51, 52). To evaluate whether HuiC-like can catalyze this step, we performed production MD simulations of the HuiC-like–SAM–(1) ternary complex as a proxy for the formation of intermediate (2). Throughout the simulation, the carbonyl oxygen of (1) remained positioned 3.2 Å from the SAM methyl group, a geometry compatible with methyl transfer (Figure 6B). Substrate (1) was stabilized by four hydrogen bonds with Ala141, Asp74, and Thr83, along with 10 hydrophobic contacts involving Gly76, Thr79, and Glu81 (Figure 6C). Together, these interactions supported stable substrate engagement, reflected in a predicted binding affinity of pKD = 5.72 (Figure 6D). SAM was similarly well accommodated within the active site, forming two hydrogen bonds with Trp250 and Ala125 and 13 hydrophobic interactions, including Arg102, Tyr146, Val143, and Val147, resulting in a predicted binding affinity of pKD = 7.12. The ternary complex remained structurally and energetically stable throughout the trajectory, supporting the feasibility of SAM-dependent methylation of substrate (1) by the HuiC-like enzyme (Supplementary Figure 18).

Given that RmlB and RmlD enzymes are known to act only on sugar-activated substrates, this suggests that the dapiramicin A sugar moiety is modified before its attachment to (2). RmlB dehydrates the nucleotide sugar dTDP-D-glucose to dTDP-4-keto-6-deoxy-D-glucose (53) (Supplementary Figure 19A). To simulate this RmlB-catalyzed step, we ran production MD simulations for the RmlB-like-NAD^+^-dTDP-D-glucose complex and found that dTDP-D-glucose (specifically, the glucose moiety) was positioned 2.8 Å from the nicotinamide ring of NAD^+^ (Supplementary Figure 19B). The substrate dTDP-D-glucose formed nine hydrophobic contacts and nine hydrogen bonds, including the well-known catalytically important residues Glu126, Tyr148, Val86, and Ser84 that mediate catalysis (53). The substrate showed a predicted binding affinity of pKD = 5.94 (Supplementary Figure 19C). Overall, this complex was structurally stable during the MD production simulation (Supplementary Figure 20).

The presence of an RmlD-like enzyme in the predicted dapiramicin A BGC may suggest that the resultant modified sugar is a rhamnose derivative (54, 55); however, given that the BGC lacks an RmlC-like enzyme, which catalyzes the step required to yield 4-keto-rhamnose, we hypothesized that the RmlD-like enzyme can accept dTDP-4-keto-6-deoxy-D-glucose as a substrate and reduce it to dTDP-6-deoxy-D-glucose (Supplementary Figure 19A). To evaluate whether the RmlD-like enzyme could be responsible for this reduction, we performed production MD simulations for the RmlD-like-NADP-dTDP-4-keto-6-deoxy-D-glucose complex and analyzed its enzyme-substrate interactions.

Our production MD analysis showed that substrate dTDP-4-keto-6-deoxy-D-glucose was well-positioned in the active site of the RmlD-like enzyme, interacting with NADP (Supplementary Figure 19D), and forming seven hydrogen bonds and 10 hydrophobic contacts. The residues Arg180, Asp102, and Arg214 form hydrogen bonds with the glucose moiety, anchoring it within the active site. The predicted substrate binding affinity for the substrate was pKD=5.94 (Supplementary Figure 19E). In comparison, the predicted binding affinity for the canonical substrate, dTDP-4-keto-rhamnose, was lower (pKD=4.35), primarily due to fewer hydrogen bonding interactions in the active site (Supplementary Figure 19F-G). Both complexes remained structurally stable throughout the MD simulations, supporting the possibility that the RmlD-like enzyme could accept the dTDP-4-keto-6-deoxygenase as a substrate (Supplementary Figure 21, Supplementary Figure 22).

Overall, these structure-guided analyses highlight a candidate BGC, suggest plausible enzyme-substrate interactions, and outline potential biosynthetic steps for dapiramicin A in *Micromonospora wenchagensis*, pending experimental validation.

## Discussion

Our systematic survey of 7-deazapurine-containing pathways across ∼2 million bacterial genomes provides the first comprehensive exploration of this biosynthetic family, revealing both the evolutionary conservation and remarkable biosynthetic diversification. Full preQ_0_- and CDG-producing systems are typically associated with tRNA and DNA modification, where they contribute to translation fidelity and genome protection against cellular defense mechanisms (1, 10). Consistent with previous reports of their broad distribution (1), our phylogenetic analyses show that CDG or preQ_0_ biosynthetic genes often occur independently of nucleic acid incorporation enzymes. This decoupling suggests that 7-deazapurine scaffolds can serve functions beyond nucleic acid modifications, including secondary metabolite biosynthesis. Guided by this insight, we refined the 7-deazapurine detection rules (12), demonstrating that genomes encoding full pathways but lacking nucleic acid incorporation enzymes and instead harboring tailoring enzymes, regulators, and/or transporters associated with secondary metabolism can identify 7-deazapurine-containing BGCs.

We identified more than nine hundred predicted 7-deazapurine-containing BGCs spanning diverse genera within Actinomycetota, including the MIBiG BGCs for toyocamycin (3, 7), sangivamycin (7), tubercidin (56), huimycin (48), and roseomycin A (16). However, the majority of predicted BGCs remain uncharacterized. Network analyses using GATOR-GC GFS and BiG-SCAPE 2.0 resolved numerous large groups composed primarily of uncharacterized BGCs that retain conserved preQ_0_ or CDG core genes but differ in their tailoring repertoires, potentially driving the structural diversification of 7-deazapurine-containing metabolites. For instance, members of the huimycin family encode glycosyltransferases and methyltransferases, whereas the roseomycin A family contains amide-bond synthetase homologs (16, 48). Beyond these characterized BGCs, many additional uncharacterized members in other groups may harbor novel enzyme families yet to be discovered. Notably, the largest uncharacterized groups encode conserved non-heme dioxygenases and β-hydroxylases with unknown functions, highlighting clear targets for biochemical investigation.

To interrogate enzymatic function, we applied molecular docking and MD simulations to representative pathways. These analyses reproduced experimentally supported enzyme-substrate interactions and provided a mechanistic rationale for substrate specificity. For example, the amide synthetases RoyL (roseomycin A) and RoyL-like (AGP59494.1, Sr-deaz BGC) positioned L-lysine and D-alanyl-D-alanine, respectively, in catalytically competent poses consistent with amide bond formation (57). Similarly, MD simulations of huimycin tailoring enzymes indicate that the SAM-dependent methyltransferase HuiC and glycosyltransferase HuiG stabilize substrates through a defined hydrogen bonding and hydrophobic interactions, consistent with established features of SAM-dependent methyltransferases, such as the conserved glycine-rich motifs (43, 58).

We then extended this structure-guided framework to dapiramicin A, proposing a candidate BGC in *Micromonospora wenchagensis*. Production MD simulations support a model in which a HuiC-like enzyme catalyzes O-methylation of preQ_0_, while RmlB- and RmlD-like enzymes generate the 6’-deoxy sugar intermediates, and glycosyltransferases could assemble the disaccharide onto the 7-deazapurine scaffold. Our MD simulations analyses suggest that the RmlD-like enzyme may accept dTDP-4-keto-6-deoxy-D-glucose as a substrate despite the absence of an RmlC homolog, implying potential flexibility in sugar biosynthesis consistent with the known plasticity of sugar-modifying enzymes (59). Although experimental validation is required, the integrated genomic and structural evidence supports a coherent biosynthetic model. Experimental characterization of predicted methyltransferase, sugar biosynthesis enzymes, and glycosyltransferases will be necessary to confirm substrate specificities and reaction order. These studies will test whether the substrate flexibility inferred from MD simulations translates into catalytic promiscuity in vitro, as observed for RoyL (16).

Despite these insights, structure-guided predictions have inherent limitations. AlphaFold models provide a single conformational state (e.g., open or closed), which can influence ligand accessibility and binding poses, and low-confidence regions further reduce reliability (60, 61). Protein binding pockets can be predicted using conserved motifs and CASTpFold-identified cavities (38); when such features are unclear or absent, blind docking across the full protein surface may be necessary, although this often positions ligands non-physiologically. Binding affinity estimates derived from GLM scoring, trained primarily on small-molecule inhibitors (42), may not fully capture enzyme-substrate interactions, although they remain useful for relative comparisons. MD simulations provide local sampling around the starting structure and may not capture large conformational changes or long-timescale allosteric effects relevant for catalysis. Nevertheless, these approaches offer valuable insights into substrate positioning, interaction networks, and plausible catalytic conformations. Combining with emerging AI/ML methods, integrated structure-guided modeling may also reveal long-range interactions and allosteric communication beyond the timescales of conventional MD simulations (62).

This work expands the recognized biosynthetic diversity of 7-deazapurine-containing pathways beyond previously characterized metabolites. The widespread occurrence of 7-deazapurine-containing BGCs across diverse genera within Actinomycetota, together with variable tailoring enzymes across 7-deazapurine families, suggests that substantial chemical diversity remains to be explored and that several known metabolites, such as dapiramicin A, may still lack assigned BGCs. By integrating large-scale targeted genome mining, phylogenetic and network analyses, and structure-guided inference, we provide a framework to prioritize uncharacterized BGCs and generate mechanistic hypotheses linking specific enzymes to CDG/preQ_0_ moieties. These hypotheses can be experimentally tested to confirm enzyme-substrate relationships and to connect enzyme function to metabolite chemistry, thereby accelerating the discovery and functional annotation of 7-deazapurine-containing secondary metabolites.

## Acknowledgements

We thank Steven Bruner and Yousong Ding for their constructive comments and suggestions.

## Author contributions

J.D.D.C.-B, conceptualization, investigation, data curation, methodology, supervision, software, data curation, formal analysis, visualization, project administration, writing–original draft, writing–review & editing. M.G.C, methodology, writing–review & editing. V.d.C.-L, supervision, conceptualization, data curation, methodology, writing—review & editing. R.D, conceptualization, supervision, methodology, formal analysis, visualization, funding acquisition, resources, project administration, writing–review & editing.

## Supplementary data

## Data availability

The data underlying this article are available in *[7-deazapurine manuscript data],* at 10.5281/zenodo.19600739

---

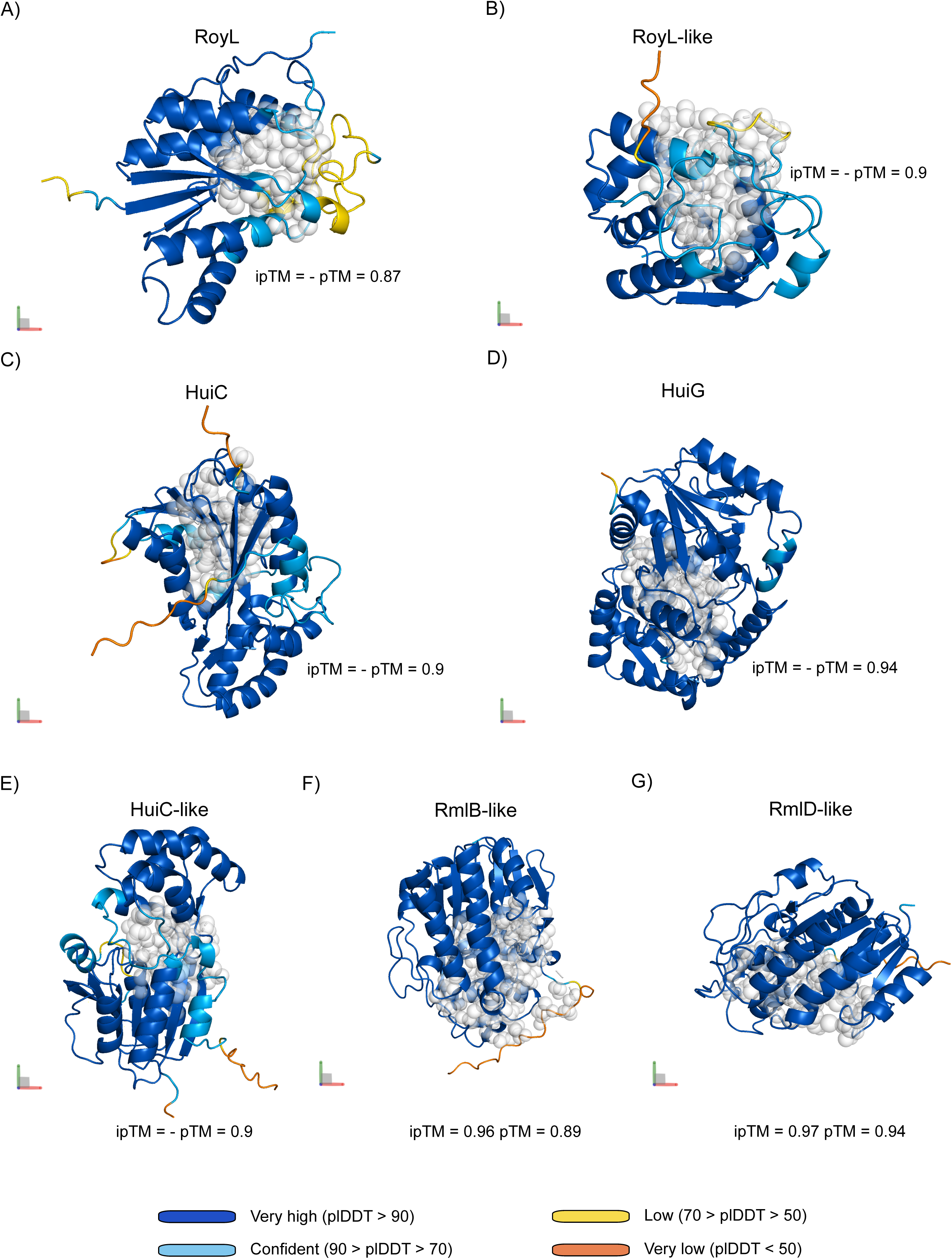

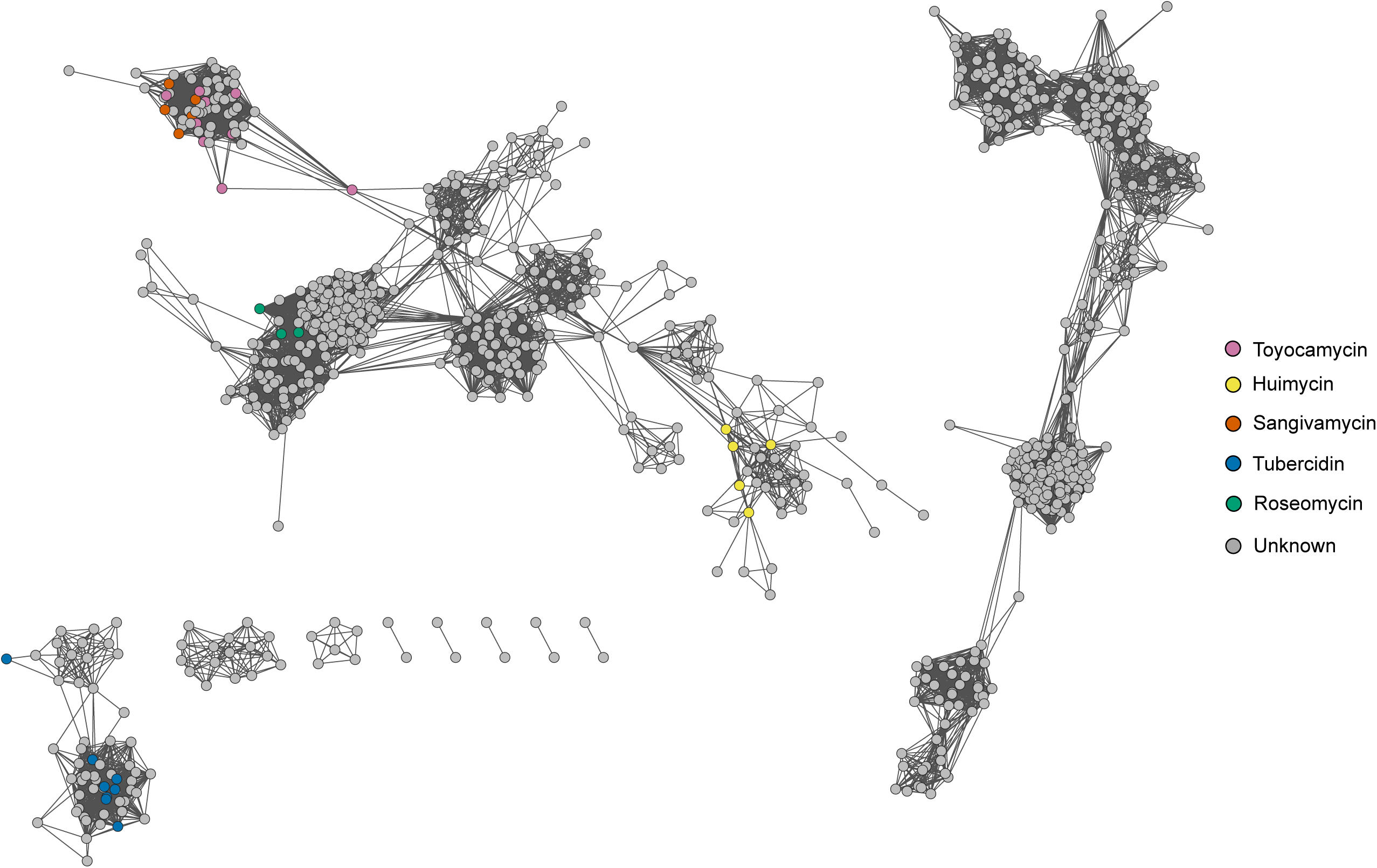

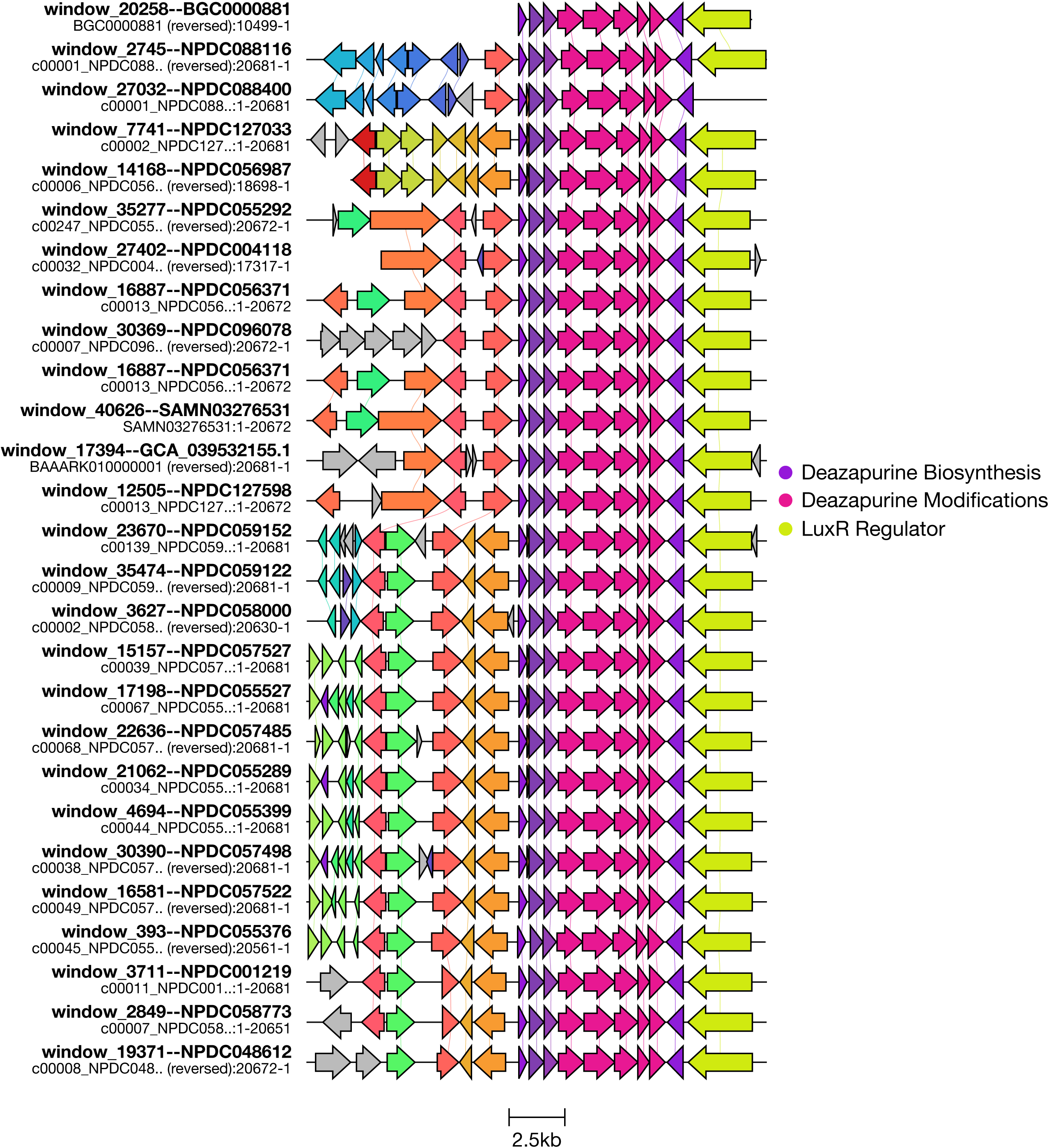

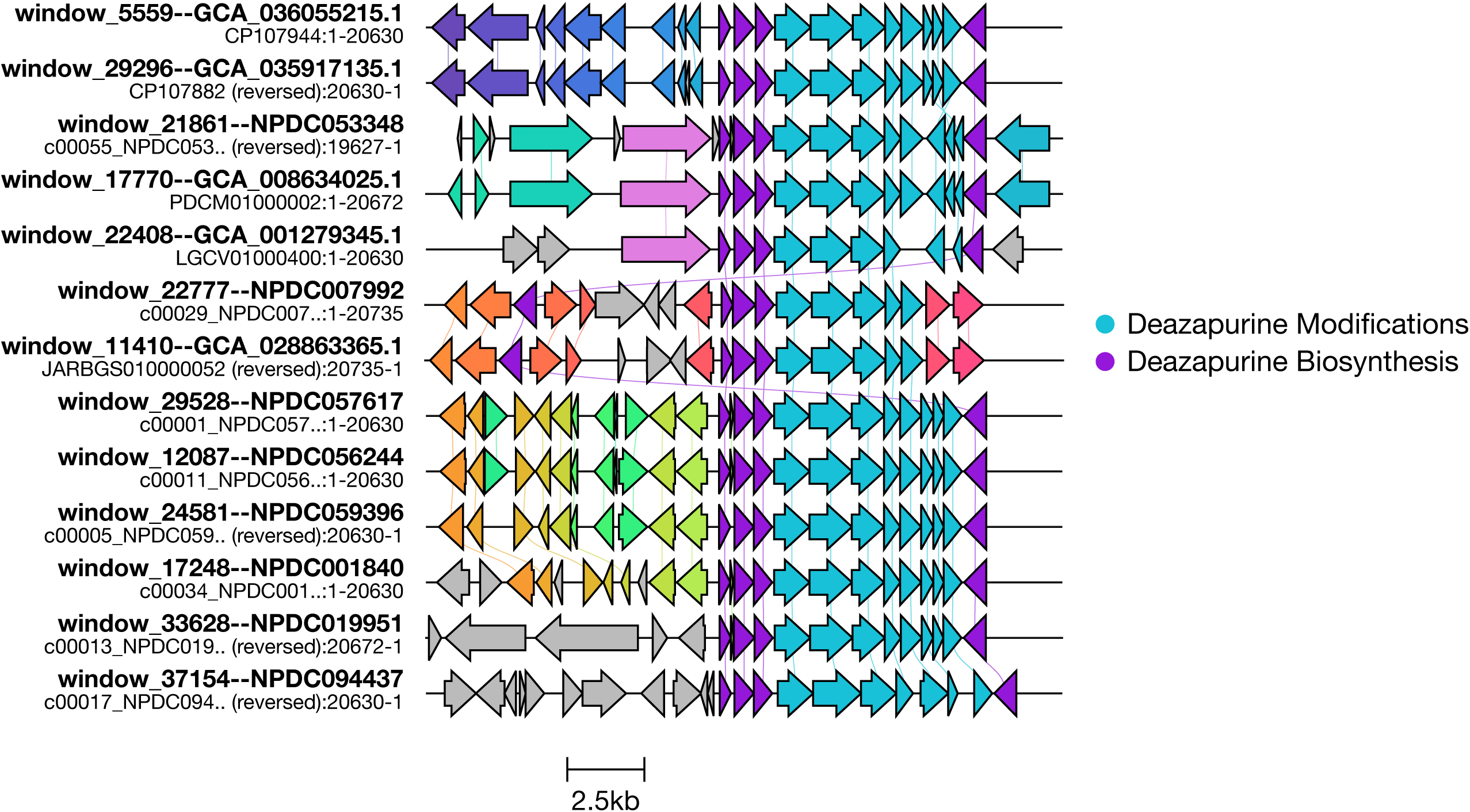

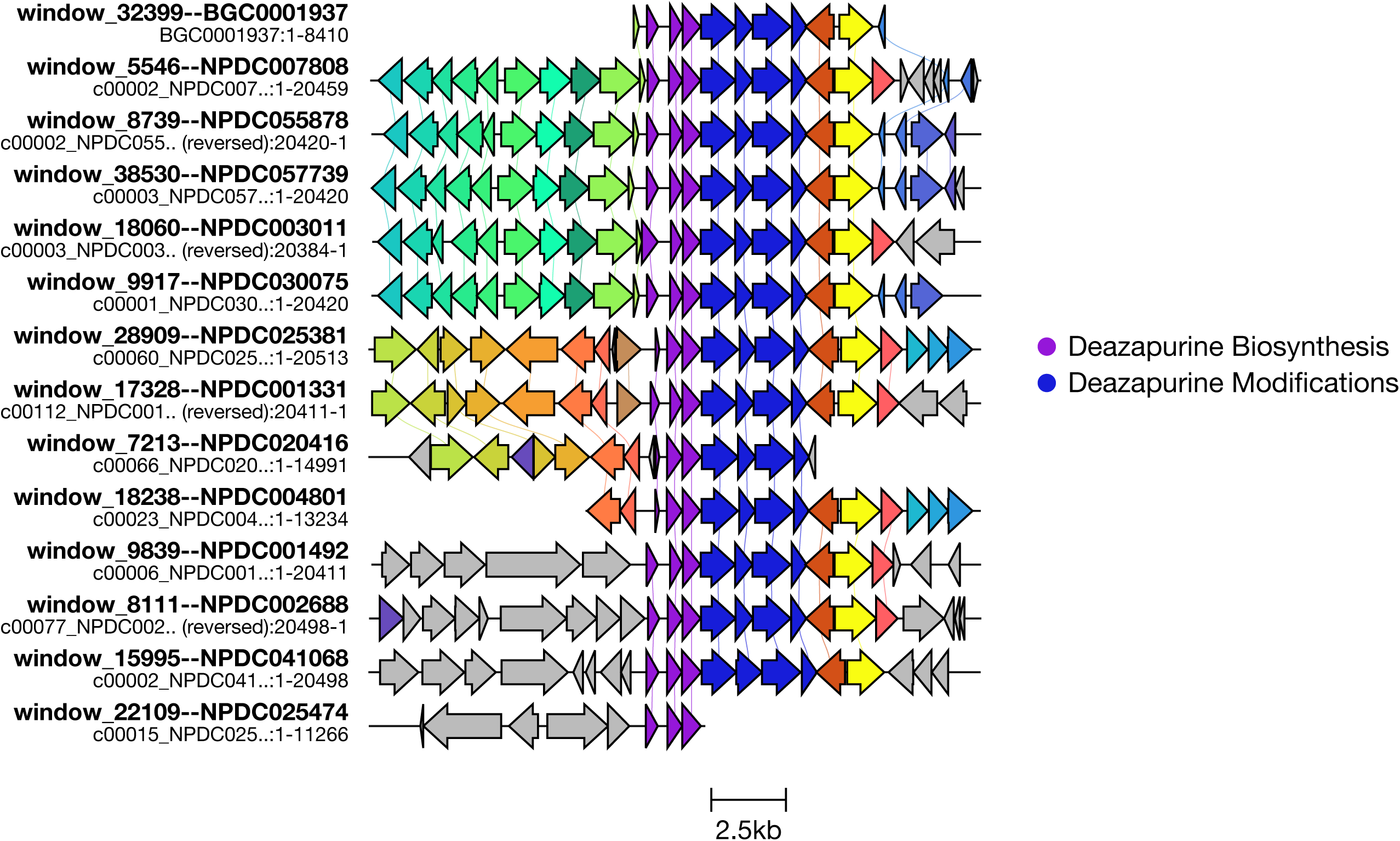

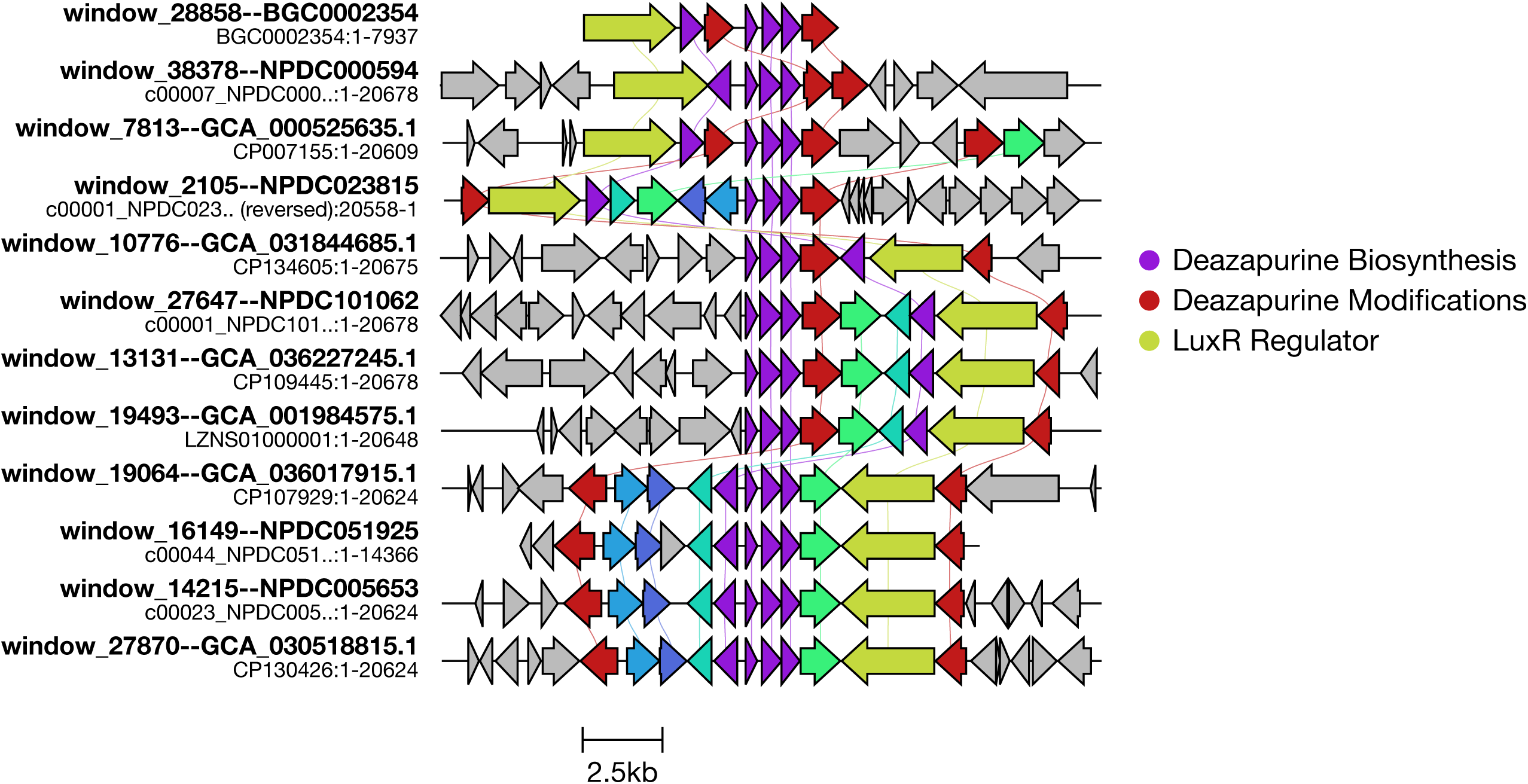

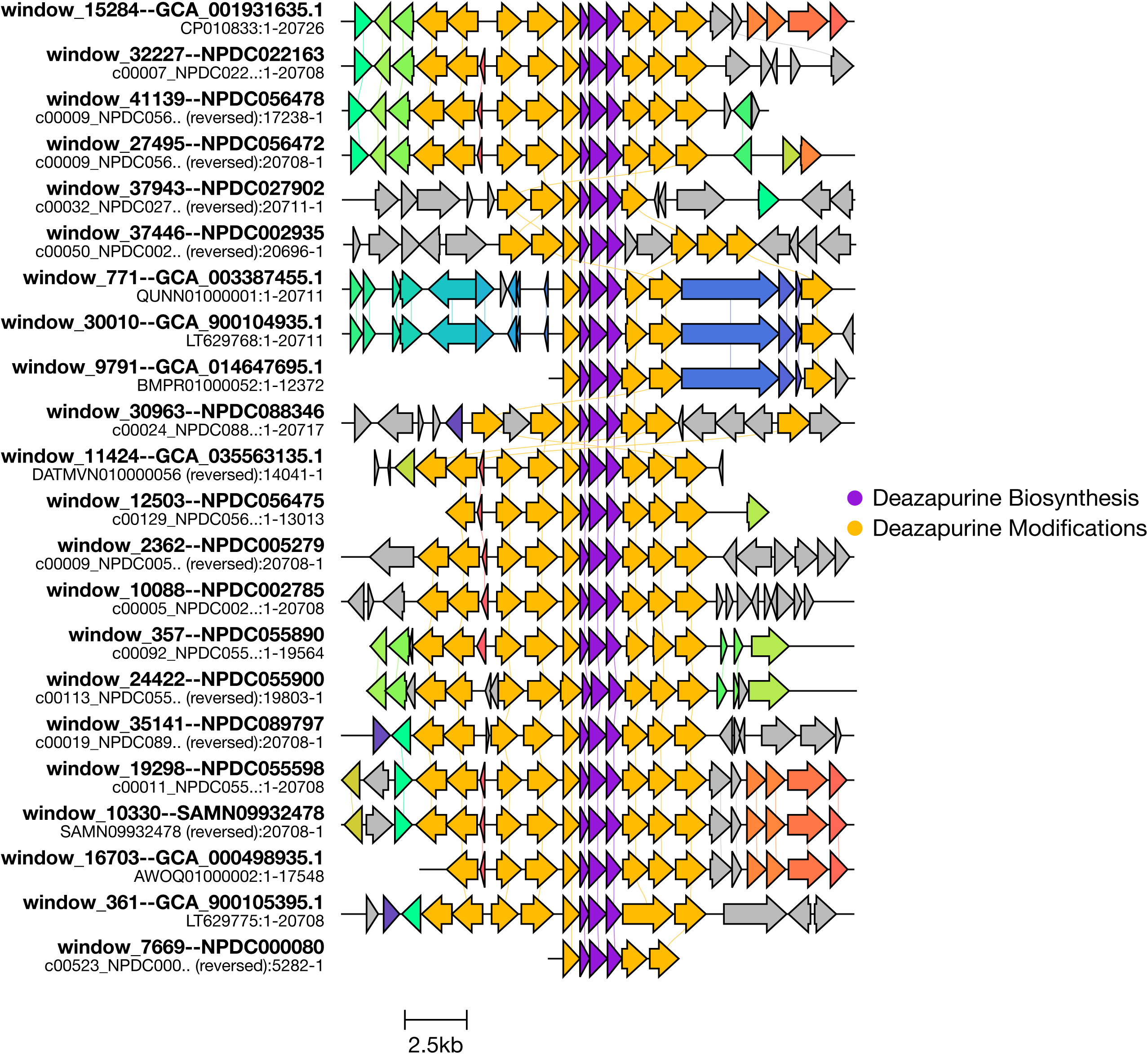

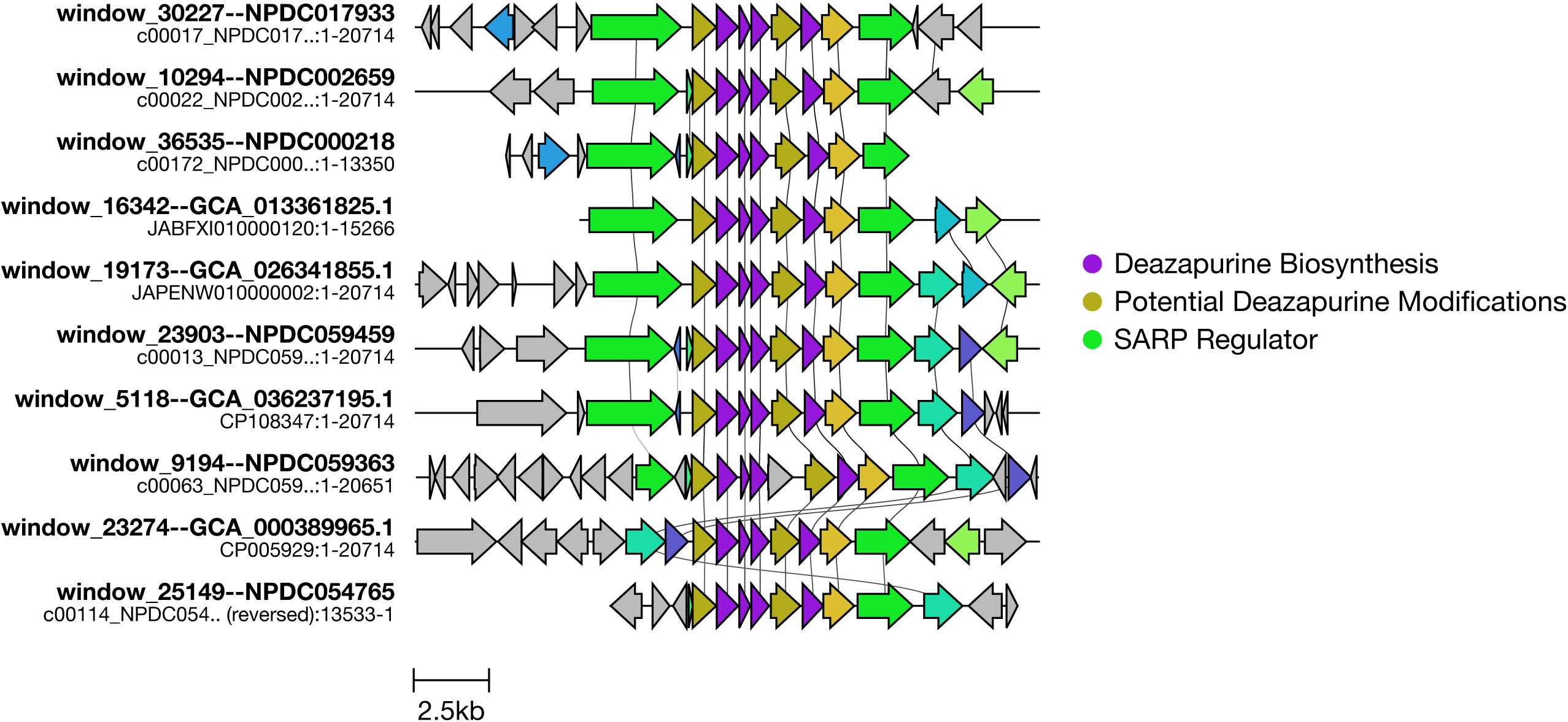

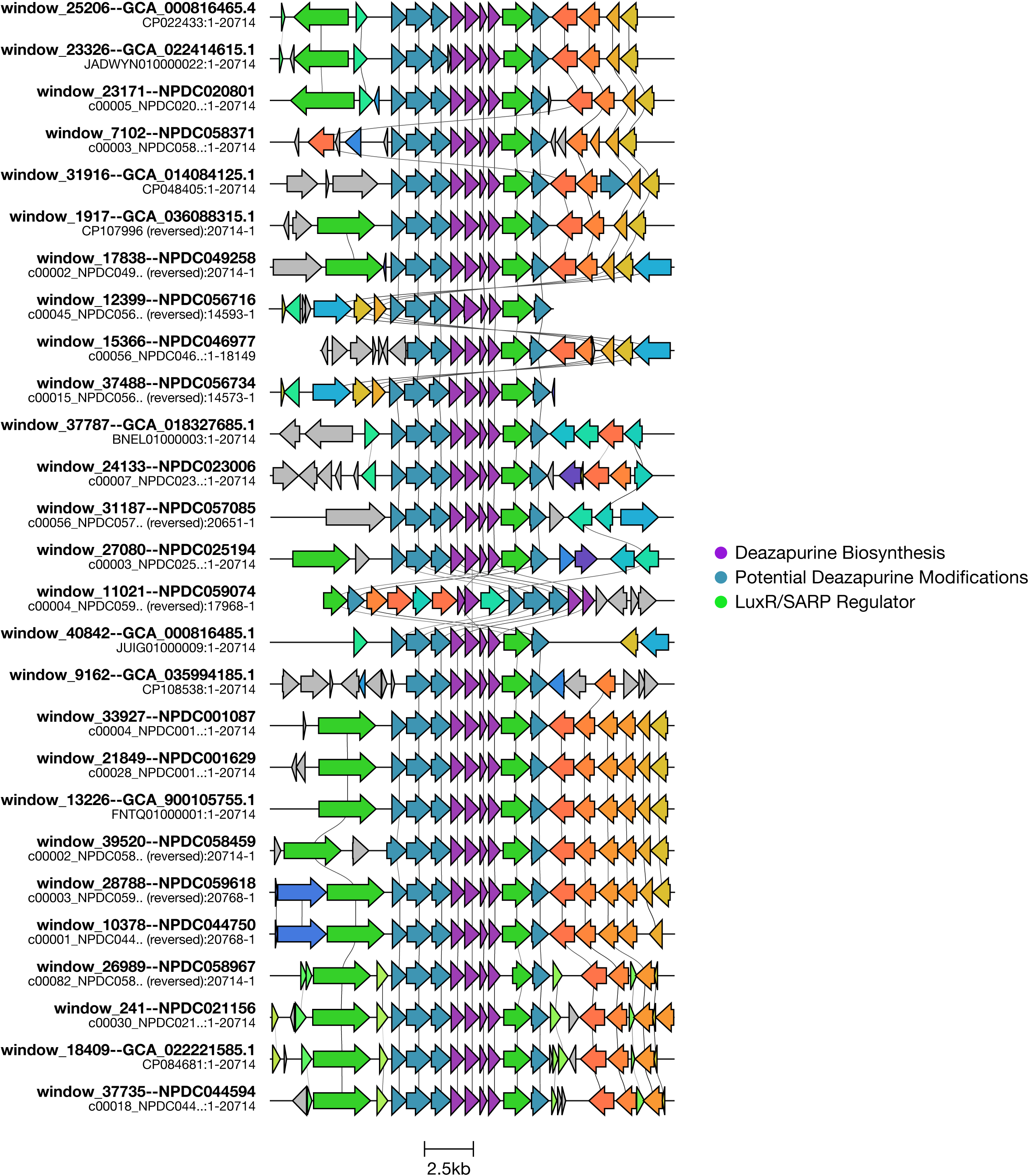

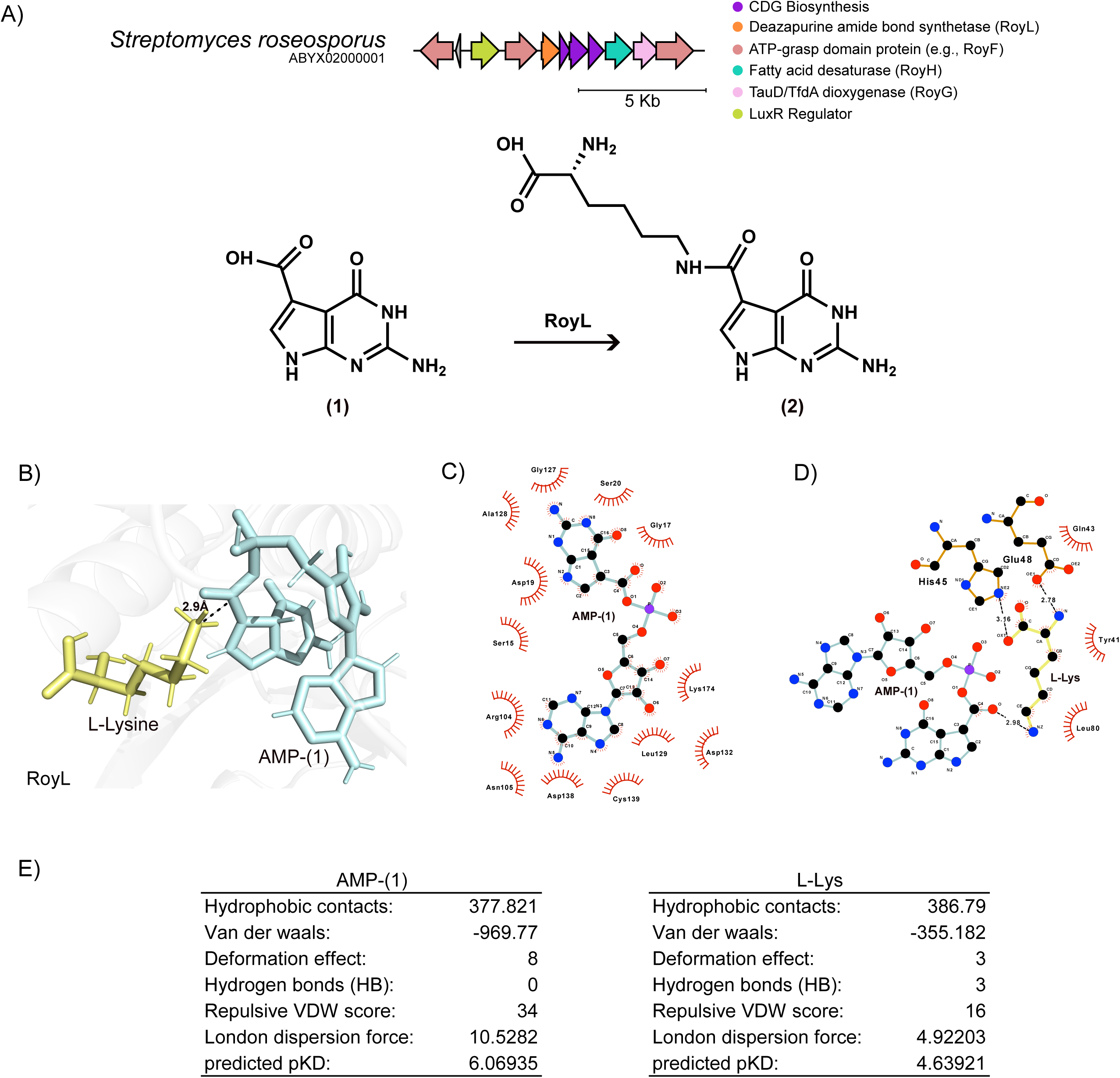

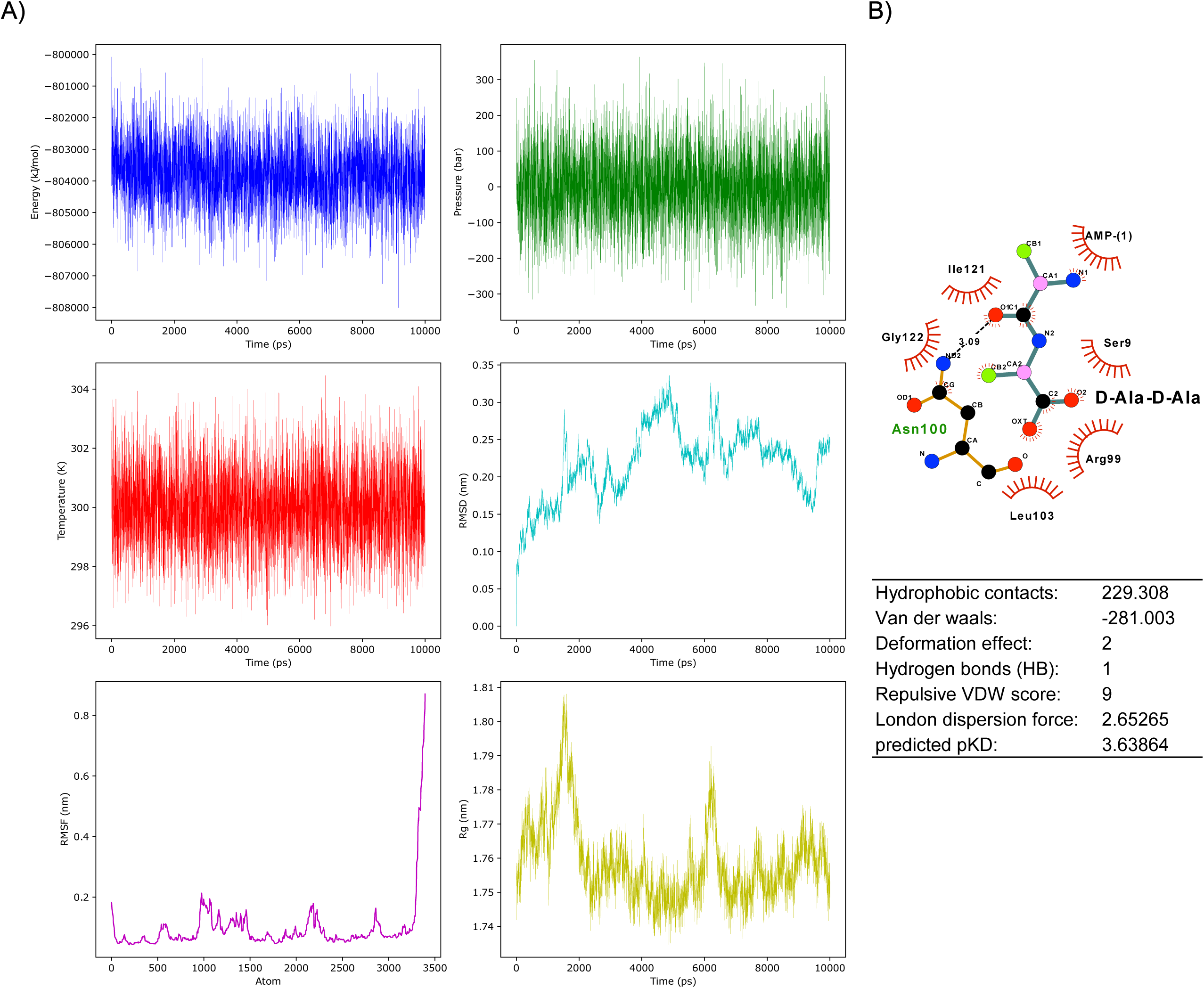

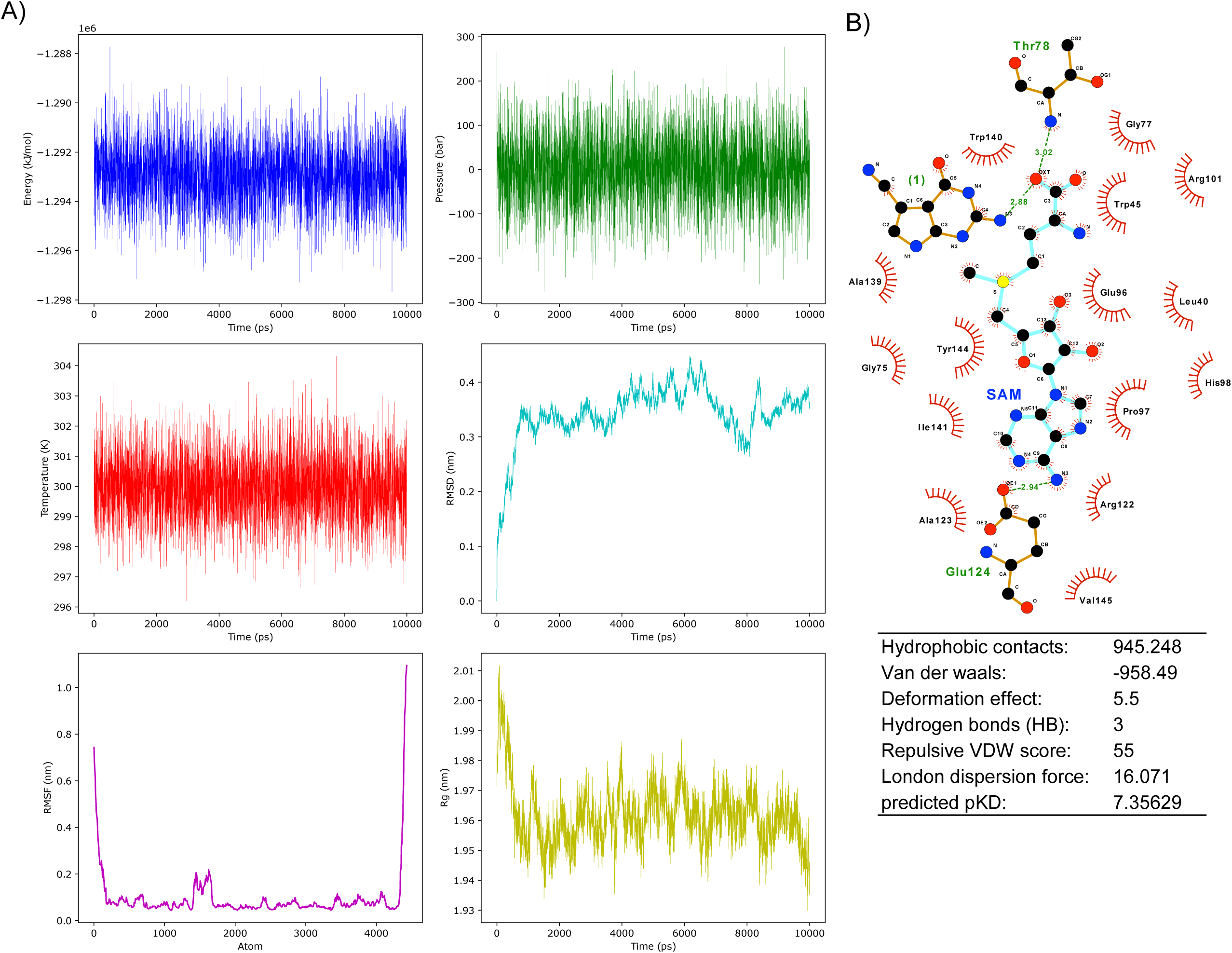

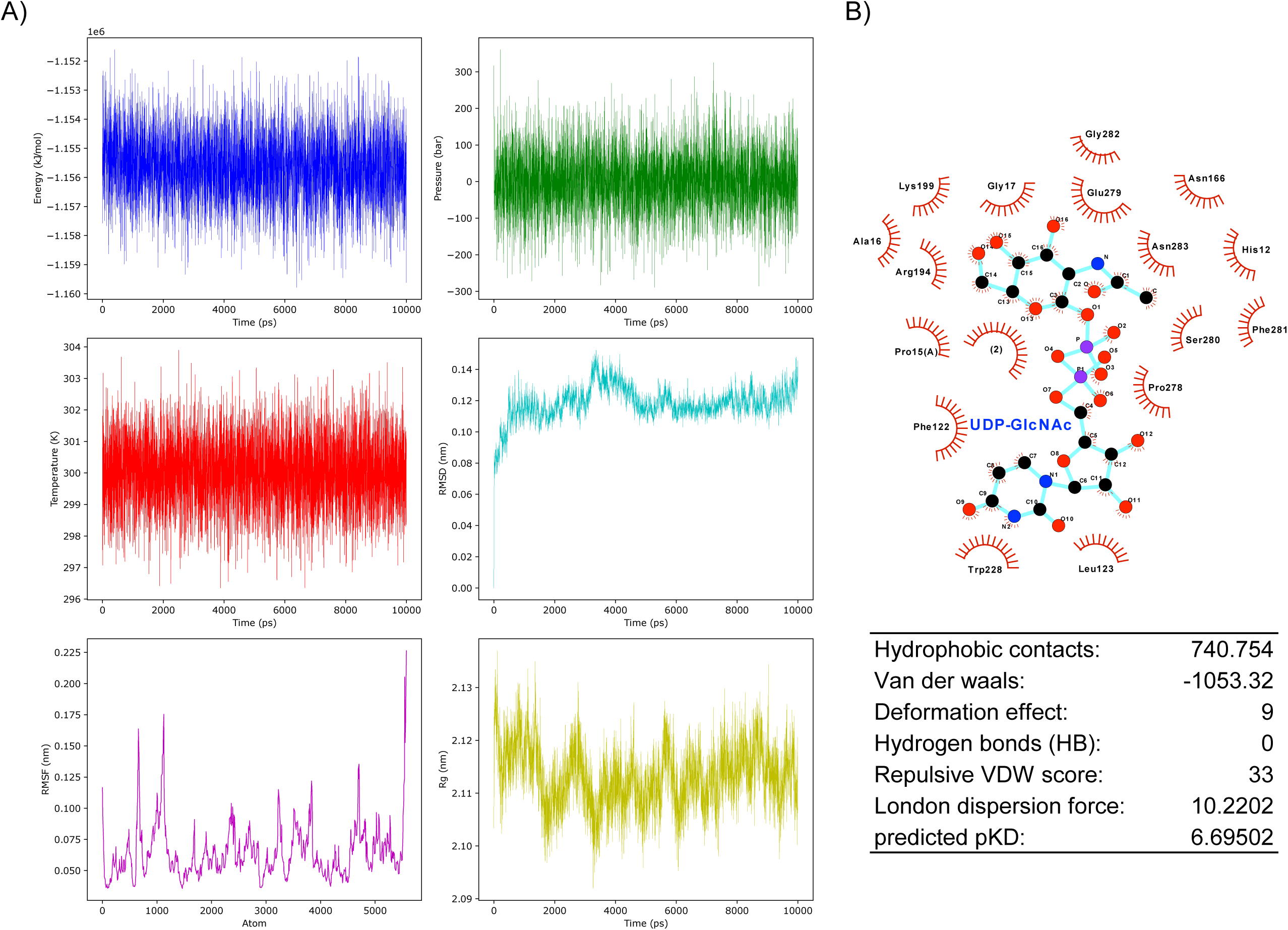

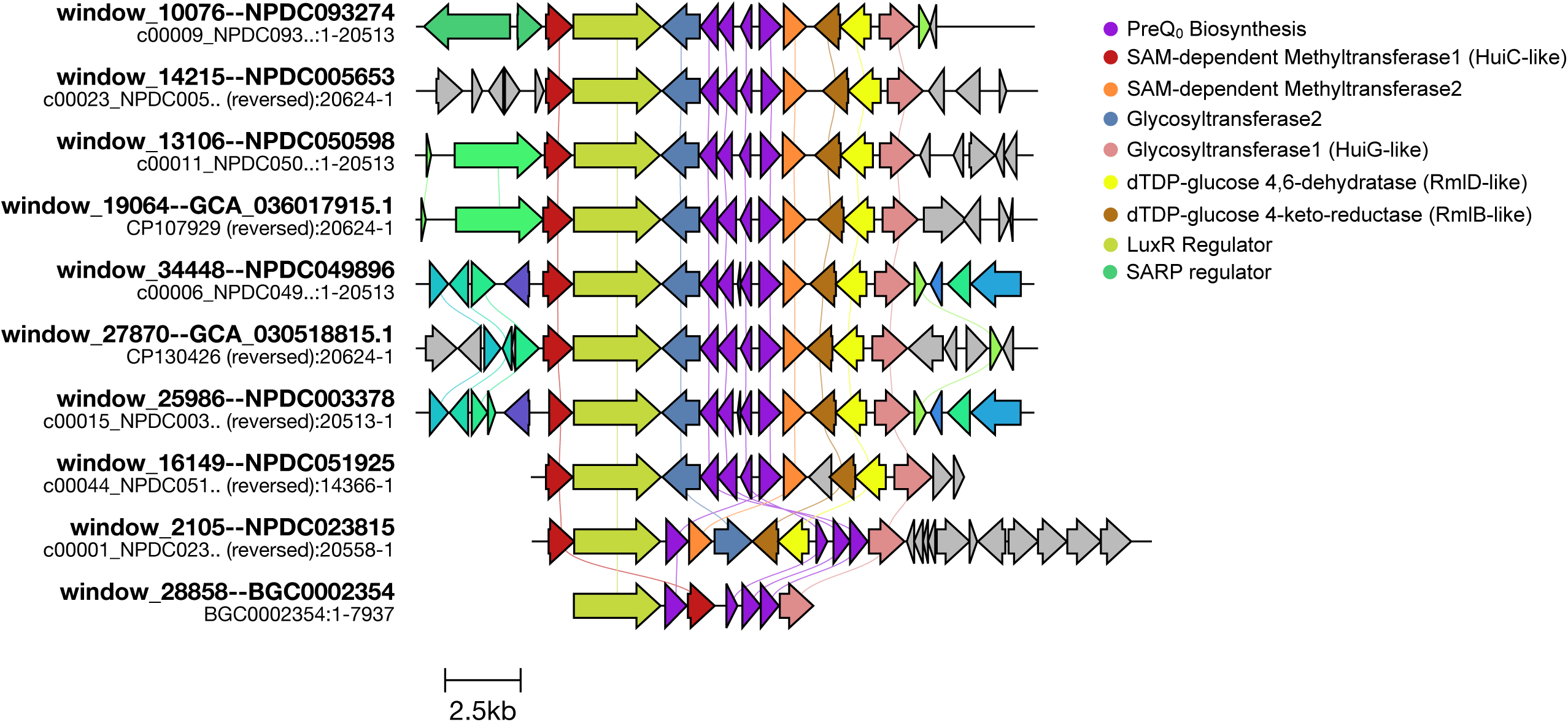

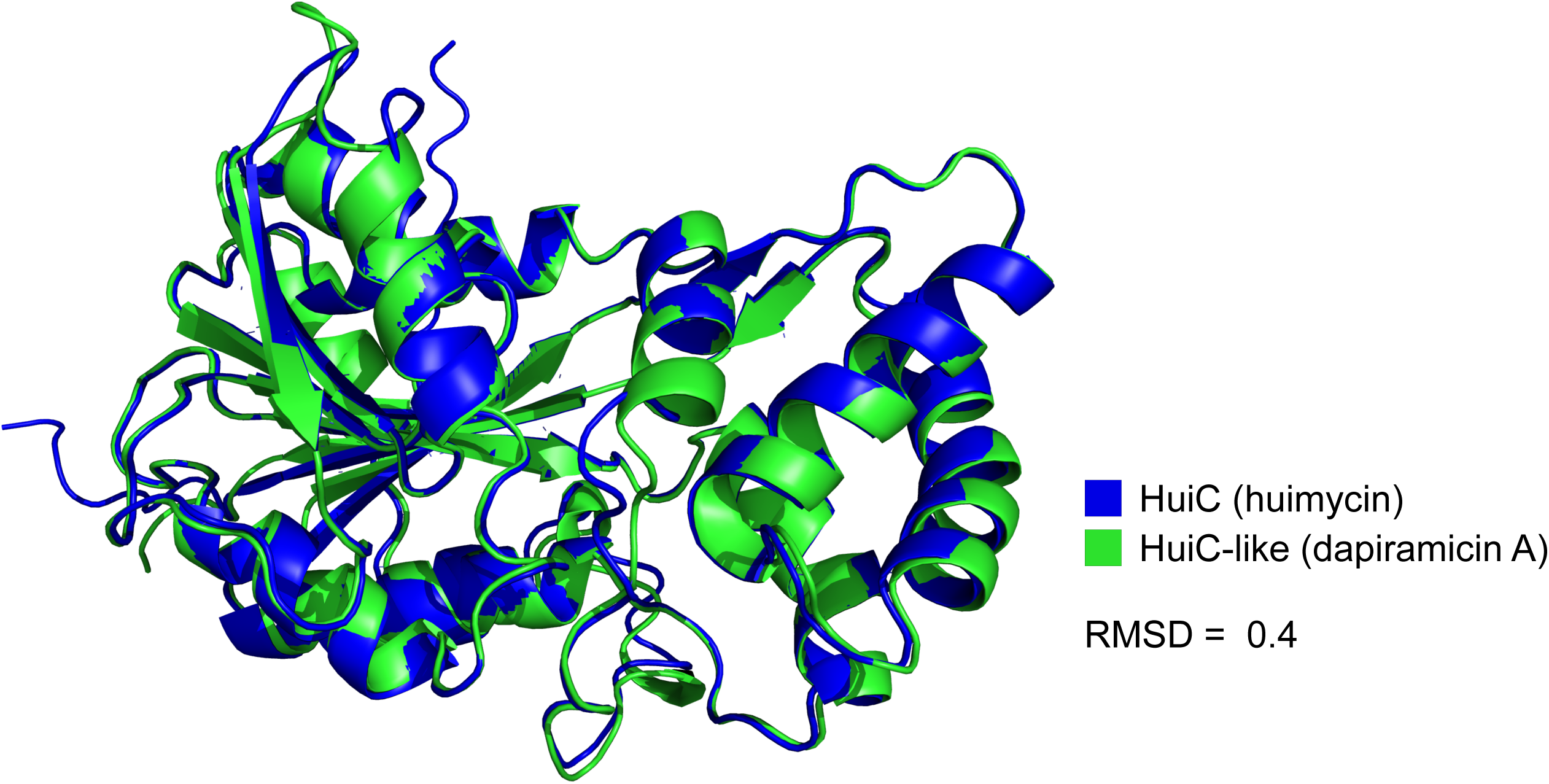

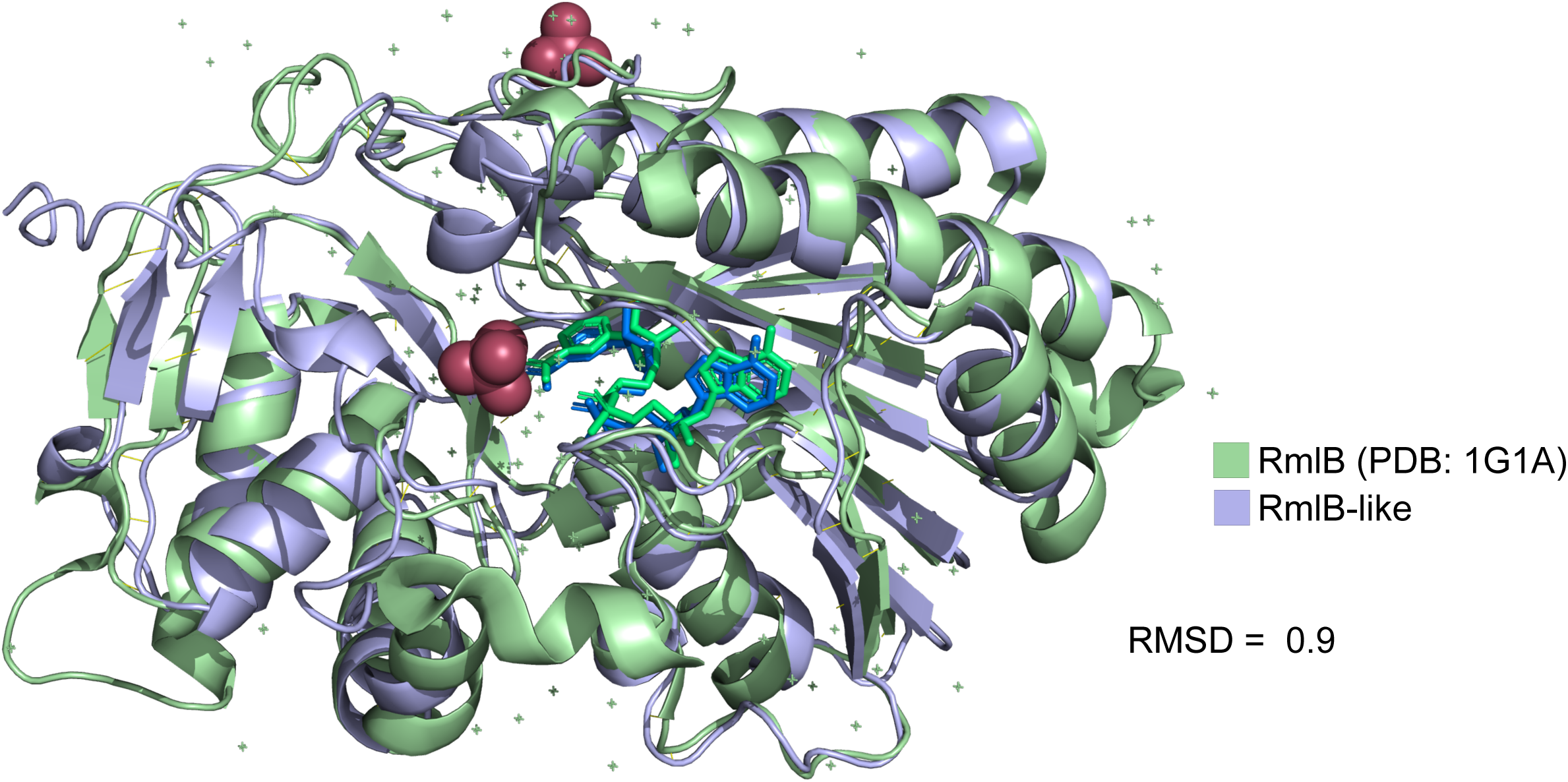

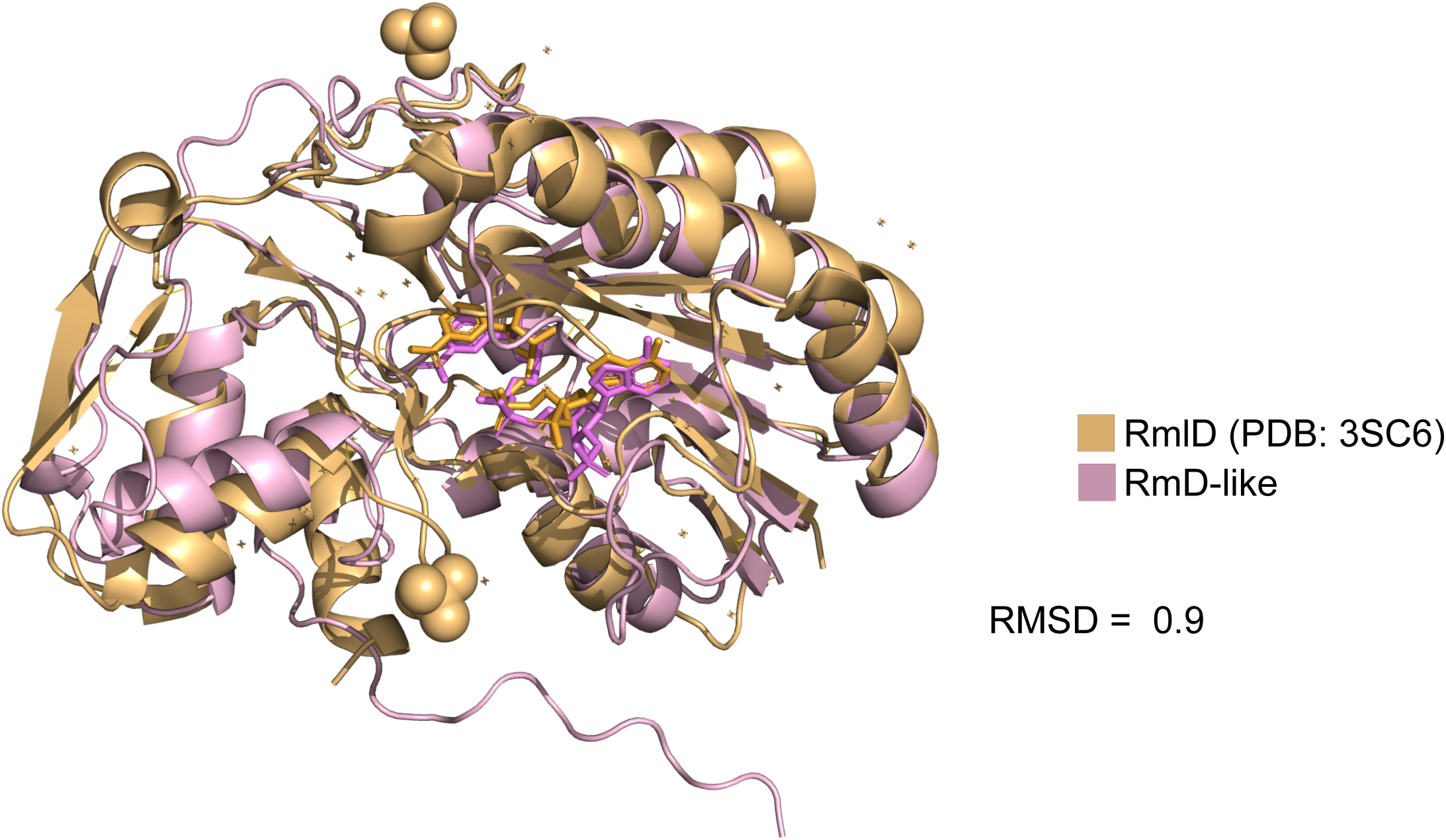

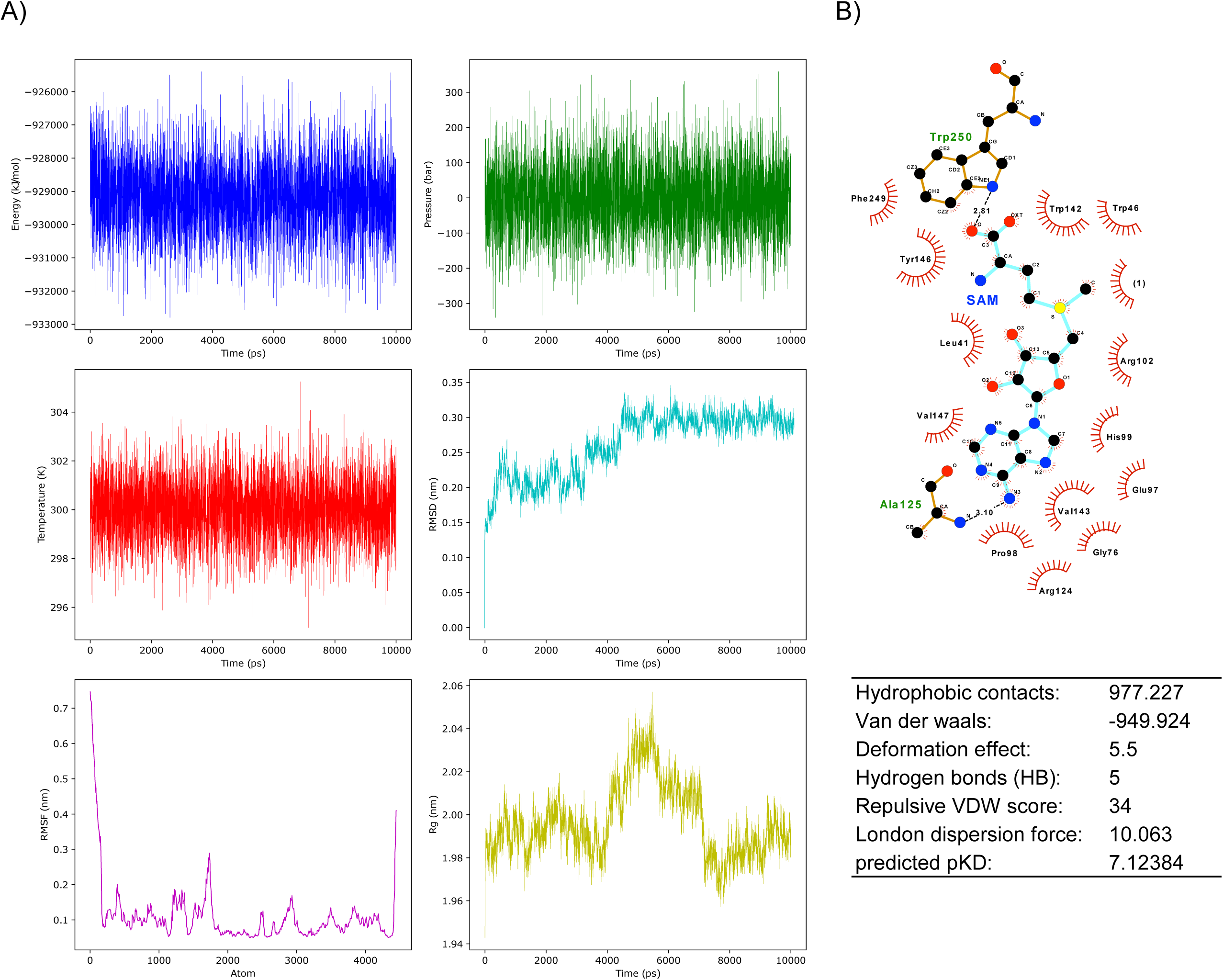

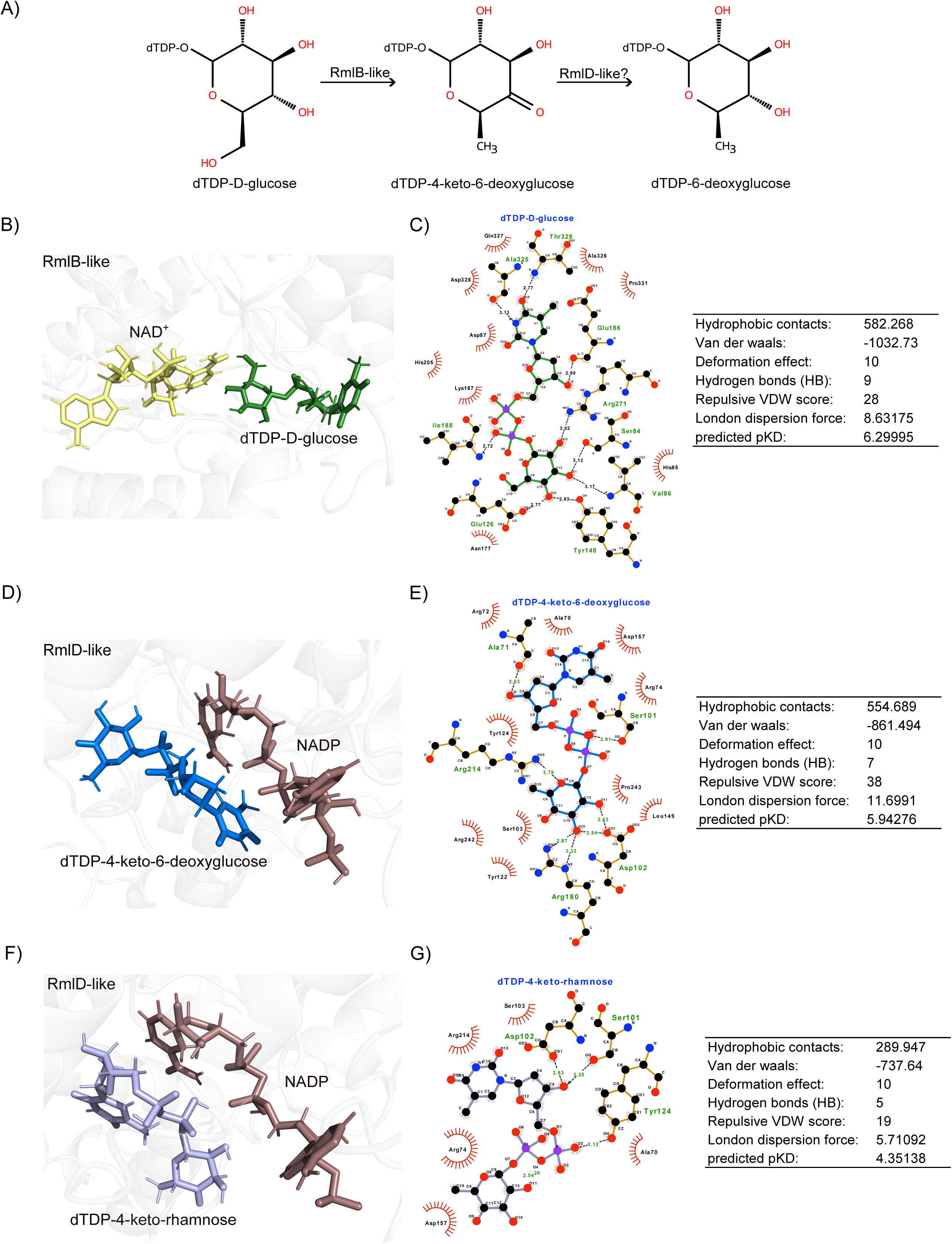

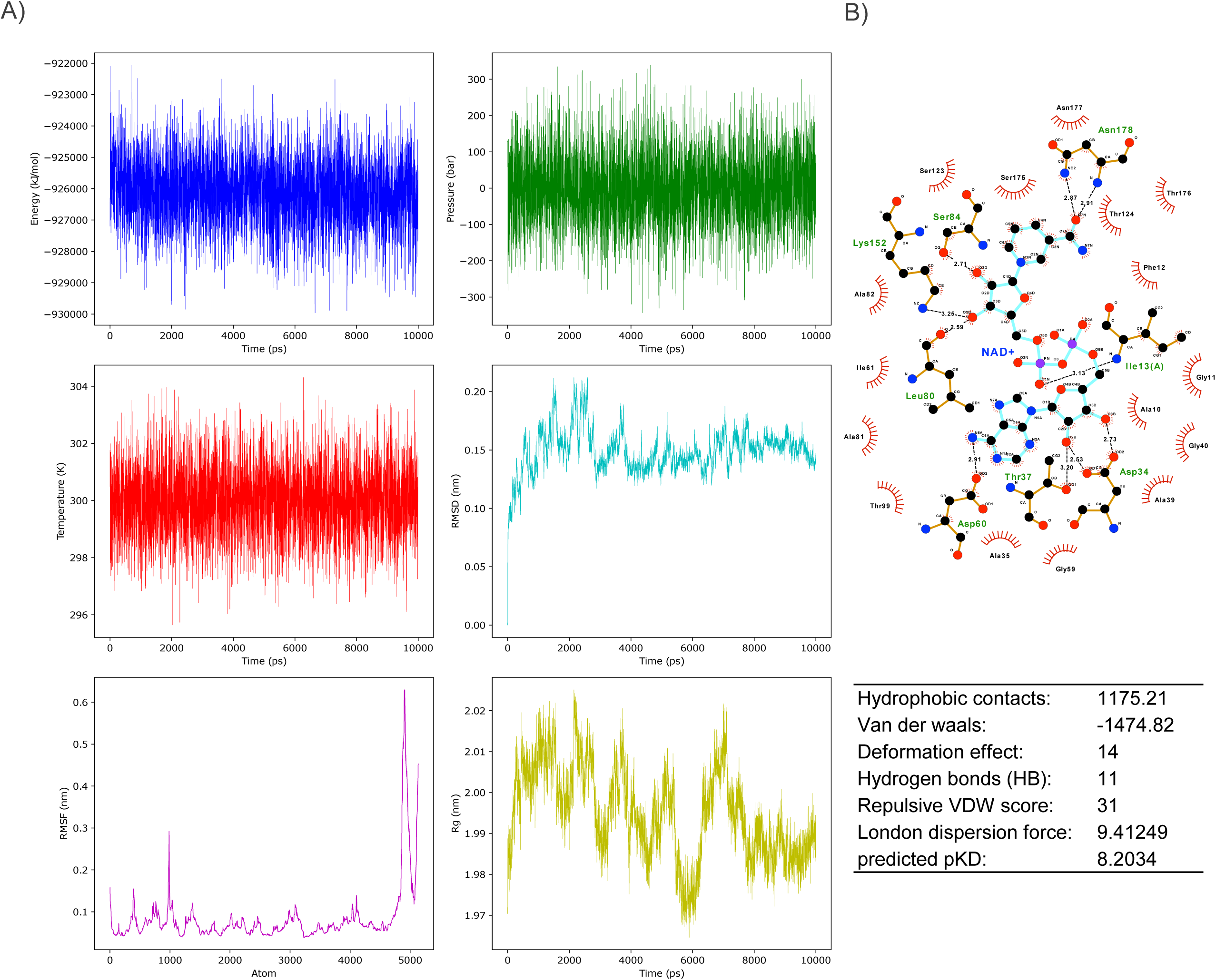

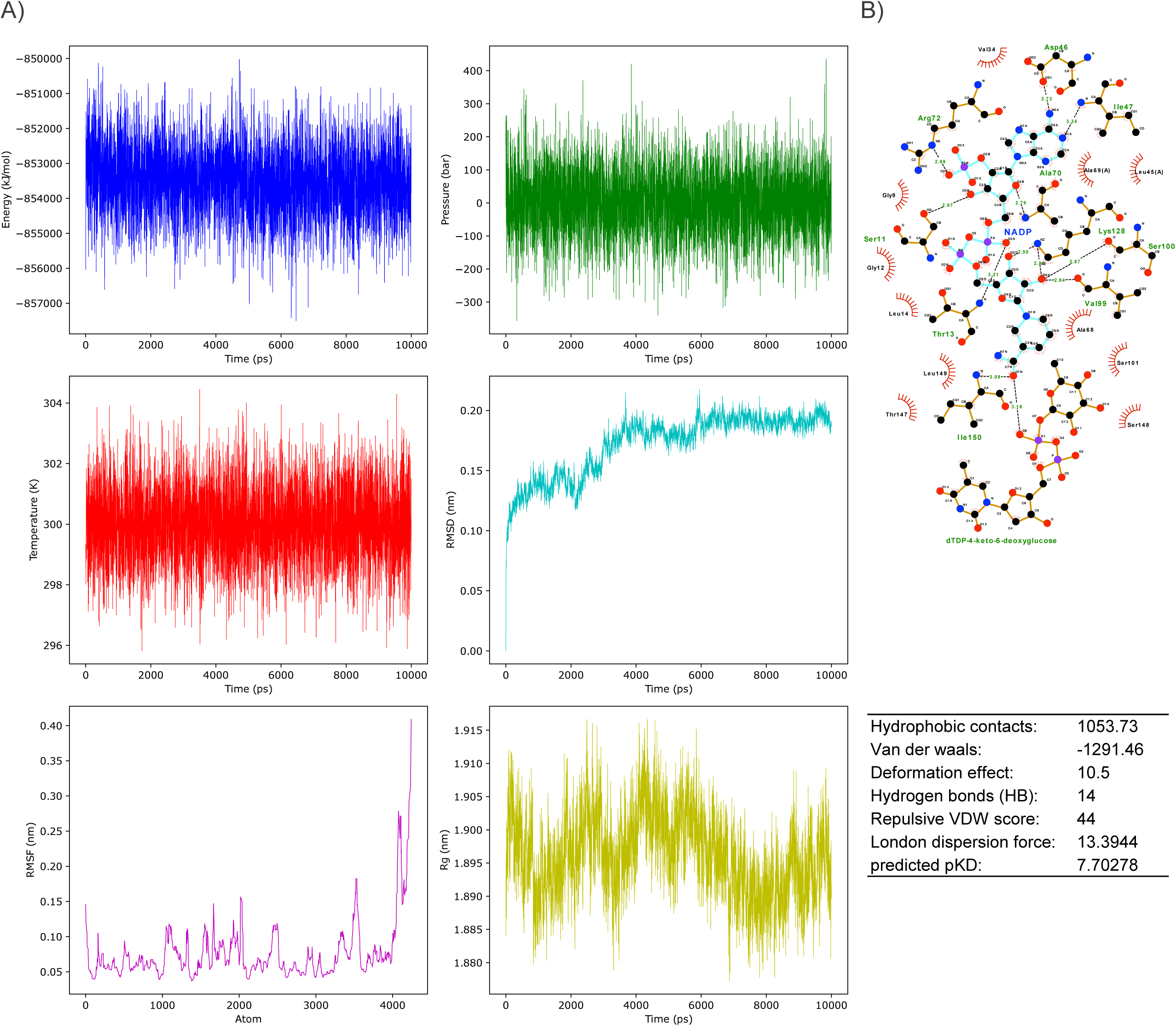

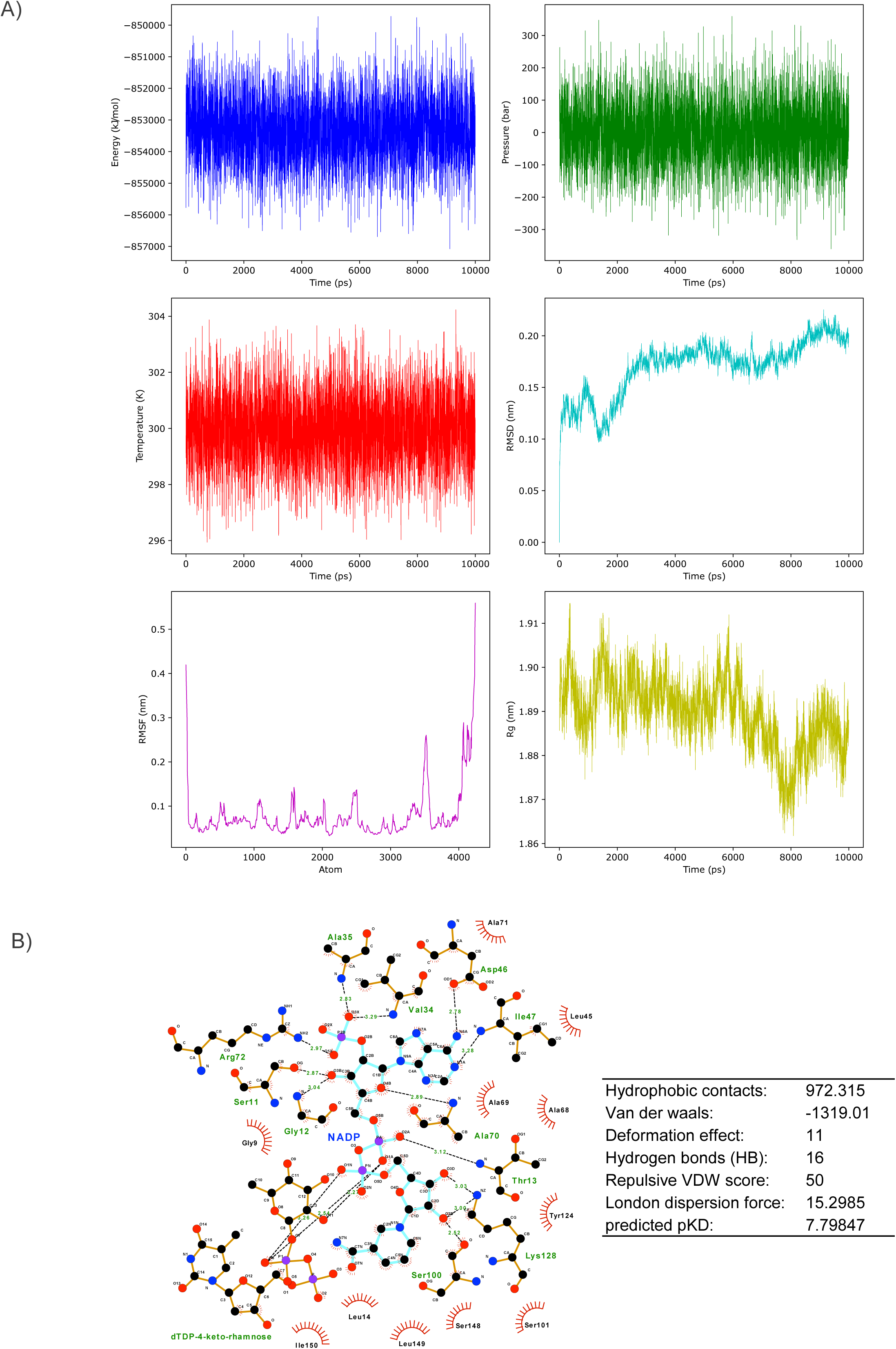

