## Supplemental Material for "Integrating targeted genome mining and structure-guided modeling reveals unexplored 7-deazapurine-containing pathways"

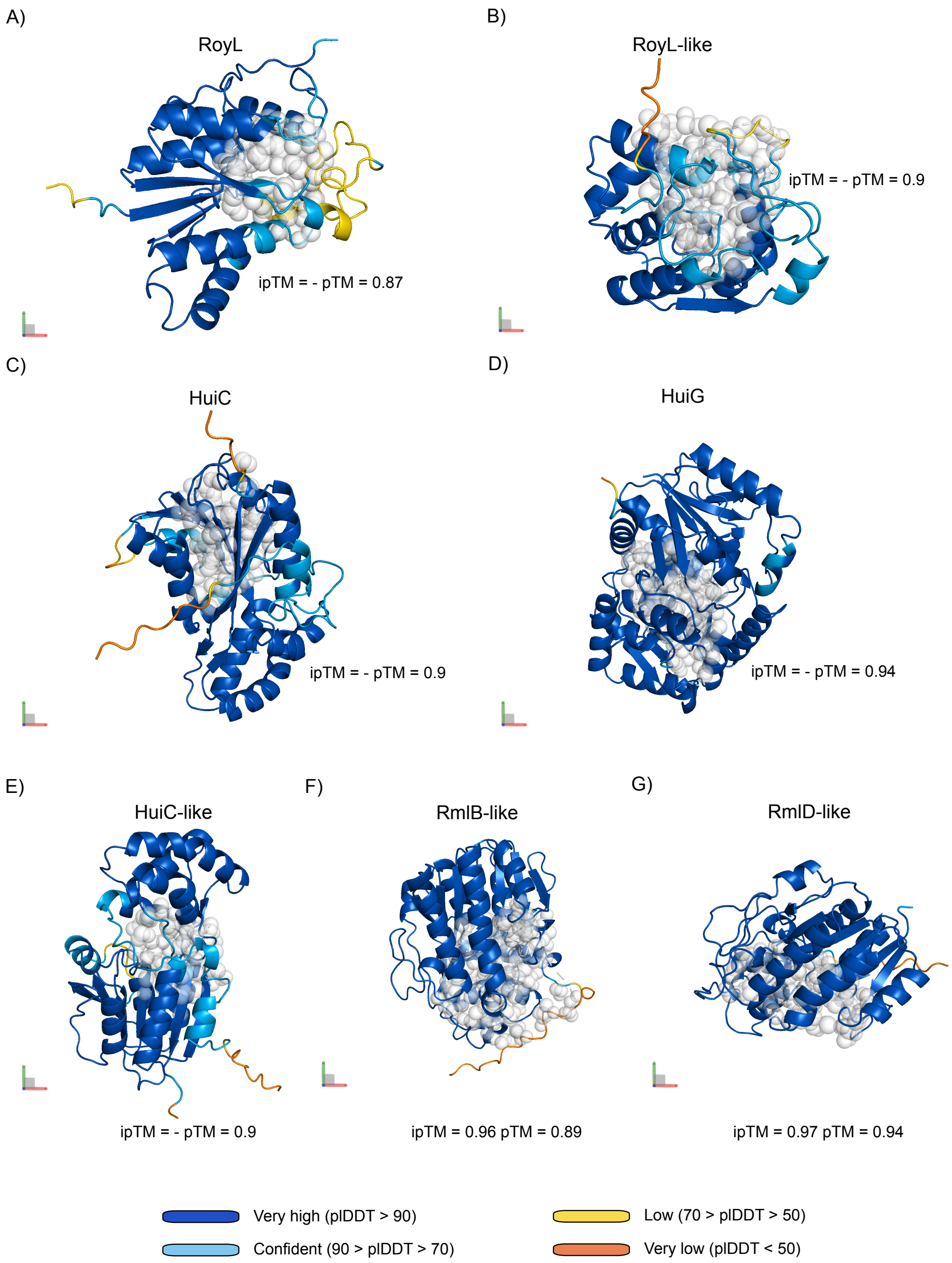


**Supplementary Figure 1.** AlphaFold-predicted structures of representative enzymes from distinct deazapurine biosynthetic pathways. Deazapurine-bond synthetases from *Streptomyces rosesporus* (SSIG_07344) and *Streptomyces rapamycinicus* NRRL 5491 (D3C57_103080) are shown in panels A-B; huimycin-associated SAM-dependent methyltransferase (KALB_4069) and glycosyltransferase (KALB_4073) in C-D; and dapiramicin-associated HuiC-like (ctg9_31), RmlB-like (ctg9_40), and RmlD-like (ctg9_39) enzymes in E-G. Structures are displayed as cartoons colored by per-residue confidence (pLDDT) with predicted binding pockets shown as gray spheres; ipTM and pTM scores are indicated to reflect structural and interface confidence.

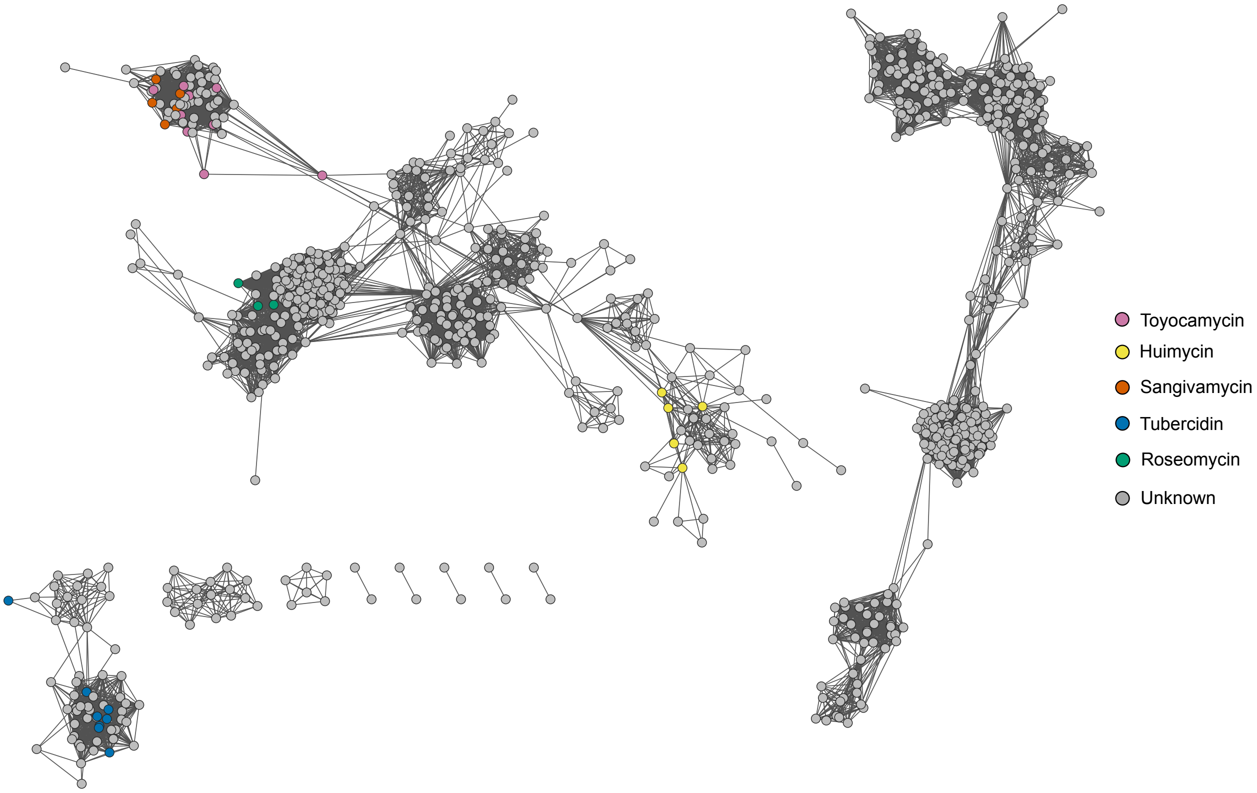


**Supplementary Figure 2**. Landscape of 7-deazapurine biosynthetic diversity according to BiG-SCAPE 2.0 using a cutoff of 0.75. BGC members corresponding to experimentally validated MIBiG pathways are highlighted. Singleton subfamilies (those containing only one BGC) were excluded to improve visualization.

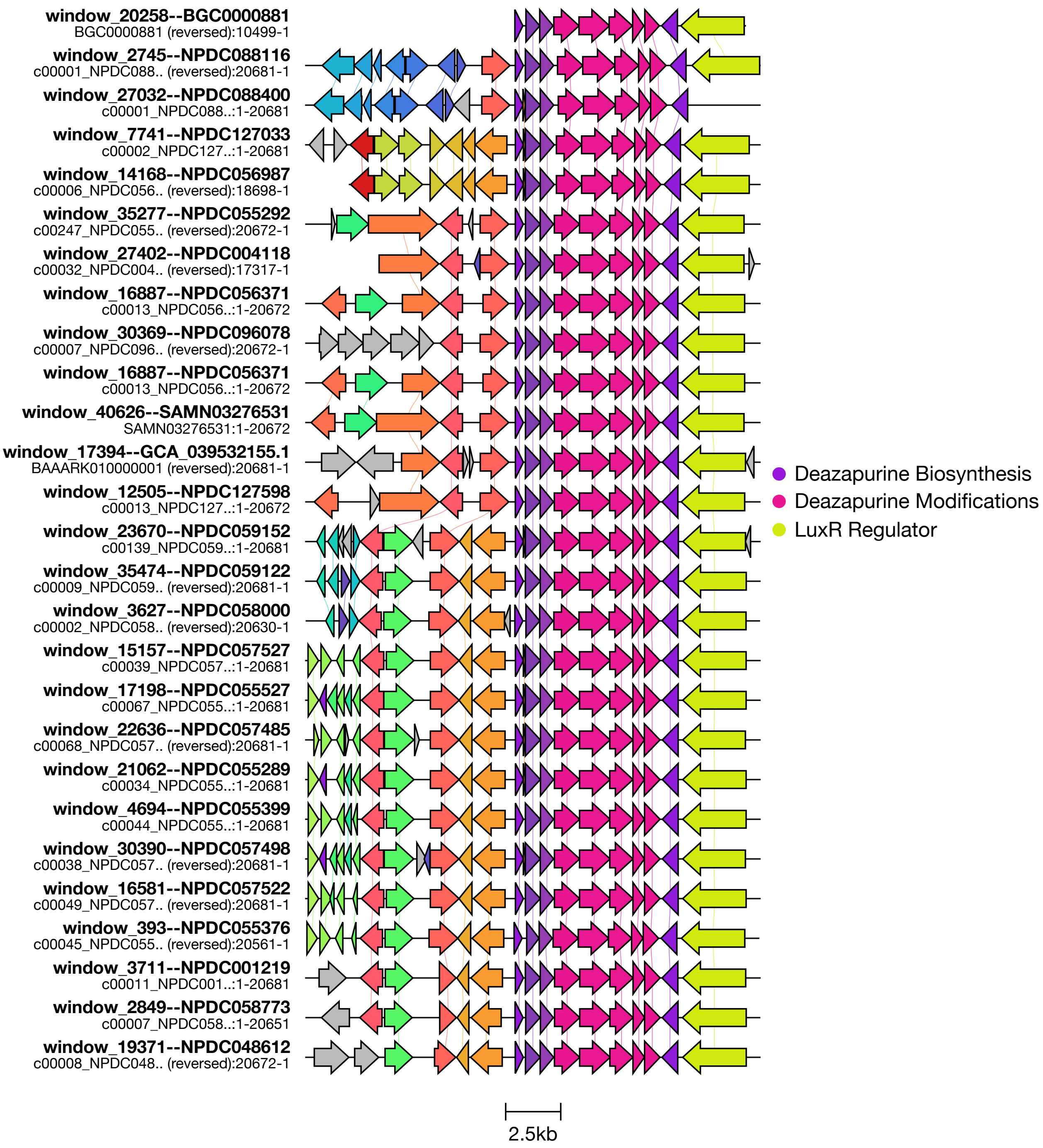


**Supplementary Figure 3.** Genomic neighborhoods of GATOR-GC–predicted members of the toyocamycin subfamily. The experimentally validated toyocamycin BGC from MIBiG is shown at the top for reference.

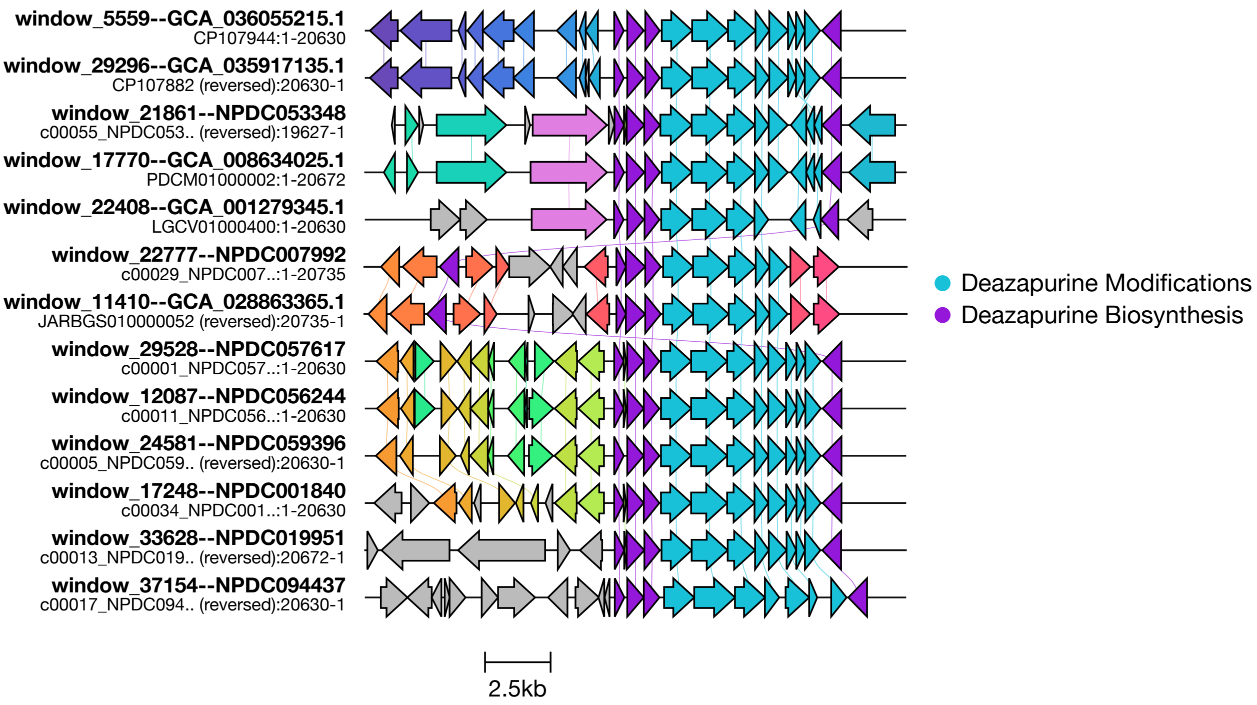


**Supplementary Figure 4.** Genomic neighborhoods of GATOR-GC–predicted members of the sangivamycin subfamily.

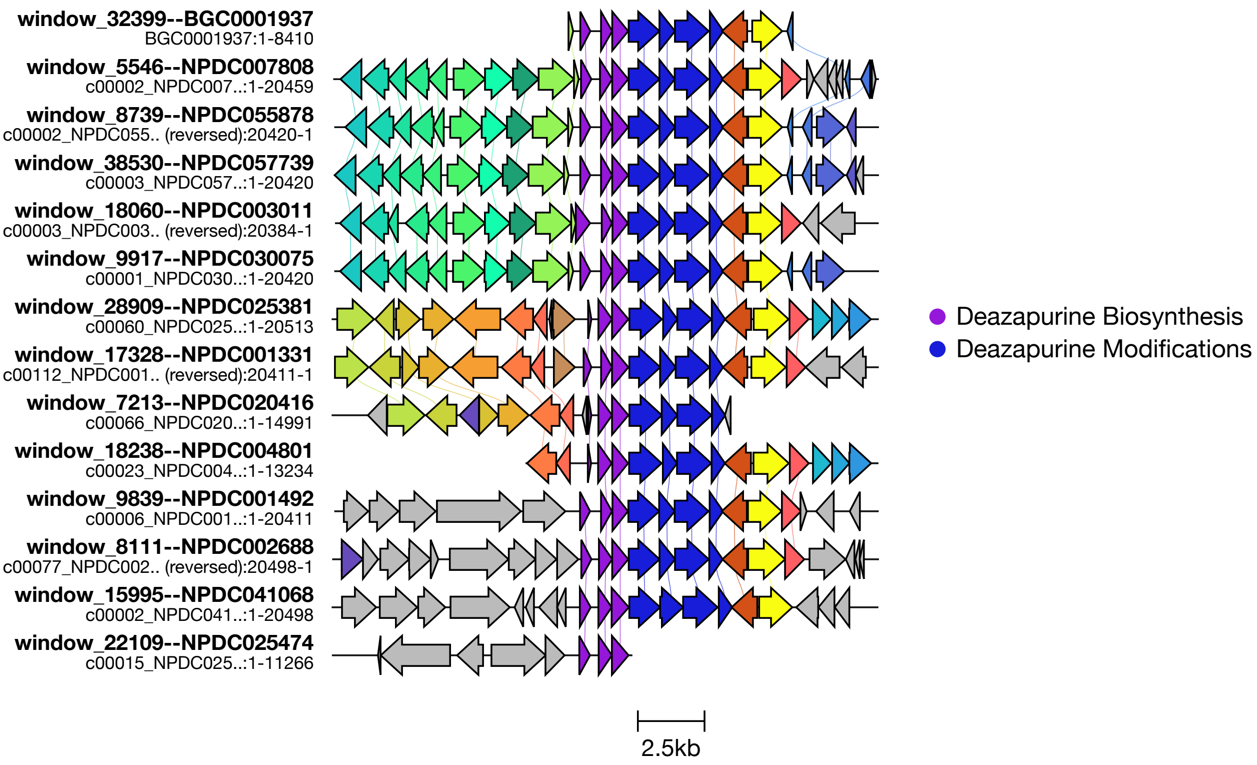


**Supplementary Figure 5.** Genomic neighborhoods of GATOR-GC–predicted members of the tubercidin subfamily. The experimentally validated tubercidin BGC from MIBiG is shown at the top for reference.

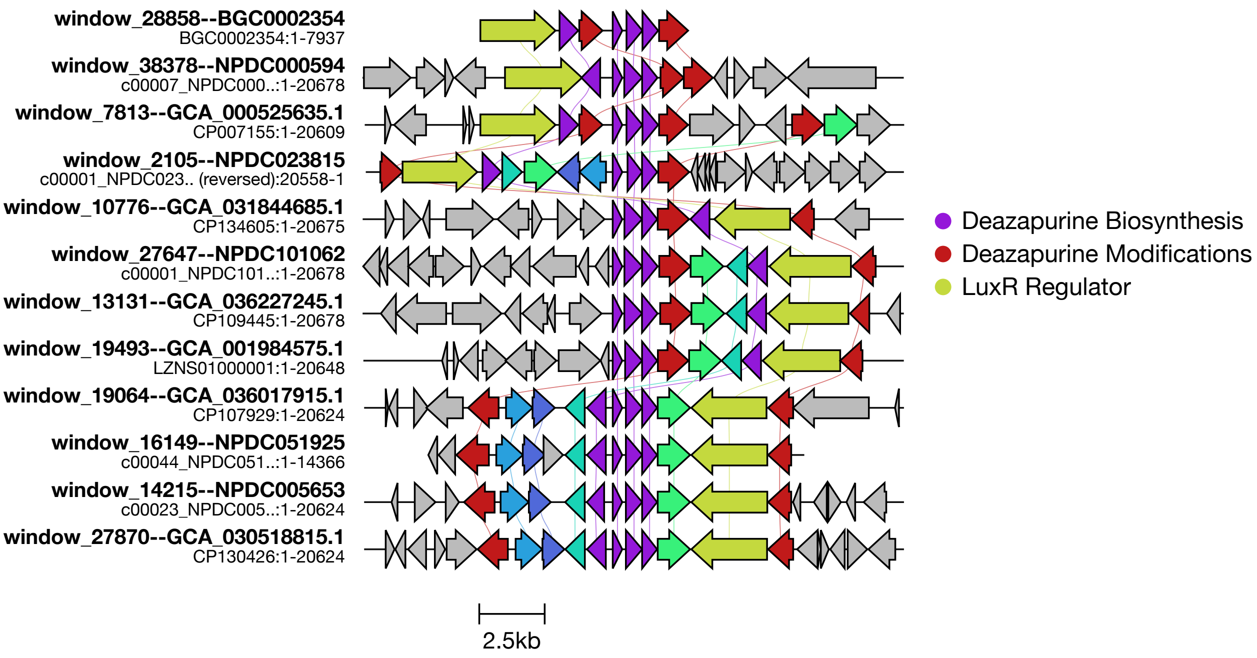


**Supplementary Figure 6.** Genomic neighborhoods of GATOR-GC–predicted members of the huimycin subfamily. The experimentally validated huimycin BGC from MIBiG is shown at the top for reference.

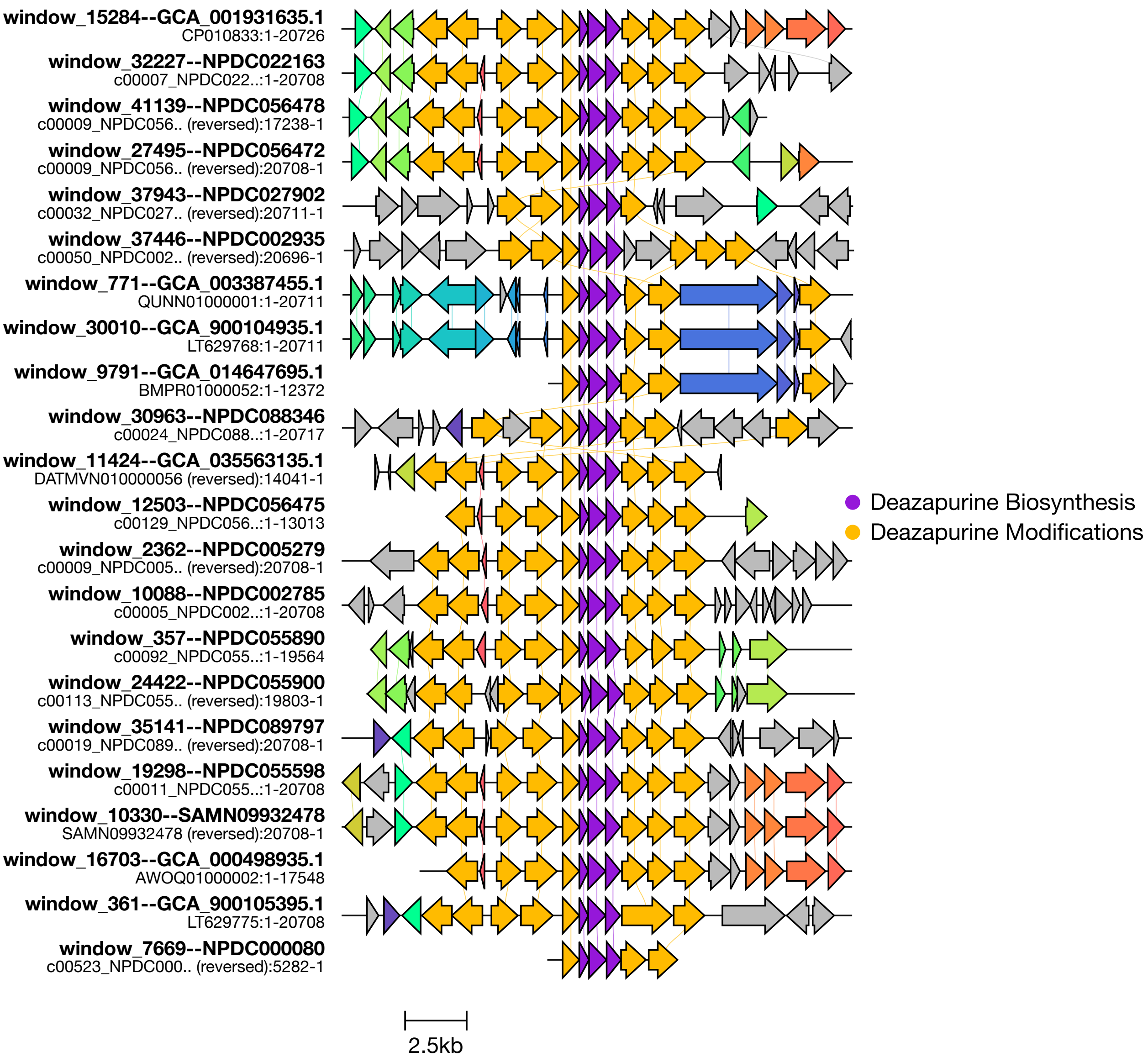


**Supplementary Figure 7.** Genomic neighborhoods of GATOR-GC–predicted members of the roseomycin subfamily.

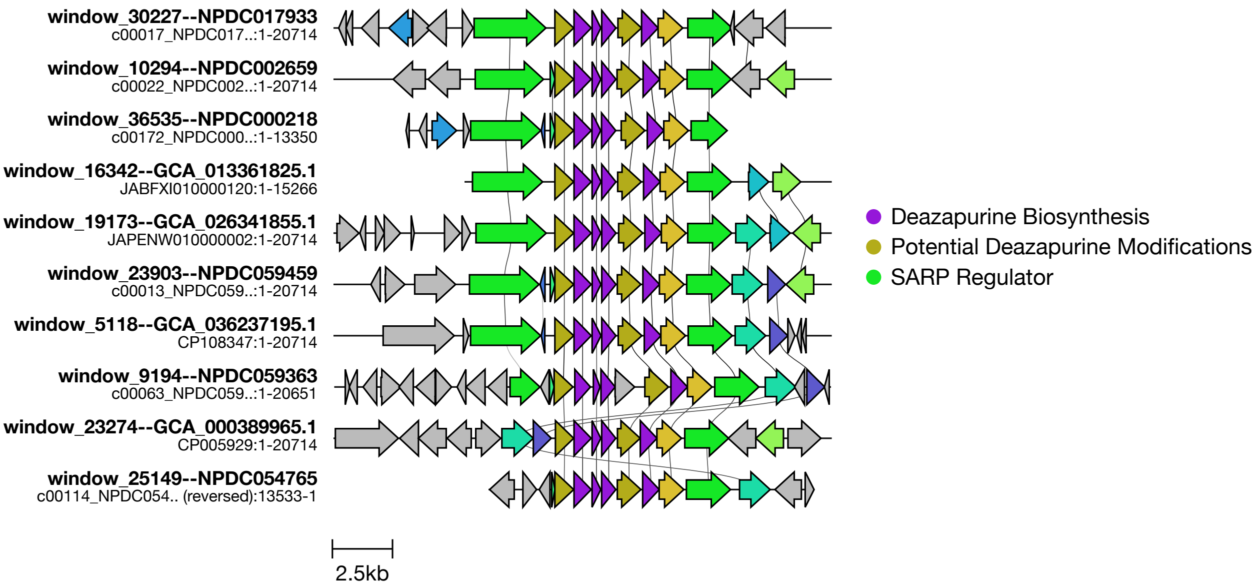


**Supplementary Figure 8.** Genomic neighborhoods of GATOR-GC–predicted members of the biggest uncharacterized subfamily. Only 10 representatives out of 81 BGCs are shown. These candidate BGCs harbor different tailoring enzymes, including non-heme dioxygenases (PF14226), Asp/Asn β-hydroxylases (PF05118), and, in some clusters, SAM-dependent methyltransferases (PF13649). All members encode transcriptional regulators with bacterial activator domains and tetratricopeptide repeats (PF03704).

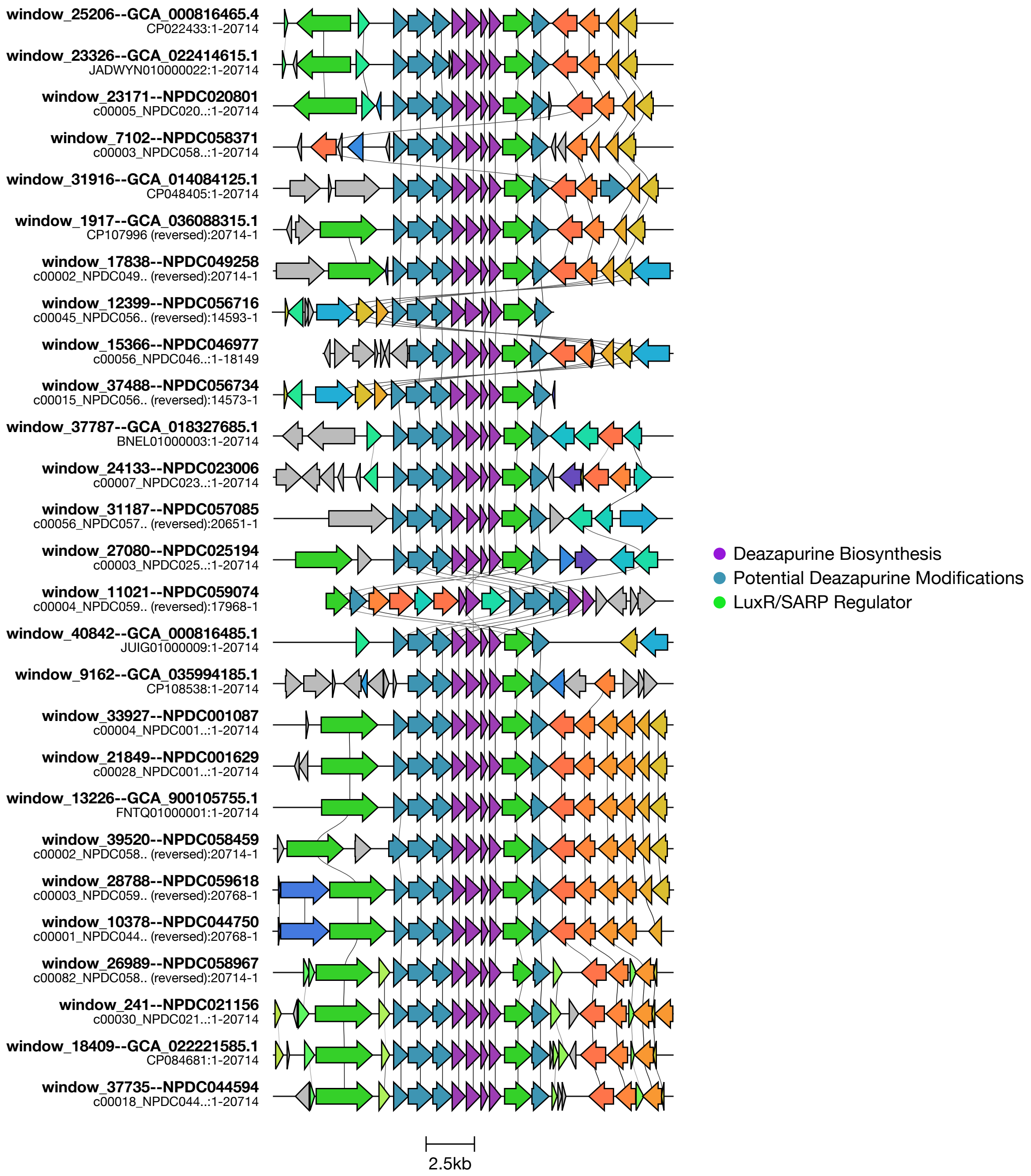


**Supplementary Figure 9.** Genomic neighborhoods of GATOR-GC–predicted members of another uncharacterized subfamily. These candidate BGCs harbor different tailoring enzymes, including non-heme dioxygenases (PF14226), aminotransferases (PF00202), FAA hydrolases (PF01557), and SAM-dependent methyltransferases (PF13649).

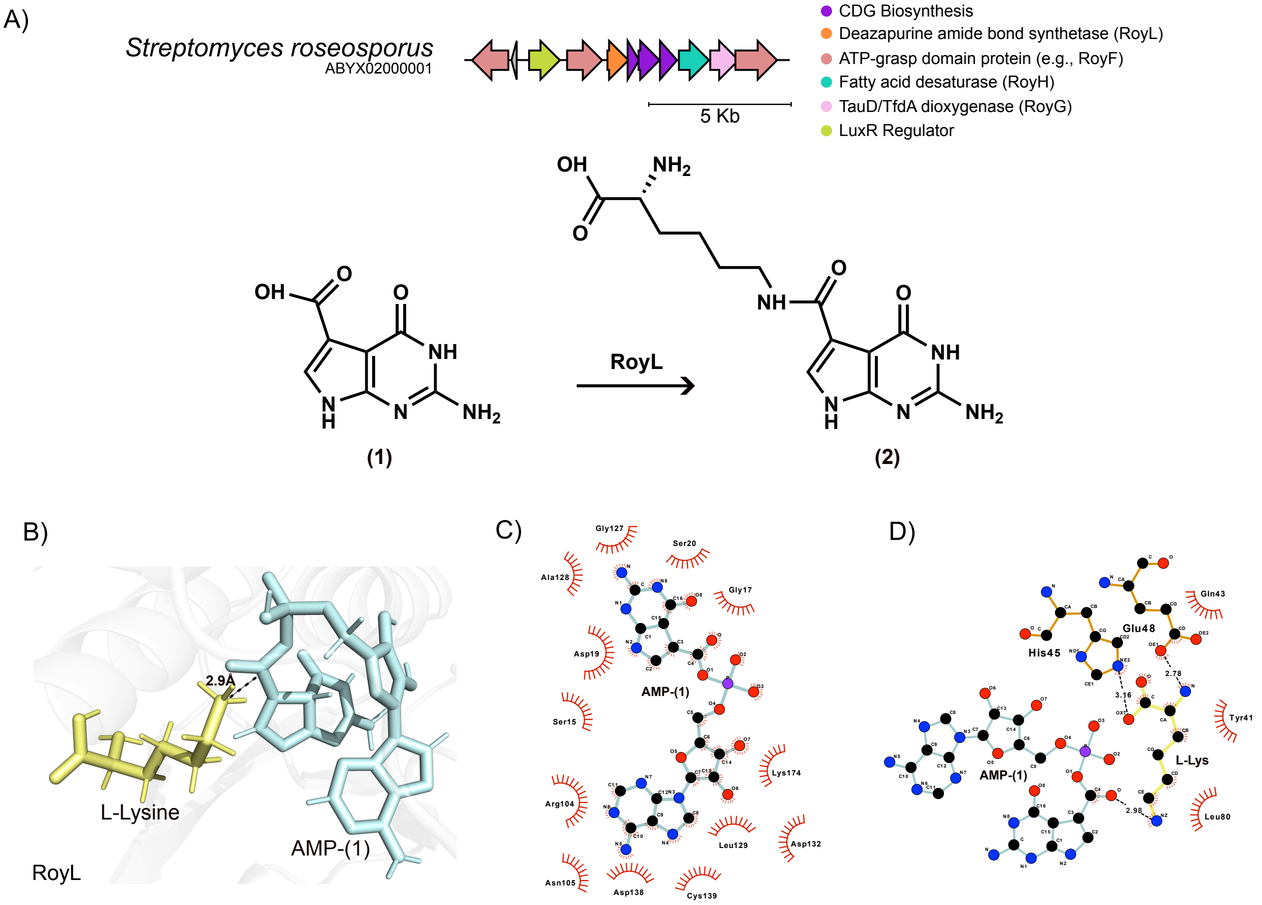


E)

| AMP-(1) | |
|---|---|
| Hydrophobic contacts: | 377.821 |
| Van der waals: | -969.77 |
| Deformation effect: | 8 |
| Hydrogen bonds (HB): | 0 |
| Repulsive VDW score: | 34 |
| London dispersion force: | 10.5282 |
| predicted pKD: | 6.06935 |

| L-Lys | |
|---|---|
| Hydrophobic contacts: | 386.79 |
| Van der waals: | -355.182 |
| Deformation effect: | 3 |
| Hydrogen bonds (HB): | 3 |
| Repulsive VDW score: | 16 |
| London dispersion force: | 4.92203 |
| predicted pKD: | 4.63921 |

**Supplementary figure 10.** Enzyme-substrate interactions in roseomycin A biosynthetic pathway. (A) Roseomycin BGC showing predicted enzyme annotations, and the biosynthetic step catalyzed by RoyL, which forms the amide bond to generate the peptidyl 7-deazapurine metabolite. B) Representative snapshot from production MD simulations of the RoyL in complex with AMP-(1) and L-lys, highlighting a potential pre-reactive amide-bond-forming state. C-D) Key hydrophobic and hydrogen-bond interactions stabilizing substrate binding within the active site, along with the E) predicted binding affinity for AMP-(1) and L-Lys

A)

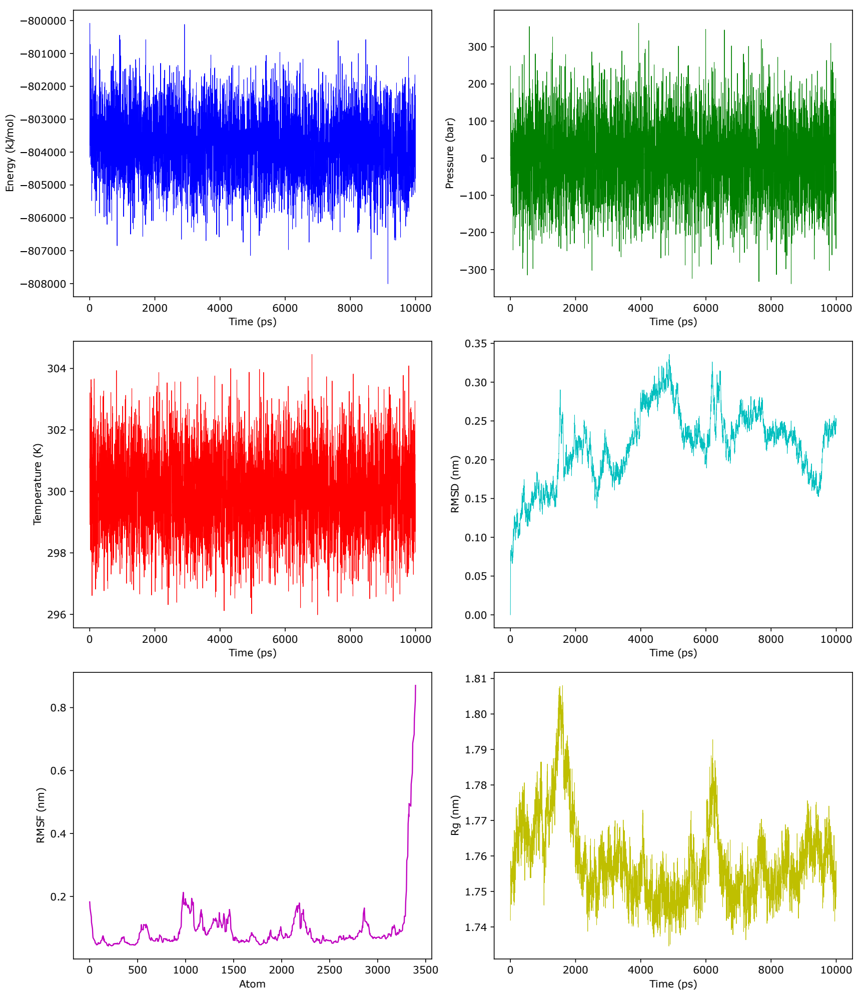


B)

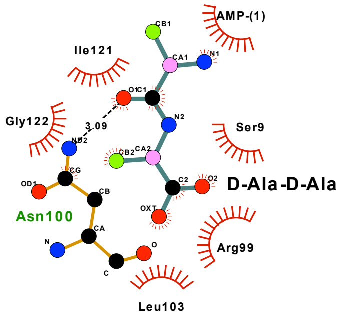


| | |
|---|---|
| Hydrophobic contacts: | 229.308 |
| Van der waals: | -281.003 |
| Deformation effect: | 2 |
| Hydrogen bonds (HB): | 1 |
| Repulsive VDW score: | 9 |
| London dispersion force: | 2.65265 |
| predicted pKD: | 3.63864 |

**Supplementary Figure 11. Structural stability and interactions of the RoyL–AMP–(1)–ᴅ-Ala-ᴅ-Ala complex during MD simulations.** A) Production MD stability metrics over the course of the simulation, as calculated by EquilibraTor. B) Hydrophobic interactions and hydrogen bonds between RoyL and the ᴅ-Ala-ᴅ-Ala substrate, along with the predicted binding affinity.

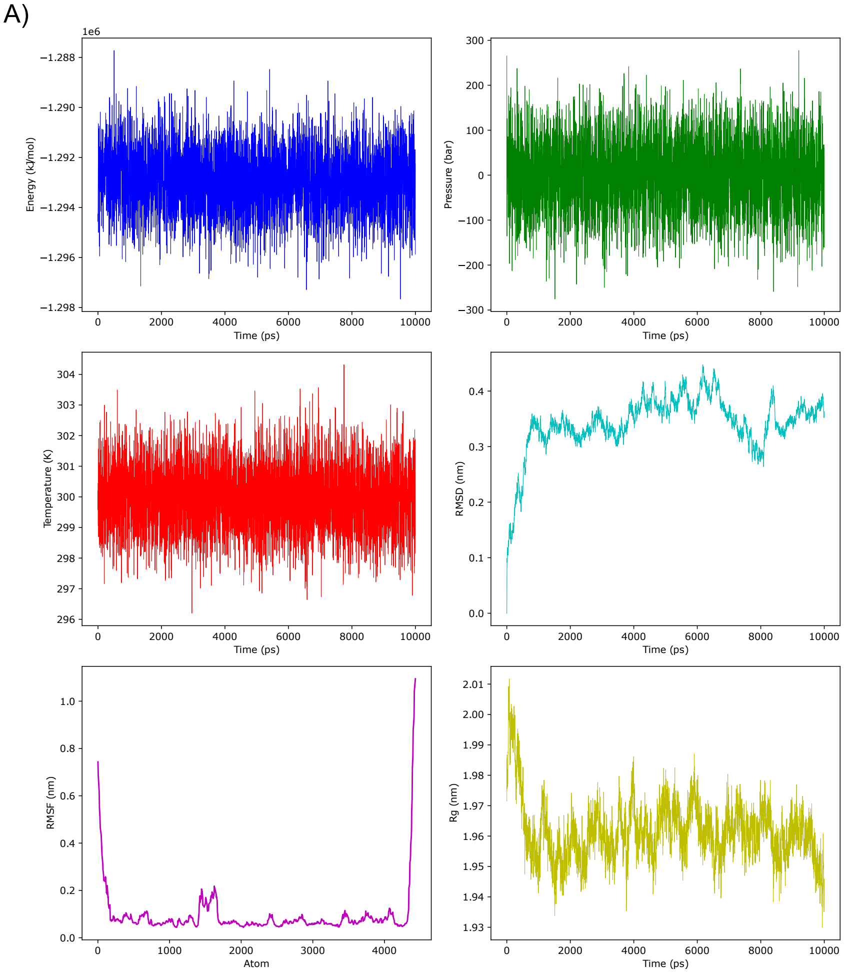


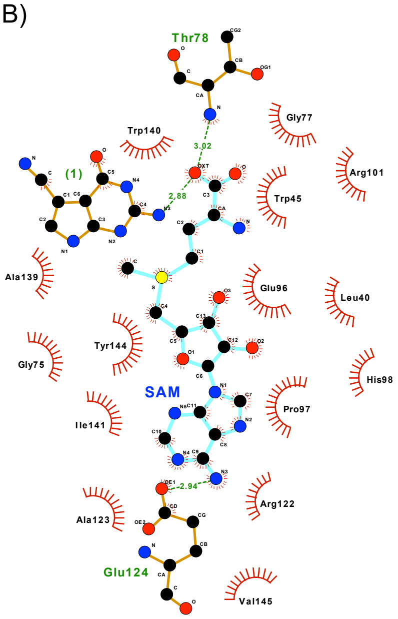


| | |
|---|---|
| Hydrophobic contacts: | 945.248 |
| Van der waals: | -958.49 |
| Deformation effect: | 5.5 |
| Hydrogen bonds (HB): | 3 |
| Repulsive VDW score: | 55 |
| London dispersion force: | 16.071 |
| predicted pKD: | 7.35629 |

**Supplementary Figure 12. Structural stability and interactions of the HuiC–SAM–(1) complex during MD simulations.** A) Production MD stability metrics over the course of the simulation, as calculated by EquilibraTor. B) Hydrophobic interactions and hydrogen bonds between HuiC and the SAM cofactor, along with the predicted binding affinity.

A)

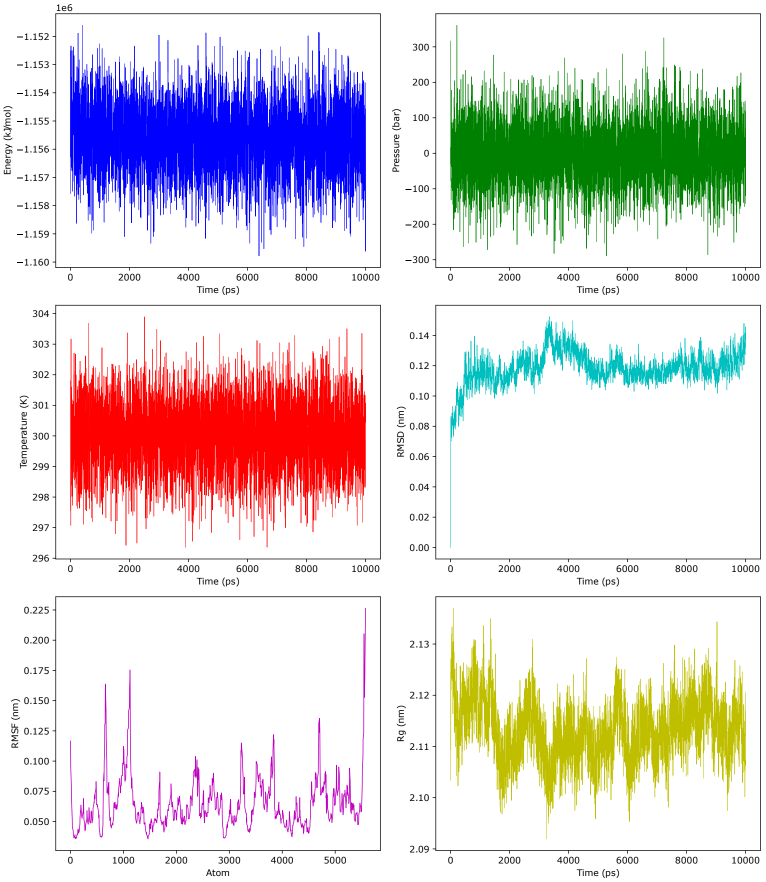


B)

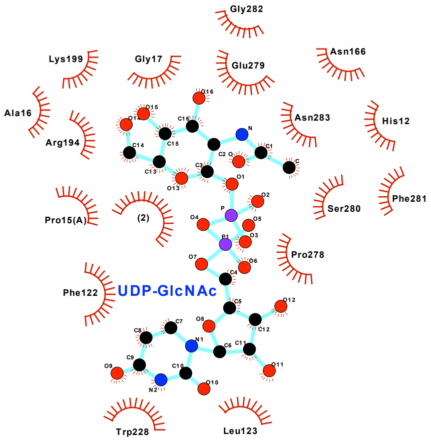


| | |
|---|---|
| Hydrophobic contacts: | 740.754 |
| Van der waals: | -1053.32 |
| Deformation effect: | 9 |
| Hydrogen bonds (HB): | 0 |
| Repulsive VDW score: | 33 |
| London dispersion force: | 10.2202 |
| predicted pKD: | 6.69502 |

**Supplementary Figure 13. Structural stability and interactions of the HuiG–UDP-GlcNAc–(2) complex during MD simulations.** A) Production MD stability metrics over the course of the simulation, as calculated by EquilibraTor. B) Hydrophobic interactions and hydrogen bonds between HuiC and the UDP-GlcNAc substrate, along with the predicted binding affinity.

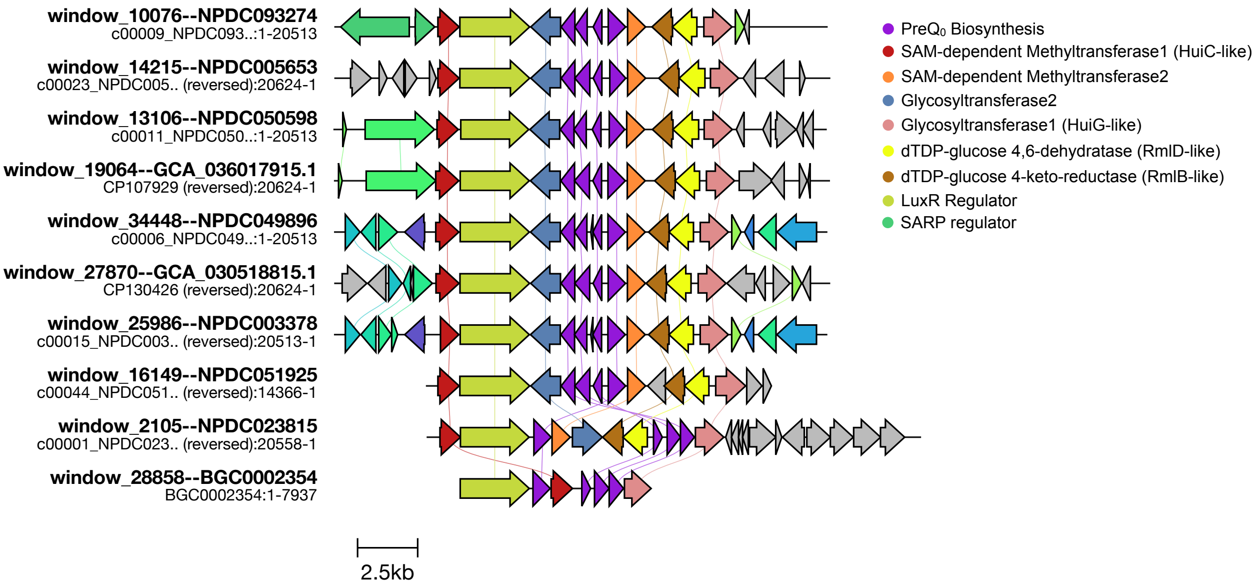


**Supplementary Figure 14.** Genomic neighborhoods of GATOR-GC–predicted members of the dapiramicin A subfamily. Genomic neighborhood comparisons were generated using clinker. The huimycin BGC is shown at the bottom end for comparison.

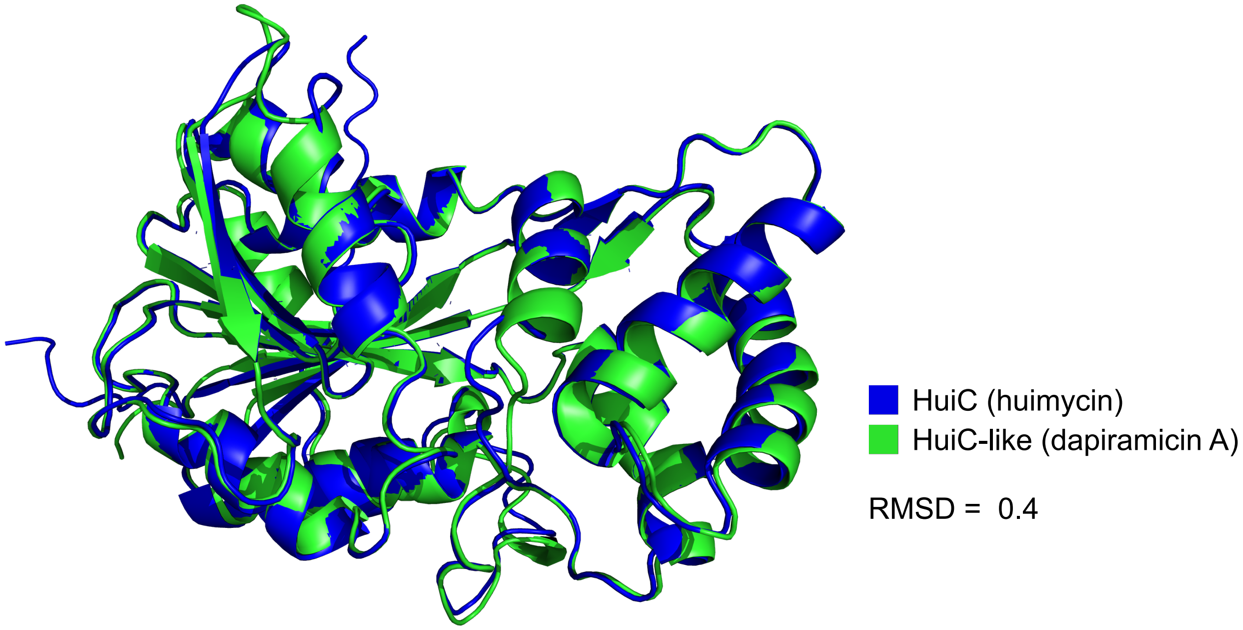


**Supplementary Figure 15. Structural alignment of HuiC homolog models.** Superposition of the reference HuiC structure (blue) and the predicted HuiC homolog (green), showing strong overall structural conservation. Major secondary structure elements, including α-helices and β-sheets, are well aligned, supporting structural similarity between the experimentally characterized enzyme and the predicted homolog.

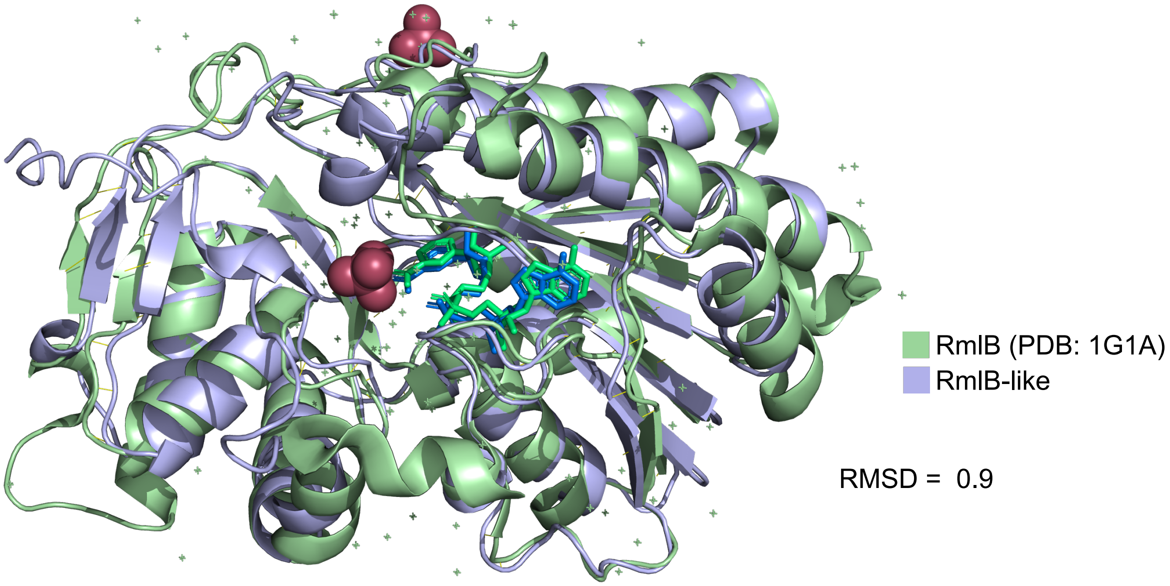


**Supplementary Figure 16. Structural alignment of the RmlB crystal structure from *Salmonella enterica* and a homolog model.** Superposition of the reference RmlB crystal structure (green) and the predicted RmlB-like homolog (purple) shows high structural similarity. The NAD cofactors occupy conserved positions in both structures, and the sulfate ion present in the crystal structure is shown. Major secondary structure elements are well aligned, supporting conservation of the overall fold and cofactor-binding architecture.

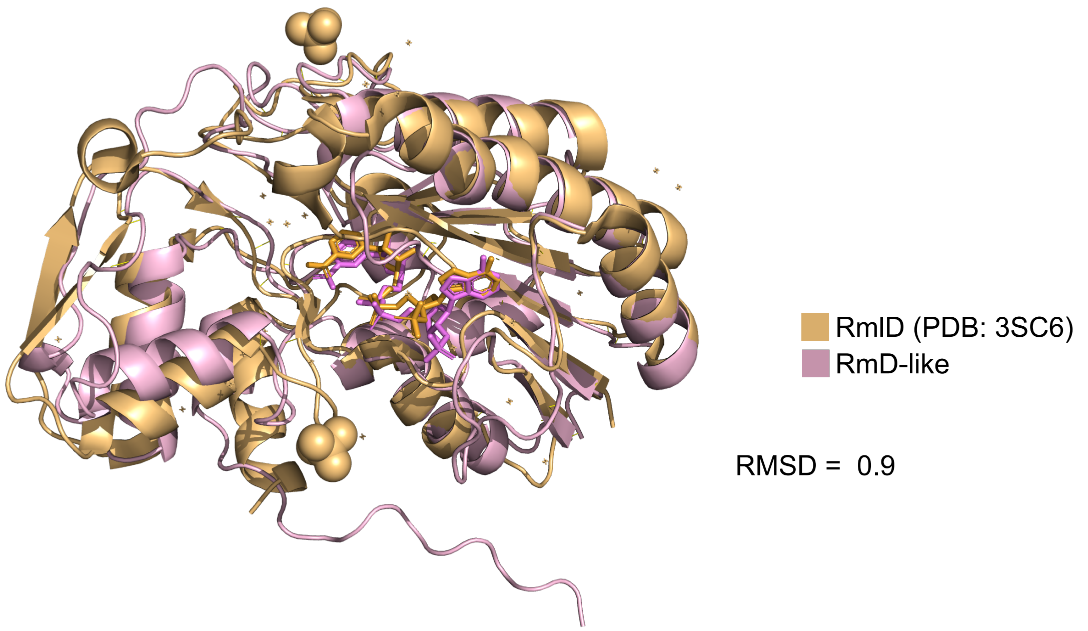


**Supplementary Figure 17. Structural alignment of the RmlD crystal structure from *Bacillus anthracis* and a homolog model.** Superposition of the reference RmlD crystal structure (orange) and the predicted RmlD-like homolog (pink) shows high structural similarity. The NADP cofactors occupy conserved positions in both structures, and the sulfate ion present in the crystal structure is shown. Major secondary structure elements are well aligned, supporting conservation of the overall fold and cofactor-binding architecture.

A)

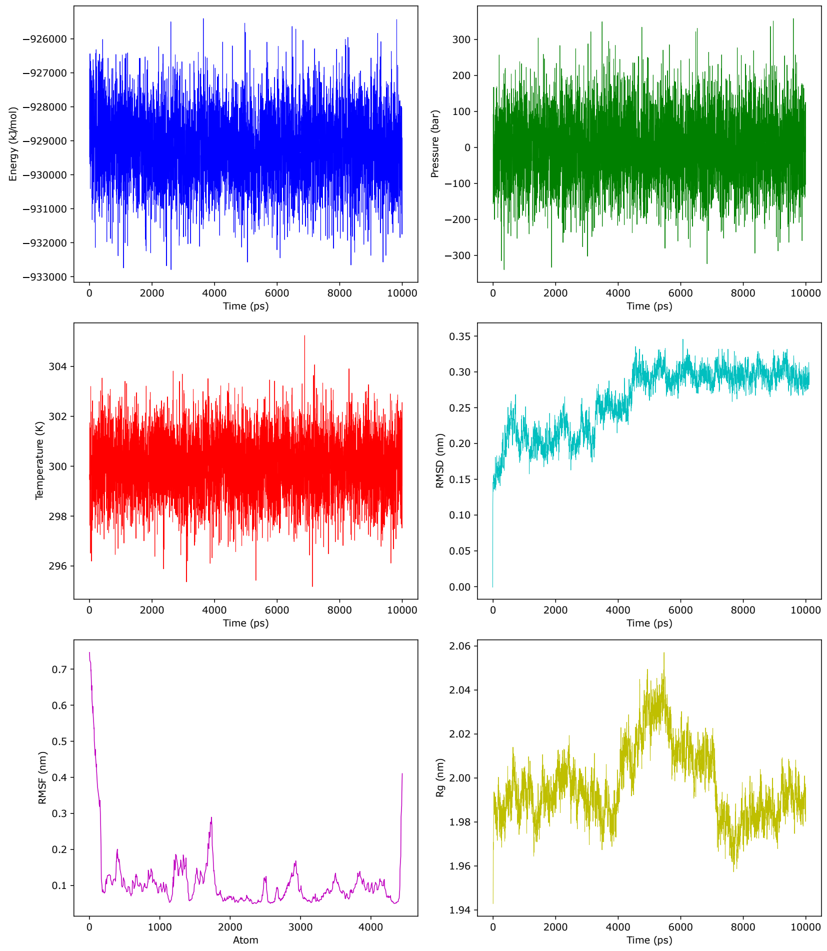


B)

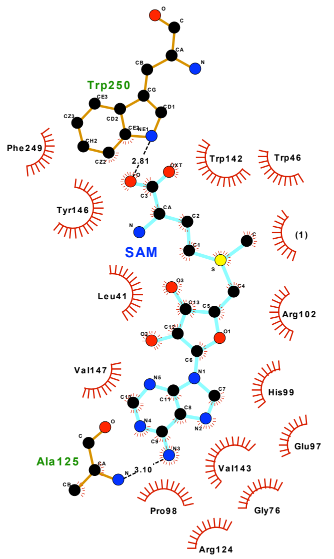


| | |
|---|---|
| Hydrophobic contacts: | 977.227 |
| Van der waals: | -949.924 |
| Deformation effect: | 5.5 |
| Hydrogen bonds (HB): | 5 |
| Repulsive VDW score: | 34 |
| London dispersion force: | 10.063 |
| predicted pKD: | 7.12384 |

**Supplementary Figure 18. Structural stability and interactions of the HuiC-like–SAM–(1) complex during MD simulations.** A) Production MD stability metrics over the course of the simulation, as calculated by EquilibraTor. B) Hydrophobic interactions and hydrogen bonds between HuiC-like and the SAM cofactor, along with the predicted binding affinity.

A)

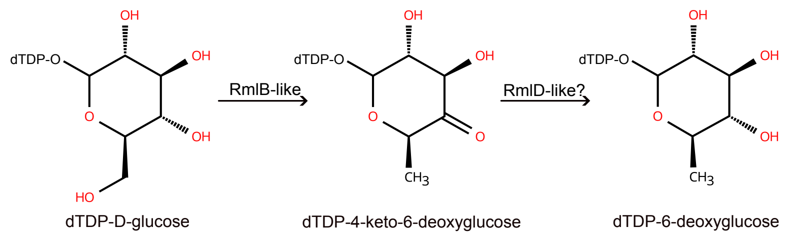


B)

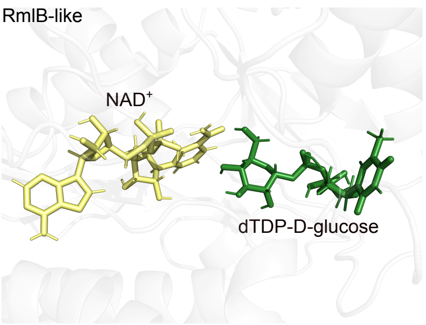


C)

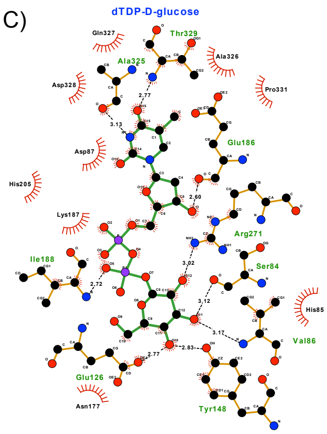


| | |
|---|---|
| Hydrophobic contacts: | 582.268 |
| Van der waals: | -1032.73 |
| Deformation effect: | 10 |
| Hydrogen bonds (HB): | 9 |
| Repulsive VDW score: | 28 |
| London dispersion force: | 8.63175 |
| predicted pKD: | 6.29995 |

D)

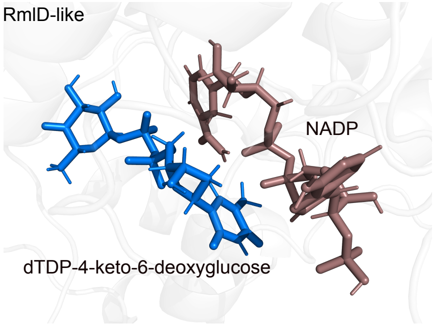


E)

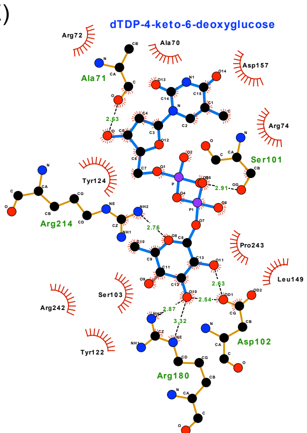


| | |
|---|---|
| Hydrophobic contacts: | 554.689 |
| Van der waals: | -861.494 |
| Deformation effect: | 10 |
| Hydrogen bonds (HB): | 7 |
| Repulsive VDW score: | 38 |
| London dispersion force: | 11.6991 |
| predicted pKD: | 5.94276 |

F)

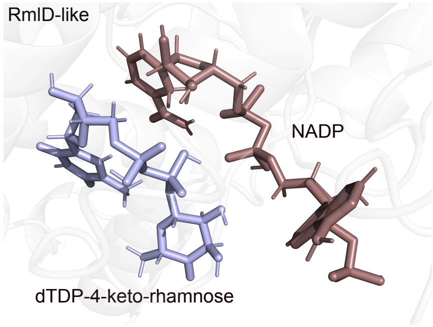


G)

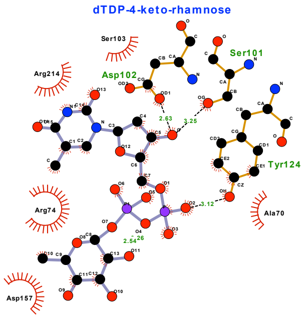


| | |
|---|---|
| Hydrophobic contacts: | 289.947 |
| Van der waals: | -737.64 |
| Deformation effect: | 10 |
| Hydrogen bonds (HB): | 5 |
| Repulsive VDW score: | 19 |
| London dispersion force: | 5.71092 |
| predicted pKD: | 4.35138 |

**Supplementary figure 19. Enzyme-substrate interactions in deoxy-sugar biosynthesis within the candidate dapiramicin A biosynthetic pathway**. (A) Putative biosynthetic steps for the formation of dTDP-6-deoxysugar from dTDP-D-glucose. B)

Representative snapshot from production MD simulations of the RmlB-like in complex with dTDP-6-deoxysugar and $NAD^+$, highlighting a potential pre-reactive state. C) Key hydrophobic and hydrogen-bond interactions stabilizing substrate binding within the active site, along with the predicted binding affinity for dTDP-D-glucose. D) Representative snapshot from production MD simulations of the RmlD-like in complex with dTDP-4-keto-6-deoxysugar and NADP, highlighting a potential pre-reactive state. E) Key hydrophobic and hydrogen-bond interactions stabilizing substrate binding within the active site, along with the predicted binding affinity for dTDP-4-keto-6-deoxysugar. F) Representative snapshot from production MD simulations of the RmlD-like in complex with dTDP-4-keto-rhamnose and NADP, highlighting a potential pre-reactive state. G) Key hydrophobic and hydrogen-bond interactions stabilizing substrate binding within the active site, along with the predicted binding affinity for dTDP-4-keto-rhamnose.

A)

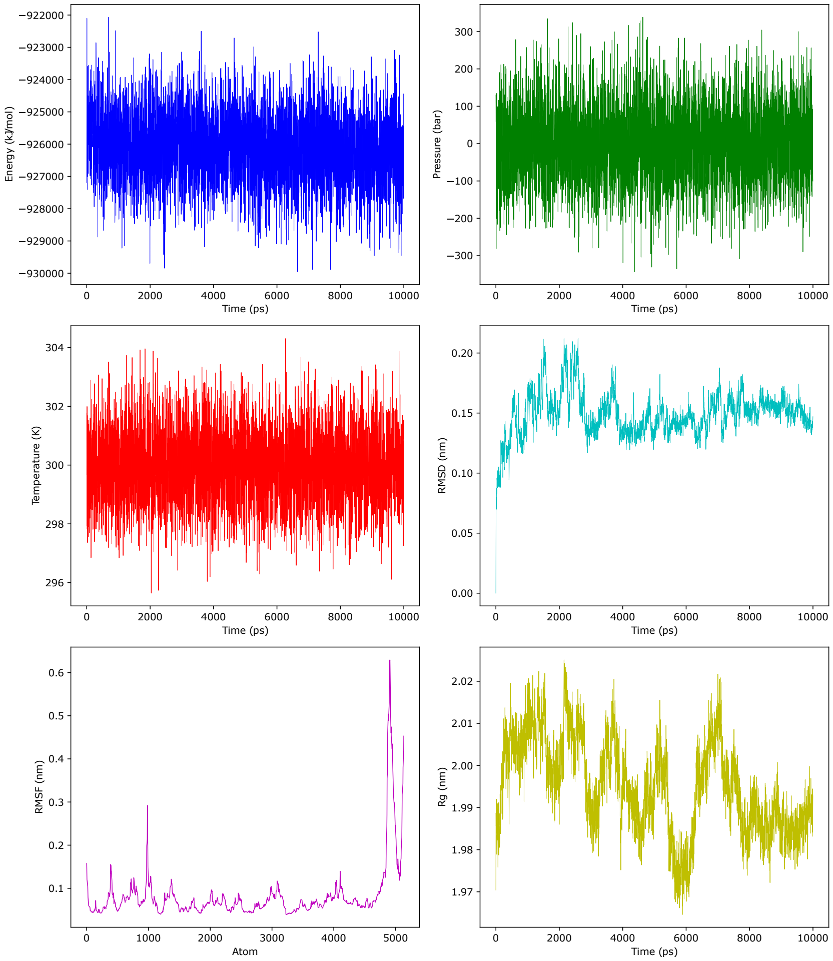


B)

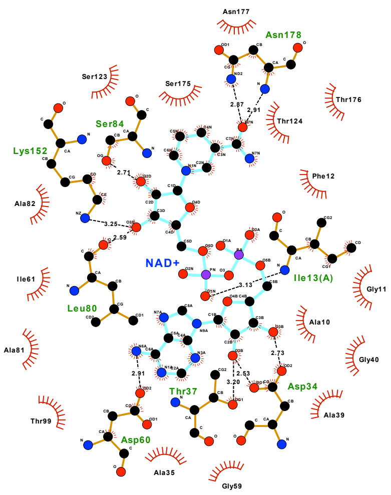


| | |
|---|---|
| Hydrophobic contacts: | 1175.21 |
| Van der waals: | -1474.82 |
| Deformation effect: | 14 |
| Hydrogen bonds (HB): | 11 |
| Repulsive VDW score: | 31 |
| London dispersion force: | 9.41249 |
| predicted pKD: | 8.2034 |

**Supplementary Figure 20. Structural stability and interactions of the RmlB-like-NAD$^{+}$-dTDP-D-glucose complex during MD simulations.** A) Production MD stability metrics over the course of the simulation, as calculated by EquilibraTor. B) Hydrophobic interactions and hydrogen bonds between RmlB-like and the NAD cofactor, along with the predicted binding affinity.

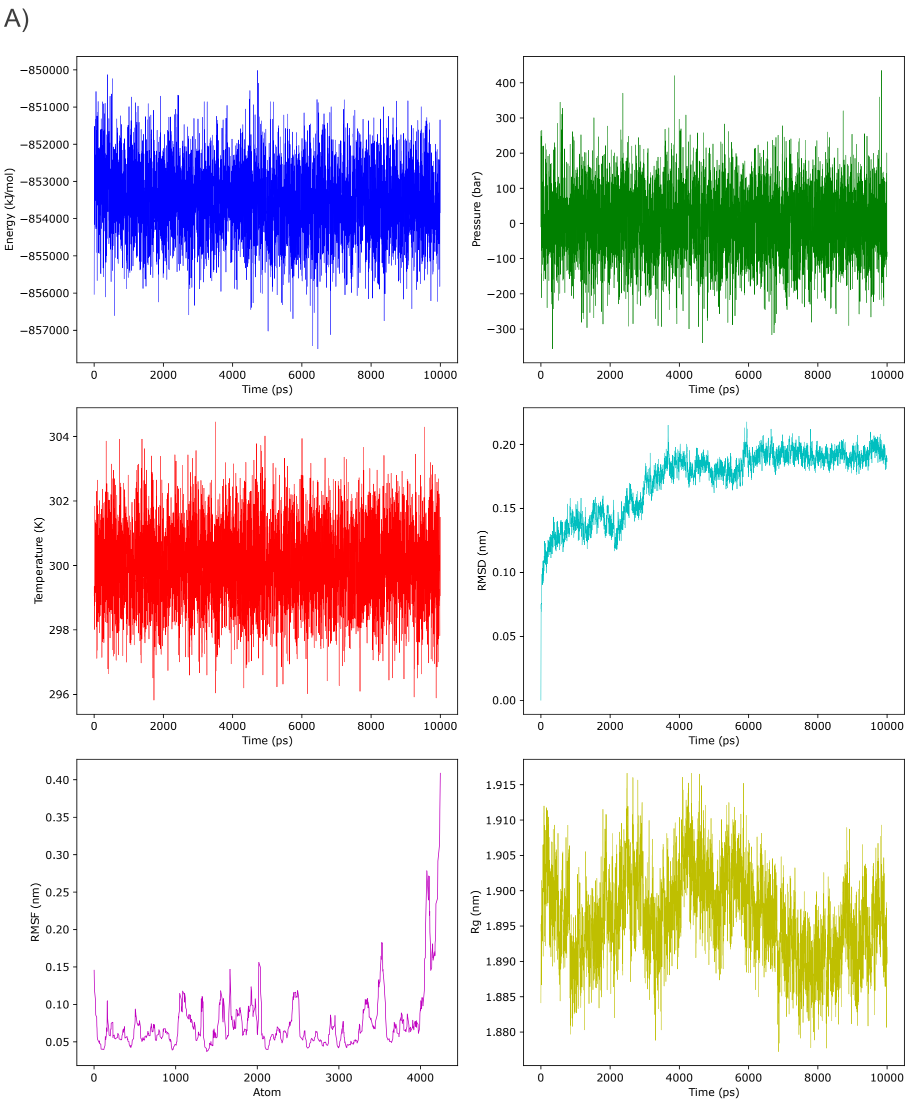


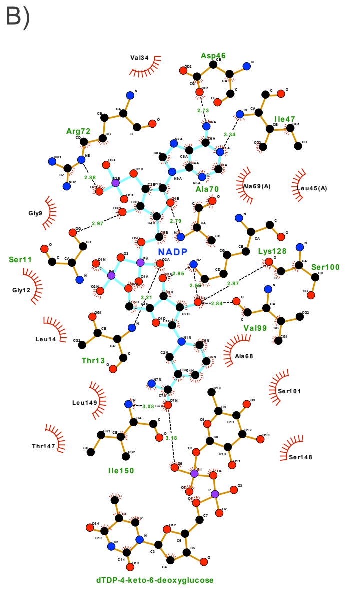


| | |
|---|---|
| Hydrophobic contacts: | 1053.73 |
| Van der waals: | -1291.46 |
| Deformation effect: | 10.5 |
| Hydrogen bonds (HB): | 14 |
| Repulsive VDW score: | 44 |
| London dispersion force: | 13.3944 |
| predicted pKD: | 7.70278 |

**Supplementary Figure 21. Structural stability and interactions of the RmlD-like-NADP-dTDP-4-keto-6-deoxyglucose complex during MD simulations.** A) Production MD stability metrics over the course of the simulation, as calculated by EquilibraTor. B) Hydrophobic interactions and hydrogen bonds between RmlD-like and the NADP cofactor, along with the predicted binding affinity.

A)

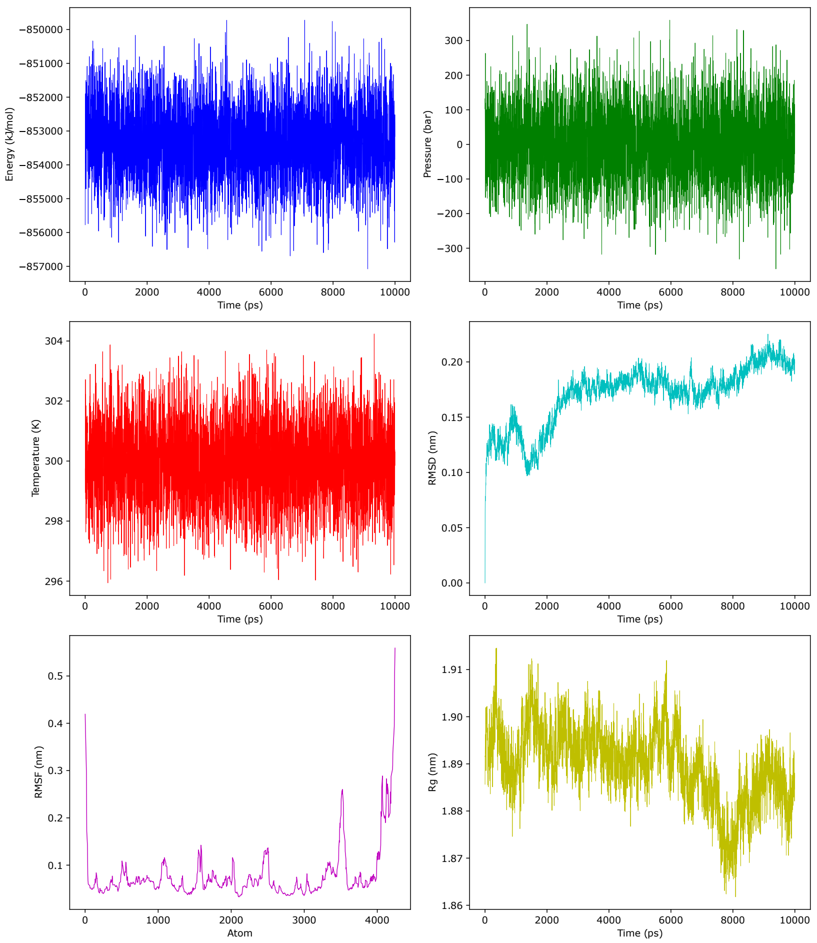


B)

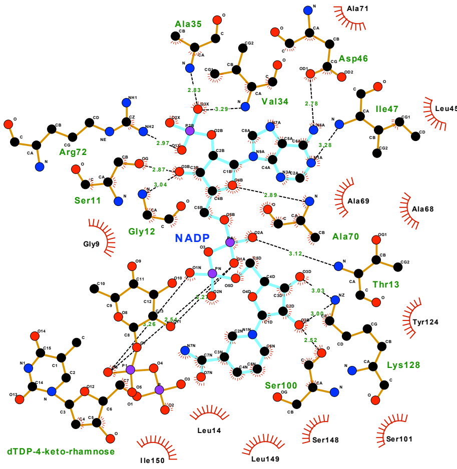


| | |
|---|---|
| Hydrophobic contacts: | 972.315 |
| Van der waals: | -1319.01 |
| Deformation effect: | 11 |
| Hydrogen bonds (HB): | 16 |
| Repulsive VDW score: | 50 |
| London dispersion force: | 15.2985 |
| predicted pKD: | 7.79847 |

**Supplementary Figure 22. Structural stability and interactions of the RmlD-like-NADP-dTDP-4-rhamnose complex during MD simulations.** A) Production MD stability metrics over the course of the simulation, as calculated by EquilibraTor. B) Hydrophobic interactions and hydrogen bonds between RmlD-like and the NADP cofactor, along with the predicted binding affinity.

**Supplementary Table S1. GATOR-GC-predicted QueE gene clusters.** This table lists the genomic windows identified by GATOR-GC that contain QueE hits. For each window, the table reports the window identifier, taxonomy classification according to GTDB-Tk, the presence or absence of genes involved in 7-deazapurine biosynthesis, and the presence or absence of enzymes responsible for the incorporation of 7-deazapurine derivatives into nucleic acids.

SupplementaryTableS1.xls
